# Defining a leptomeningeal blood–cerebrospinal fluid barrier as a specialized vascular interface

**DOI:** 10.64898/2026.08.19.745529

**Authors:** Philip V. Seegren

**Author notes:** Corresponding Author: Philip Seegren.

## Abstract

Central nervous system vascular barriers comprise anatomically distinct interfaces that regulate molecular exchange and immune communication between the circulation and neural tissues. Although the blood brain barrier has been extensively characterized, whether endothelial cells within the leptomeningeal vasculature represent a specialized vascular population distinct from cortical blood brain barrier endothelial cells has remained unclear. Here, we integrate cross study transcriptomic analyses, single nucleus RNA sequencing, and experimental models of neonatal meningitis to define the molecular and functional organization of leptomeningeal endothelial cells. We show that leptomeningeal endothelial cells possess a transcriptional program distinct from cortical blood brain barrier endothelial cells, characterized by enhanced extracellular matrix remodeling and immune interface programs together with reduced expression of canonical Wnt/β catenin signaling transcripts. These molecular differences coincide with a transcriptionally distinct stromal Wnt ligand environment, vascular architecture, and context-dependent remodeling during infection. Together, our findings define the leptomeningeal blood cerebrospinal fluid barrier as a specialized CNS vascular interface with distinct molecular, structural, and functional properties, expanding the current framework of CNS barrier organization.

## Introduction

The central nervous system relies on multiple vascular and epithelial barrier interfaces to regulate molecular exchange, immune surveillance, and tissue homeostasis. These include the blood–brain barrier (BBB)^1^, the choroid plexus blood–cerebrospinal fluid barrier (BCSFB)^2^, circumventricular organs, the blood–retinal barrier^3^, and the meninges^4^. Although these interfaces collectively preserve the neural environment, they differ in anatomical organization, permeability, and physiological function. These differences arise in part from local microenvironmental cues that instruct endothelial identity, a principle increasingly recognized across vascular beds throughout the body^5^. Molecular and spatial profiling has established that endothelial and perivascular cells adopt region-specific transcriptional programs across the cerebrovascular tree^6^, circumventricular organs^7^, choroid plexus^8^, and retina^9^, establishing regional vascular specialization as a fundamental principle of CNS barrier biology. However, whether this principle extends to the leptomeningeal vasculature remains unclear.

The leptomeninges comprise the arachnoid barrier, the cerebrospinal fluid-filled subarachnoid space, and the pia mater, forming the tissue that envelops the brain and spinal cord. Unlike cortical BBB vessels embedded within the neural parenchyma, leptomeningeal vessels reside within a fibroblast-rich stromal environment adjacent to cerebrospinal fluid and serve as a major site of immune surveillance, immune cell trafficking, and fluid exchange^10,11^. The leptomeningeal stroma is organized into anatomically and molecularly distinct compartments composed of specialized pial, arachnoid barrier, and dural border fibroblast populations together with mural, and immune cells^6,12,13^. However, although recent single-cell studies have cataloged leptomeningeal endothelial cells, their molecular identity remains poorly defined, with limited direct comparison to cortical BBB endothelium and no framework for defining a specialized leptomeningeal vascular barrier. More recently, developmental analyses demonstrated that these layer-specific fibroblast populations establish the structural organization of the leptomeninges, providing a developmental framework for this highly specialized border tissue^14^. Complementing these findings, specialized anatomical conduits linking the subarachnoid space and dura further establish the leptomeninges as a dynamic neuroimmune interface rather than a passive protective covering of the brain^15^. Together, these studies establish the leptomeninges as a highly organized barrier tissue with specialized stromal, immune, and developmental programs. However, it remains unknown whether the endothelial cells that vascularize this compartment exhibit comparable specialization. As a result, the existence of a distinct leptomeningeal vascular barrier has not been established.

Endothelial specialization is a defining feature of vascular barrier biology and arises through continuous communication between endothelial cells and their surrounding tissue microenvironment. Within the cortical neurovascular unit^16^, endothelial cells are integrated with pericytes^17^, astrocytic end feet^18^, neurons, and extracellular matrix components that collectively regula te vascular stability, barrier integrity^19^, leukocyte trafficking^20^, and molecular transport^1,21^. These interactions establish molecular programs that distinguish CNS endothelial cells from peripheral endothelium and are essential for BBB development and maintenance. Among the signaling pathways that instruct CNS endothelial identity, canonical Wnt/β-catenin signaling has emerged as the central developmental pathway governing BBB specification^22^. Within the cortical neurovascular unit, neural-derived Wnt7a and Wnt7b, together with Norrin signaling in the retina, activate endothelial β-catenin signaling through GPR124, RECK, LRP5/6, and TSPAN12 to induce CNS-specific endothelial gene programs, suppress vascular permeability, and maintain blood–brain and blood–retinal barrier integrity^22–26^. Current models of CNS endothelial specification are largely based on the cortical neurovascular unit, where neural-derived Wnt and Norrin ligands instruct endothelial barrier identity. Whether comparable ligand environments exist within the leptomeninges, where endothelial cells reside outside the neural parenchyma and are instead surrounded by fibroblasts and cerebrospinal fluid, remains unknown. Likewise, it is unclear whether endothelial cells across distinct CNS vascular interfaces share a common repertoire of Wnt receptors and co-receptors or whether they possess region-specific signaling programs adapted to their local microenvironment.

Growing evidence indicates that cerebrovascular dysfunction is a central feature of neurological disease, with BBB impairment implicated in Alzheimer’s disease, Parkinson’s disease, ALS, multiple sclerosis, stroke, traumatic brain injury, and neuroinflammatory disorders^27–29^. Human vascular single-cell atlases have further demonstrated that neurological disease is accompanied by cell-type-specific remodeling of endothelial transcriptional programs, linking vascular cell states to genetic risk and disease-associated signaling pathways rather than uniform barrier failure^30^. Together, these studies establish endothelial identity as a key determinant of CNS vascular function in both health and disease. However, nearly all current models of CNS endothelial dysfunction are derived from the cortical neurovascular unit, leaving it unknown whether analogous molecular programs exist within the leptomeningeal vasculature or whether this interface exhibits distinct responses to physiological and pathological challenge.

Here, we combined cross-study transcriptomic meta-analysis, single-nucleus RNA sequencing, whole-mount leptomeninges microscopy, and an experimental model of neonatal bacterial meningitis to determine whether endothelial cells associated with the leptomeninges represent a specialized CNS vascular interface. We first defined a conserved brain endothelial transcriptional signature and identified reproducible endothelial heterogeneity that persisted beyond technical, cellular, and canonical endothelial subtype variation, raising the possibility that anatomical origin contributes to endothelial specialization. We then demonstrate that leptomeningeal endothelial cells constitute a transcriptionally distinct endothelial population characterized by extracellular matrix remodeling, adhesion-associated, and immune-interface programs that distinguish them from cortical BBB endothelial cells. We further identify a unique leptomeningeal Wnt signaling environment and show that leptomeningeal endothelial cells express a distinct repertoire of canonical BBB-associated Wnt signaling components. Finally, we define the structural organization of the leptomeningeal vasculature and demonstrate that it exhibits compartment-specific inflammatory activation, vascular permeability, and basement membrane remodeling during neonatal bacterial meningitis. Together, these findings establish the leptomeningeal blood–cerebrospinal fluid barrier (LBCSFB) as a specialized CNS vascular interface with distinct molecular, structural, and functional properties.

## Results

### A cross-study endothelial atlas defines a reproducible brain endothelial transcriptional identity

To establish the broader context for endothelial specialization, we assembled a cross-study transcriptomic atlas of endothelial and CNS-associated cell populations from publicly available bulk RNA-seq datasets. Following manual curation and harmonization of sample metadata, the atlas comprised 316 samples from 49 independent studies, including brain endothelial cells, peripheral endothelial populations, cultured brain microvascular endothelial cells (BMECs), microglia, pericytes, and whole-brain tissue **(Fig. 1A; Extended Data Fig. 1A–D)**. Principal component analysis of the integrated dataset separated major cell and tissue classes, and endothelial-focused PCA distinguished brain endothelial cells from peripheral endothelial populations **(Fig. 1B–C)**. Marker analysis confirmed enrichment of canonical endothelial genes across endothelial samples and appropriate lineage-specific marker expression in non-endothelial populations, supporting the integrity of the curated atlas **(Extended Data Fig. 1E–F)**.

**Fig. 1:**
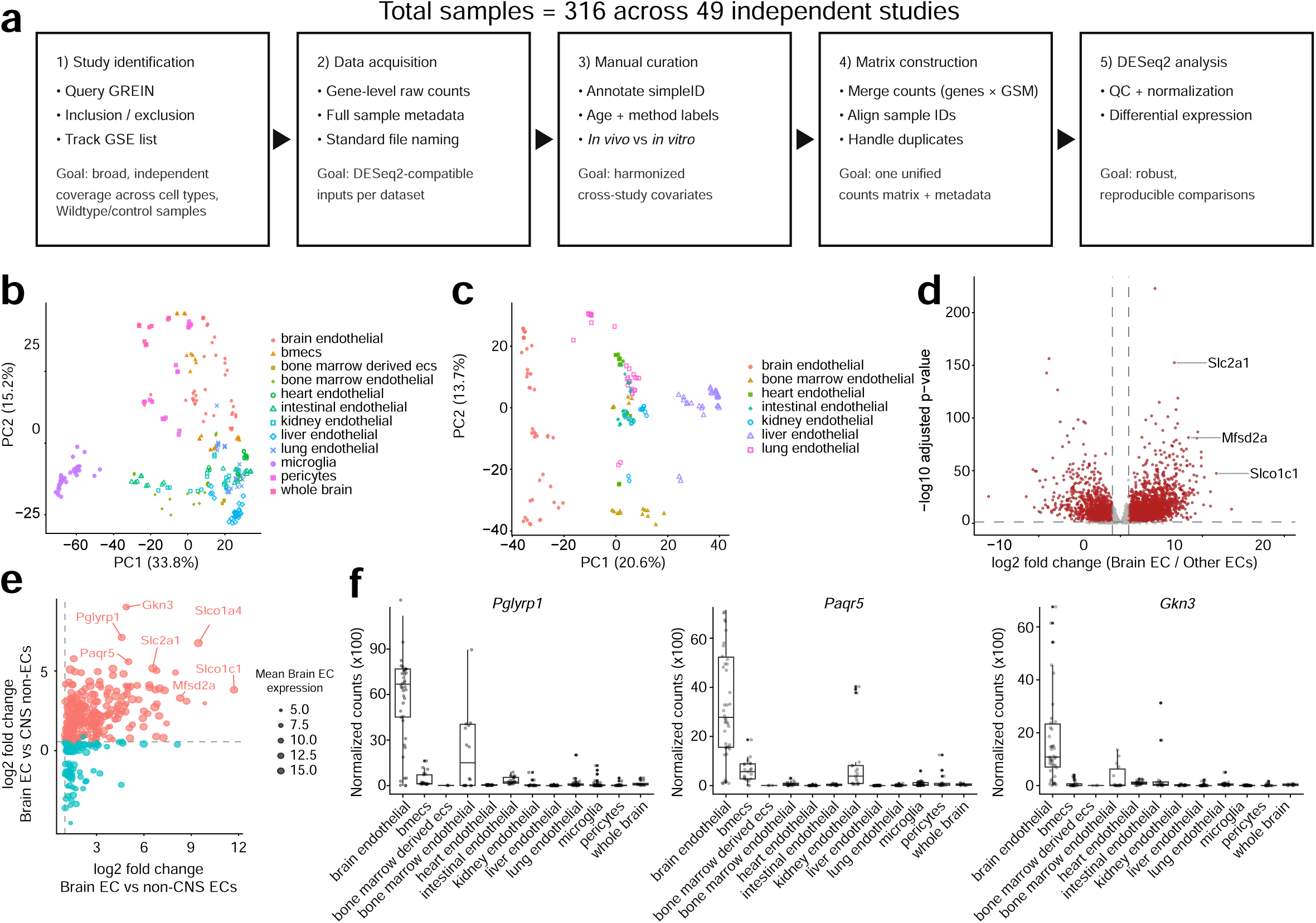
Cross-study meta-analysis defines reproducible Brain EC identity. (a) Schematic overview of the cross-study RNA-seq atlas construction workflow. Publicly available datasets were identified through GREIN, manually curated, harmonized into a unified gene-by-sample count matrix, and analyzed using DESeq2. (b) Principal component analysis (PCA) of all curated samples following variance-stabilizing transformation. Samples are colored by manually assigned cell or tissue identity. (c) PCA restricted to endothelial populations. Brain endothelial cell (Brain EC) samples separate from peripheral endothelial populations, indicating a conserved Brain EC transcriptional identity across independent studies. (d) Differential expression analysis comparing Brain ECs with all other endothelial populations. Each point represents a gene plotted by shrunken log2 fold change and −log10 adjusted p-value. Canonical BBB-associated genes, including Slc2a1, Mfsd2a, and Slco1c1, are highlighted. (e) Candidate Brain EC marker prioritization based on enrichment relative to non-CNS endothelial populations and CNS non-endothelial populations. Point color indicates enrichment category and point size indicates mean Brain EC expression. Genes enriched relative to both comparison groups are highlighted, with selected candidates labeled. (f) Normalized count distributions for representative Brain EC-enriched candidate genes (*Pglyrp1*, *Paqr5*, and *Gkn3*) across endothelial and CNS-associated sample groups. Boxes indicate the interquartile range with the median shown as the center line; whiskers extend to 1.5× the interquartile range and individual points represent individual RNA-seq samples.

Differential expression analysis comparing brain endothelial cells with other endothelial populations identified strong enrichment of canonical blood–brain barrier (BBB)-associated genes, including Slc2a1, Mfsd2a, and Slco1c1 **(Fig. 1D)**. Examination of the most enriched genes across all sample groups demonstrated that each cellular compartment was characterized by expected lineage-associated transcriptional signatures, with BBB transporters and CNS endothelial markers selectively enriched within brain endothelial samples **(Extended Data Fig. 2)**. These findings validated the ability of the integrated atlas to recover known brain endothelial transcriptional programs.

To identify a robust and reproducible brain endothelial signature, candidate genes were filtered using four complementary criteria: enrichment relative to non-CNS endothelial populations, high expression across independent brain endothelial studies, expression above the mean of other endothelial samples, and enrichment relative to CNS non-endothelial populations **(Extended Data Fig. 3A–B)**. Integration of these metrics prioritized genes that consistently distinguished brain endothelial cells across datasets while minimizing study-specific effects **(Fig. 1E)**. Visualization of representative candidates confirmed selective expression within brain endothelial samples **(Fig. 1F)**. The intersection of these criteria identified a conserved brain endothelial transcriptional signature that included canonical BBB transporters and signaling molecules such as Mfsd2a, Slc2a1, Slc7a5, Slco1c1, Slco1a4, Lef1, and Zic3, while also highlighting less-characterized candidates including Pglyrp1, Paqr5, and Gkn3 that exhibited robust and reproducible enrichment across independent brain endothelial datasets **(Fig. 1F; Extended Data Fig. 3C)**. Together, these analyses define a conserved brain endothelial identity that is distinct from both peripheral endothelial and CNS non-endothelial populations.

### Cross-study meta-analysis reveals residual brain endothelial heterogeneity beyond non-endothelial and endothelial subtype variation

Although brain endothelial samples shared a conserved BBB-associated transcriptional identity, principal component analysis (PCA) of brain endothelial samples alone revealed substantial study-associated variation (Fig. 2A). Despite this heterogeneity, expression of canonical BBB transporters and Wnt-responsive genes—including *Slco1c1, Mfsd2a, Slc2a1, Cldn5, Abcb1a, Abcg2, Lrp5, Lrp6, Reck, Tspan12, Ctnnb1, Axin2, Lef1, Tcf7, Apcdd1, Nkd1, Notum,* and *Spock2*—remained broadly preserved across independent studies **(Extended Data Fig. 4A)**. Brain endothelial PCA colored by age and isolation method demonstrated that study-to-study variation was not explained by a single experimental variable alone **(Extended Data Fig. 4B)**.

**Fig. 2:**
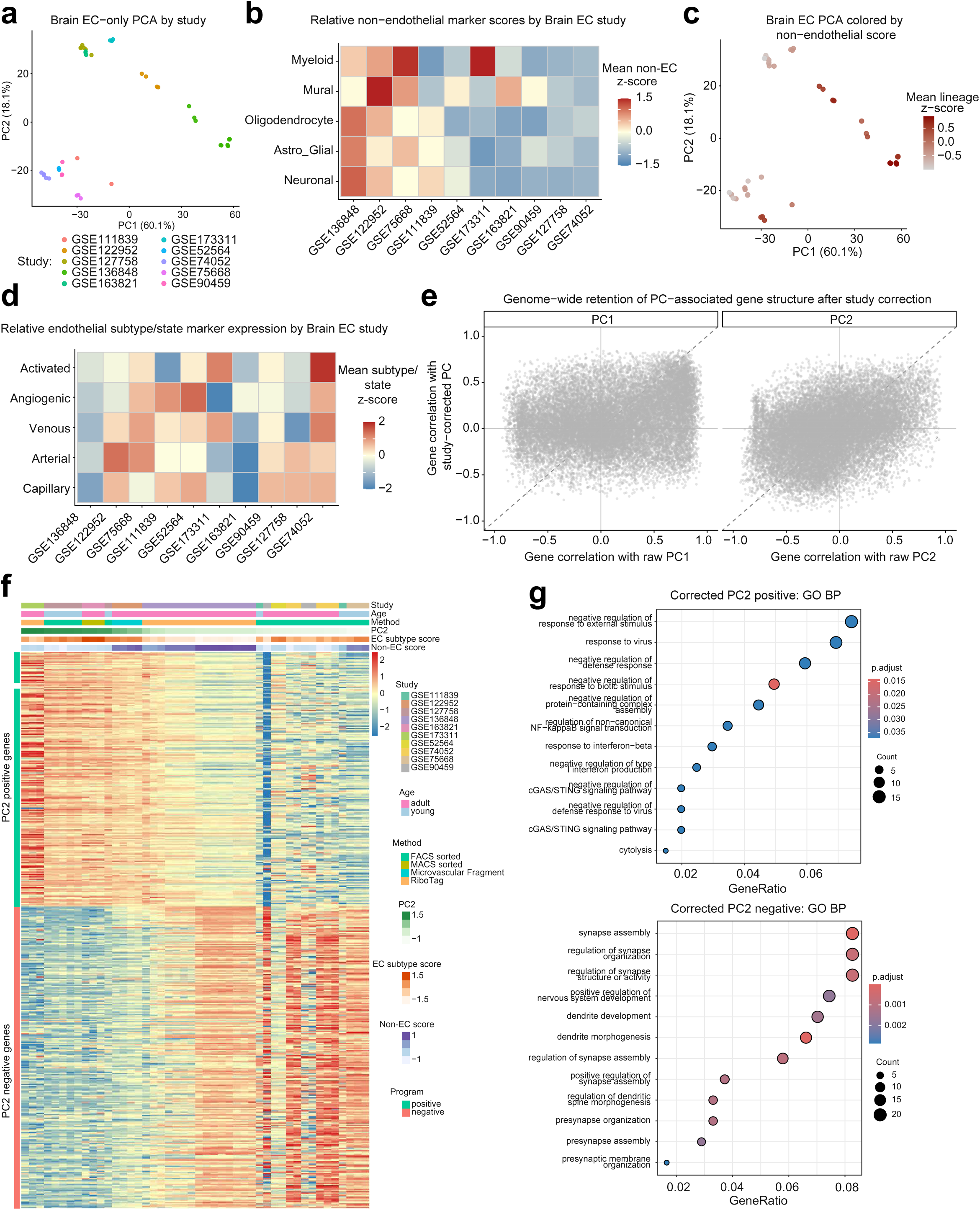
Cross-study meta-analysis reveals residual Brain EC heterogeneity. (a) Principal component analysis (PCA) of Brain EC-only RNA-seq samples colored by GEO study identity. Brain EC samples show substantial study-associated structure despite sharing a conserved endothelial identity. (b) Heatmap showing relative non-endothelial lineage marker scores across independent Brain EC studies. Mean lineage scores were calculated for neuronal, astroglial, oligodendrocyte, mural, and myeloid marker programs and displayed as study-level z-scores. (c) Brain EC PCA colored by total non-endothelial lineage score. Non-endothelial lineage signal contributes to variation among Brain EC studies, particularly along PC1. (d) Heatmap showing relative endothelial subtype/state marker expression across Brain EC studies. Mean program scores are shown for capillary, arterial, venous, angiogenic, and activated endothelial signatures. (e) Genome-wide retention of PC-associated gene structure after study correction. Each point represents a single gene. The x-axis shows the Spearman correlation between gene expression and the corresponding raw principal component (PC1 or PC2), whereas the y-axis shows the correlation between the same gene and the corresponding study-corrected principal component. Dashed diagonal lines indicate perfect agreement between raw and study-corrected gene correlations. The close correspondence of gene correlations before and after study correction indicates that the gene programs defining each principal component are largely preserved despite removal of study-associated variation. (f) Heatmap of genes retaining significant PC2 association following study correction. Genes are ordered by retained PC2 association and samples are ordered by raw PC2. Annotations indicate study, age, isolation method, PC2 score, endothelial subtype/state score, and non-endothelial score. Rows represent individual genes grouped into two transcriptional programs according to the sign of their correlation with PC2: the positive program contains genes positively correlated with PC2 scores, whereas the negative program contains genes negatively correlated with PC2 scores (g) Gene Ontology Biological Process enrichment of corrected PC2-associated genes. Corrected PC2-positive genes are enriched for antiviral, interferon-associated, cGAS/STING, and defense-response pathways, whereas corrected PC2-negative genes are enriched for synapse assembly, dendrite development, and nervous system organization pathways. GeneRatio denotes the proportion of genes from the corresponding PC2-associated gene program that are annotated to a given GO term (number of overlapping genes divided by the total number of genes submitted for enrichment analysis). Point size indicates the number of genes contributing to the GO term, and point color indicates the Benjamini– Hochberg adjusted *P* value.

Because non-endothelial transcript admixture could contribute to apparent heterogeneity, we quantified non-endothelial marker programs associated with neuronal, astroglial, oligodendrocyte, mural, and myeloid populations across brain endothelial studies **(Fig. 2B)**. Brain endothelial samples retained substantially higher expression of endothelial markers than non-endothelial markers, although the degree of non-endothelial signal varied between studies **(Extended Data Fig. 5A)**. Total non-endothelial score was strongly associated with PC1 but showed little relationship with PC2, indicating that non-endothelial-associated signal contributes substantially to variation along PC1 but has little association with PC2, suggesting that additional sources of variation contribute to the residual transcriptional structure captured by PC2 (**Fig. 2C; Extended Data Fig. 5B–C)**. We next quantified endothelial subtype and state programs, including capillary, arterial, venous, angiogenic, and activated signatures. These programs varied across studies **(Fig. 2D; Extended Data Fig. 5D)**, and collectively explained a portion of the observed variation, but neither non-endothelial nor endothelial subtype/state programs appeared sufficient to explain the transcriptional structure associated with PC2 **(Extended Data Fig. 5E–G)**.

**T**o determine whether this residual transcriptional structure persisted beyond study-specific effects, we corrected gene expression values for study identity and recalculated principal component coordinates from the corrected data. We then compared, for every gene in the dataset, its Spearman correlation (ρ) with the raw principal components to its correlation with the corresponding study-corrected principal components. If study-associated technical variation were the primary driver of the observed PCA structure, genes would be expected to lose their associations with the principal components after correction. Although study correction substantially reduced study-associated variance in PCA space **(Extended Data Fig. 6A–B),** many genes retained both the direction and magnitude of their correlations with PC1 and PC2 **(Extended Data Fig. 6C).** Consistent with this observation, genome-wide gene correlations before and after study correction remained strongly related for both principal components **(Fig. 2E).** Genes defining the positive and negative extremes of both PC1 and PC2 largely preserved their associations after correction, indicating that study correction altered sample positioning in PCA space while preserving much of the underlying gene-level transcriptional organization.

We therefore focused on genes that remained correlated with PC2 following correction for study identity, non-endothelial score, and endothelial subtype/state score. Ranking genes according to their retained correlation with the fully corrected PC2 revealed a continuous transcriptional axis spanning independent brain endothelial datasets **(Extended Data Fig. 7A–B).** Visualization of these retained genes demonstrated coordinated opposing transcriptional programs that persisted after correction for study identity, non-endothelial score, and endothelial subtype/state score **(Fig. 2F; Extended Data Fig. 6D).** Gene Ontology analysis revealed that genes positively correlated with the corrected PC2 were enriched for antiviral defense, interferon signaling, cGAS/STING signaling, and innate immune regulatory pathways, whereas genes negatively correlated with the corrected PC2 were enriched for synapse assembly, dendrite development, and nervous system organization programs **(Fig. 2G).** Analysis of the uncorrected PC2 gene sets revealed broader pathway enrichments prior to correction, including ribosome biogenesis and vesicular transport programs on the positive end of the axis and developmental and sensory-system pathways on the negative end **(Extended Data Fig. 7C–D)**.

Together, these analyses indicate that brain endothelial cells maintain a conserved BBB transcriptional core while also exhibiting reproducible transcriptional heterogeneity across independent studies. This residual variation was only partially accounted for by study identity, non-endothelial signal, isolation method, age, and canonical endothelial subtype/state composition. Instead, the remaining transcriptional variation was organized along a reproducible axis in which genes positively correlated with the corrected PC2 were enriched for innate immune and antiviral pathways, whereas genes negatively correlated with the corrected PC2 were enriched for neurovascular interaction and synaptic organization programs. The persistence of unexplained endothelial heterogeneity across independent datasets raised the possibility that anatomical origin contributes to endothelial specialization. We therefore next asked whether endothelial cells associated with anatomically distinct CNS vascular interfaces occupy different positions along this continuum.

### Leptomeningeal endothelial cells exhibit a transcriptional state distinct from cortical BBB endothelial cells

To determine whether endothelial cells associated with the leptomeningeal blood–cerebrospinal fluid barrier (LBCSFB) represent a molecularly distinct vascular population, we integrated endothelial nuclei derived from cortical and leptomeningeal tissues and compared their transcriptional profiles **(Fig. 3A–B)**. Although cortical and leptomeningeal endothelial cells occupied a shared endothelial manifold, nuclei segregated according to anatomical origin, consistent with origin-associated endothelial transcriptional states rather than a completely homogeneous endothelial population.

**Fig. 3:**
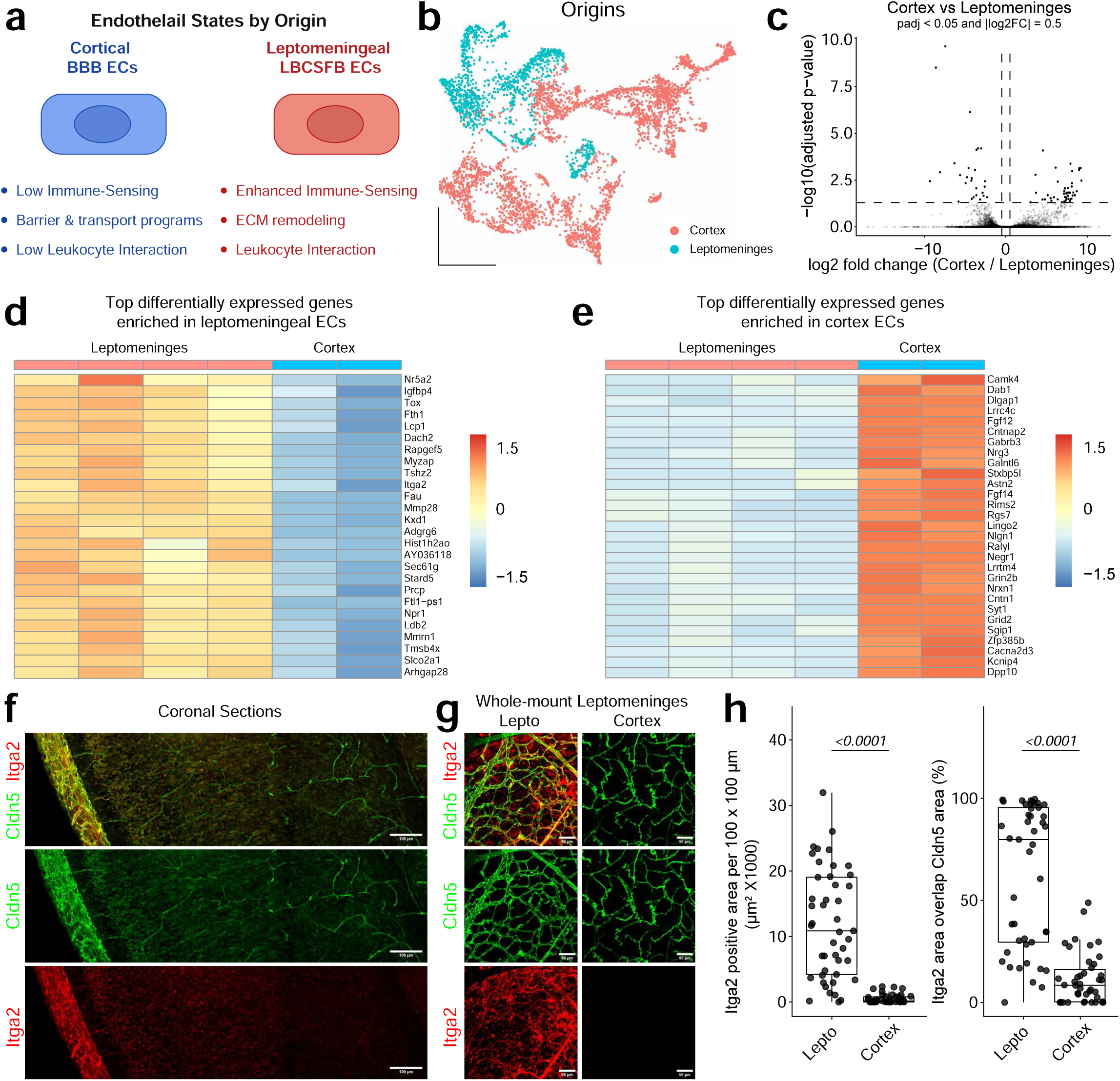
Endothelial cells of the leptomeningeal blood–CSF barrier exhibit a transcriptional state distinct from cortical BBB endothelial cells. (a) Conceptual schematic illustrating major biological distinctions between cortical BBB endothelial cells and leptomeningeal blood–CSF barrier (LBCSFB) endothelial cells. Cortical endothelial cells are characterized by barrier and transport programs and reduced immune interaction, whereas leptomeningeal endothelial cells exhibit enhanced immune sensing, extracellular matrix remodeling, and leukocyte interaction programs. (b) UMAP visualization of integrated endothelial nuclei colored by anatomical origin. Cortical and leptomeningeal endothelial cells occupy partially overlapping but distinct transcriptional states. (c) Volcano plot showing pseudobulk differential expression analysis comparing cortical and leptomeningeal endothelial cells. Differentially expressed genes were identified using DESeq2 pseudobulk analysis. Dashed lines indicate significance thresholds (adjusted P < 0.05 and |log2 fold change| > 0.5). (d) Heatmap of representative genes enriched in leptomeningeal endothelial cells. Expression values represent variance-stabilized transformed (VST) counts scaled by row. (e) Heatmap of representative genes enriched in cortical endothelial cells. Expression values represent row-scaled VST counts. (f) Representative coronal brain sections immunostained for ITGA2 (red) and Claudin-5 (green). ITGA2 expression is enriched in leptomeningeal vessels along the pial surface and is largely absent from cortical microvessels. (g) Representative leptomeningeal whole-mount images showing ITGA2 (red) and Claudin-5 (green) expression in leptomeningeal and cortical vascular networks. ITGA2 robustly labels leptomeningeal endothelial vessels but is minimally detected on cortical BBB vessels. (h) Quantification of ITGA2 protein expression. Left, ITGA2-positive area normalized to a 100 × 100 μm ROI. Right, percentage of Claudin-5-positive vascular area overlapping with ITGA2 signal. Each point represents an individual ROI. Statistical significance was determined by two-sided Wilcoxon rank-sum tests.

Pseudobulk differential expression analysis identified substantial transcriptional differences between cortical and leptomeningeal endothelial cells **(Fig. 3C)**. Leptomeningeal endothelial cells preferentially expressed genes associated with extracellular matrix remodeling and vascular interface function, including Itga2, Mmp28, Adgrg6, Mmrn1, and Slco2a1 **(Fig. 3D)**. In contrast, cortical endothelial cells preferentially expressed genes linked to neurovascular signaling and neuronal interaction programs, including Dpp10, Cntnap2, Grin2b, Nlgn1, Cntn1, and Grid2 **(Fig. 3E)**. These patterns were reproducible across biological replicates and supported origin-specific transcriptional specialization.

Because Itga2 was among the genes enriched in leptomeningeal endothelial cells, we next examined ITGA2 protein localization in leptomeningeal and cortical vessels. In coronal sections, ITGA2 signal was prominent along Cldn5-positive leptomeningeal vessels at the brain surface, whereas cortical parenchymal vessels exhibited little detectable ITGA2 signal **(Fig. 3F)**. Whole-mount preparations similarly revealed robust ITGA2 labeling throughout the leptomeningeal vascular network, with minimal ITGA2 signal associated with cortical vessels **(Fig. 3G)**. Quantification confirmed significantly greater ITGA2-positive area and ITGA2 overlap with Cldn5-positive vessels in leptomeningeal regions compared with cortex **(Fig. 3H)**. These data validate the transcriptomic enrichment of Itga2 at the protein level and support the conclusion that leptomeningeal endothelial cells exhibit a distinct extracellular matrix adhesion-associated phenotype.

To identify higher-order biological programs associated with each endothelial population, we performed Gene Ontology enrichment analysis on ranked differential expression results. Genes enriched in leptomeningeal endothelial cells were associated with leukocyte migration, myeloid leukocyte migration, antibacterial defense responses, and immune effector functions, whereas genes enriched in cortical endothelial cells were associated with synaptic signaling, glutamatergic transmission, neuronal plasticity, and neurovascular interaction pathways (**Extended Data Fig. 8A–B)**.

Consistent with these findings, module score analysis revealed enrichment of endothelial adhesion, extracellular matrix remodeling, membrane trafficking, and immune-interface programs in leptomeningeal endothelial cells **(Extended Data Fig. 8C)**. In contrast, cortical endothelial cells exhibited higher neurovascular interaction and canonical Wnt-BBB signaling module scores, whereas BBB transporter-associated programs were comparatively preserved between the two endothelial populations (Extended Data Fig. 8C). Representative genes contributing to each module exhibited corresponding origin-associated expression patterns across the integrated endothelial manifold **(Extended Data Fig. 8D)**.

Quality-control analyses supported the robustness of the integrated endothelial comparison. Endothelial nuclei from multiple cortical and leptomeningeal preparations contributed broadly across the integrated manifold rather than segregating solely into sample-specific clusters **(Extended Data Fig. 9A)**. Unsupervised clustering identified five endothelial subpopulations distributed across the integrated dataset **(Extended Data Fig. 9B)**, while quality-control metrics, including total transcript counts, detected genes, and mitochondrial transcript content, did not show an obvious relationship with anatomical origin **(Extended Data Fig. 9C)**. Sample-to-sample distance analysis separated cortical and leptomeningeal pseudobulk profiles while preserving relationships among biological replicates **(Extended Data Fig. 9D)**. In addition, scDblFinder analysis predicted low doublet frequencies across clusters, and predicted doublets did not account for the observed origin-associated transcriptional structure **(Extended Data Fig. 9E)**.

To further validate endothelial identity and evaluate whether origin-associated differences were simply explained by endothelial subtype composition or contamination, we examined canonical vascular zonation and lineage marker expression. Arterial, capillary, and venous markers were distributed across endothelial clusters, indicating that the integrated dataset contained endothelial cells spanning the vascular tree rather than a single vascular subtype **(Extended Data Fig. 10A)**. Canonical endothelial and BBB markers, including Cldn5, Slc2a1, Pecam1, Flt1, Mfsd2a, and Abcb1a, were broadly expressed across the integrated endothelial population **(Extended Data Fig. 10B)**. Conversely, mural, fibroblast, neuronal, glial, and myeloid markers showed limited expression across endothelial clusters, arguing against non-endothelial contamination as the primary driver of the observed origin-associated differences **(Extended Data Fig. 10C)**. Finally, unbiased cluster marker analysis identified additional transcriptional heterogeneity within endothelial subclusters, but these cluster signatures did not fully account for the segregation of cortical and leptomeningeal endothelial cells by anatomical origin **(Extended Data Fig. 10D)**.

Together, these analyses support the conclusion that leptomeningeal endothelial cells represent a specialized CNS endothelial population characterized by enhanced immune-interface, extracellular matrix remodeling, and adhesion-associated programs, whereas cortical endothelial cells preferentially express neurovascular interaction and canonical BBB-associated Wnt signaling programs.

### Leptomeningeal endothelial cells reside within a distinct Wnt ligand environment compared with cortical BBB endothelial cells

Comparison of cortical and leptomeningeal endothelial cells revealed widespread differences in transcriptional state, including altered expression of genes associated with extracellular matrix organization, vascular signaling, and canonical BBB identity **(Fig. 3)**. Because canonical Wnt/β-catenin signaling is a central regulator of blood–brain barrier development and maintenance, we next asked whether these distinct endothelial programs might reflect differences in the local signaling environments surrounding each vascular compartment. Specifically, we examined the cellular distribution of Wnt ligands within the leptomeninges and cortex to determine whether endothelial cells at these interfaces are exposed to distinct extracellular Wnt signaling niches.

To identify cellular sources of Wnt ligands, we analyzed independent single-cell and single-nucleus datasets from the leptomeninges, cortex, and combined leptomeningeal/dural tissues **(Fig. 4A– C)**. Clustering of leptomeningeal nuclei identified six major cell populations, including endothelial cells, macrophages, pia-associated fibroblasts, arachnoid-associated fibroblasts, dural border cells, and arachnoid barrier cells **(Fig. 4A)**. Analysis of cortical datasets identified endothelial, mural, glial, epithelial, and neuronal populations **(Fig. 4B)**, whereas the combined leptomeningeal/dural dataset provided an independent survey of fibroblast and barrier-cell populations across the meningeal compartment **(Fig. 4C)**. Cell-type identities were validated using lineage markers **(Extended Data Fig. 13B–C)**.

**Fig. 4:**
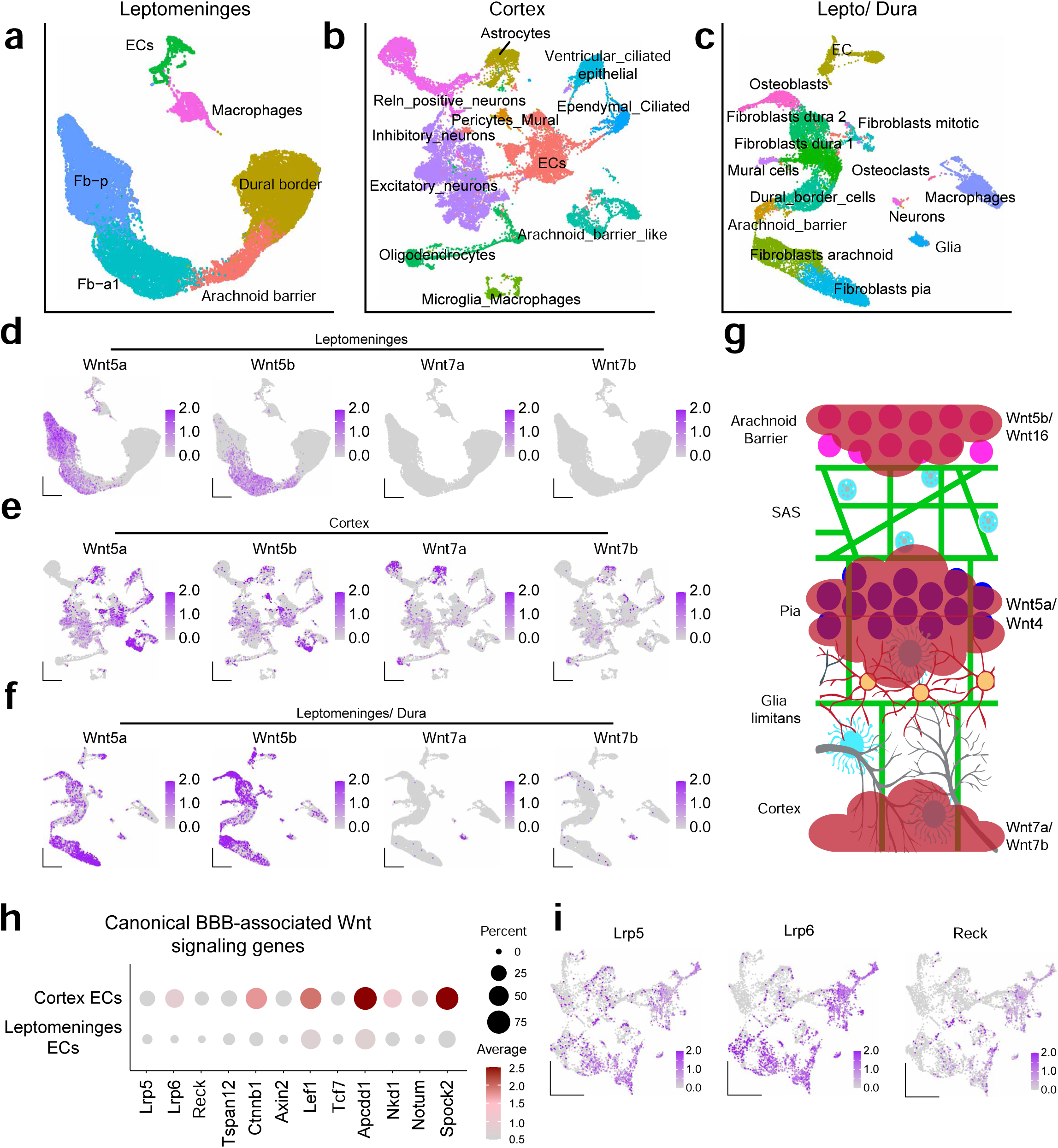
Distinct Wnt ligand environments in the leptomeninges and cortex. (a-c) UMAP visualization of cell populations from integrated single-nucleus RNA-sequencing datasets of the leptomeninges (A), cerebral cortex (B), and leptomeninges/dura (C). Leptomeningeal populations include endothelial cells (ECs), macrophages, pia-associated fibroblasts (Fb-p), arachnoid-associated fibroblasts (Fb-a1), dural border cells, and arachnoid barrier cells. Cortical populations include ECs, mural cells, astrocytes, oligodendrocytes, microglia/macrophages, ependymal and ventricular epithelial cells, and excitatory and inhibitory neuronal populations. (d-f) Feature plots showing expression of selected Wnt ligands across leptomeningeal (D), cortical (E), and leptomeningeal/dural (F) cell populations. Wnt5a expression was enriched within pia-associated fibroblasts, whereas Wnt5b and Wnt16 were enriched within arachnoid barrier-associated populations. In contrast, Wnt7a and Wnt7b expression was primarily detected within cortical parenchymal populations and was largely absent from leptomeningeal cell types. (g) Working model illustrating distinct Wnt signaling niches at CNS vascular interfaces. Cortical endothelial cells reside adjacent to Wnt7a/Wnt7b-producing parenchymal cells and exhibit robust canonical BBB-associated Wnt signaling. In contrast, leptomeningeal endothelial cells are positioned within a fibroblast-rich meningeal environment characterized by Wnt5a-, Wnt5b-, and Wnt16-expressing fibroblast and barrier-cell populations. (h) Dot plot comparing expression of canonical BBB-associated Wnt signaling pathway components in cortical and leptomeningeal endothelial cells. Dot size indicates the percentage of expressing cells and color indicates average normalized expression. Cortical endothelial cells exhibited greater expression of canonical BBB Wnt-response genes, including Apcdd1, Lef1, Tcf7, and Spock2, whereas leptomeningeal endothelial cells showed reduced expression of multiple downstream Wnt signaling targets. (i) Representative feature plots showing expression of the BBB-associated Wnt signaling components Lrp5, Lrp6, and Reck across integrated endothelial nuclei.

Examination of Wnt ligand expression revealed striking compartment-specific differences. Within the leptomeninges, Wnt5a expression was enriched in pia-associated fibroblasts, whereas Wnt5b and Wnt16 were concentrated within arachnoid barrier-associated populations **(Fig. 4D; Extended Data Fig. 11A)**. In contrast, expression of the canonical BBB-associated ligands Wnt7a and Wnt7b was largely absent throughout leptomeningeal populations. A broader survey of Wnt ligands, Frizzled receptors, and Norrin signaling components confirmed preferential expression of multiple non-canonical Wnt family members within leptomeningeal fibroblast and barrier-cell populations **(Extended Data Fig. 11)**.

Analysis of cortical cell populations demonstrated a markedly different pattern. Wnt7a and Wnt7b were readily detected within parenchymal cell populations, whereas Wnt5-family ligands showed a broader but less compartmentalized distribution **(Fig. 4E)**. A comprehensive survey of cortical Wnt ligands, Frizzled receptors, and Norrin pathway components further confirmed robust representation of canonical BBB-associated Wnt signaling components within the cortical environment **(Extended Data Fig. 12)**.

To determine whether these observations were reproducible across datasets, we next examined an independent leptomeningeal/dural atlas. Similar to the primary leptomeningeal dataset, Wnt5a, Wnt5b, and Wnt16 were enriched within fibroblast and barrier-associated populations, whereas Wnt7a and Wnt7b remained sparse and were largely localized to contaminating glial cells **(Fig. 4F)**. Analysis of the Allen Brain Cell Atlas yielded comparable expression patterns, further supporting segregation of Wnt5-family and Wnt7-family ligands between meningeal and parenchymal compartments **(Extended Data Fig. 13A)**.

Together, these analyses reveal that cortical and leptomeningeal vascular interfaces are embedded within distinct Wnt ligand environments. Cortical vessels reside within a parenchymal niche enriched for canonical BBB-associated Wnt7-family ligands, whereas leptomeningeal vessels are surrounded by fibroblast– and barrier-cell populations that preferentially express Wnt5-family ligands and other non-canonical Wnt pathway components **(Fig. 4G)**. These findings suggest that differences in local Wnt ligand availability may contribute to the distinct endothelial transcriptional programs observed between cortical and leptomeningeal vascular compartments.

Because canonical Wnt signaling is a central regulator of BBB endothelial identity, we next examined expression of BBB-associated Wnt pathway genes. Cortical endothelial cells expressed higher levels of multiple components of the canonical CNS endothelial Wnt signaling network, including Lrp5, Lrp6, Reck, Tspan12, Axin2, Lef1, Tcf7, Apcdd1, Nkd1, Notum, and Spock2 **(Fig. 4H)**. UMAP visualization confirmed that expression of representative pathway components, including Lrp5, Lrp6, and Reck, was concentrated within cortical endothelial populations and comparatively reduced throughout leptomeningeal endothelial cells **(Fig. 4I)**. These findings indicate that leptomeningeal endothelial cells exhibit reduced expression of the canonical BBB-associated Wnt transcriptional program relative to cortical BBB endothelial cells.

We next surveyed alternative pathways with potential links to β-catenin regulation, including non-canonical Wnt/planar cell polarity, VEGF/PI3K/AKT, and PGE2/cAMP/PKA signaling programs. Multiple planar cell polarity components, including Vangl2, Prickle2, Daam1, and Mapk9, were preferentially expressed in cortical endothelial cells, whereas downstream cytoskeletal regulators such as Rhoa and Rock2 were broadly expressed in both endothelial populations **(Extended Data Fig. 14A–B**). VEGF/PI3K/AKT and PGE2/cAMP/PKA pathway components exhibited broadly similar expression patterns across cortical and leptomeningeal endothelial cells, without clear selective enrichment in leptomeningeal endothelial cells **(Extended Data Fig. 14C–D)**.

### Structural organization of the leptomeningeal blood–CSF barrier differs from the cortical blood– brain barrier

The cellular architecture surrounding CNS blood vessels differs substantially between the leptomeningeal blood–cerebrospinal fluid barrier (LBCSFB) and the cortical blood–brain barrier (BBB). To define the structural organization of the LBCSFB, we compared mural cell coverage and glial organization in leptomeningeal and cortical vascular networks using whole-mount immunofluorescence imaging.

A schematic model illustrates the organization of leptomeningeal vessels within the subarachnoid compartment, positioned above the underlying glia limitans and overlying the pia-associated cellular layer, in contrast to cortical BBB vessels embedded within astrocytic endfeet **(Fig. 5A)**. Consistent with this organization, flatmount imaging of the pia showed Cldn5-positive vessels penetrating a pial-surface layer containing GFAP– and Aqp4-positive structures **(Fig. 5B)**. Imaging of the glia limitans revealed GFAP– and Aqp4-positive astroglial structures that were spatially distinct from much of the leptomeningeal vascular network, suggesting that leptomeningeal vessels are not ensheathed by astrocytic endfeet in the same manner as cortical BBB vessels **(Fig. 5C)**. High-contrast imaging further demonstrated that Aqp4-positive structures at the glia limitans showed only limited overlap with Cldn5-positive vessels in the leptomeningeal preparation, whereas cortical vessels exhibited substantially greater Aqp4 association **(Extended Data Fig. 15A–B)**.

**Fig. 5:**
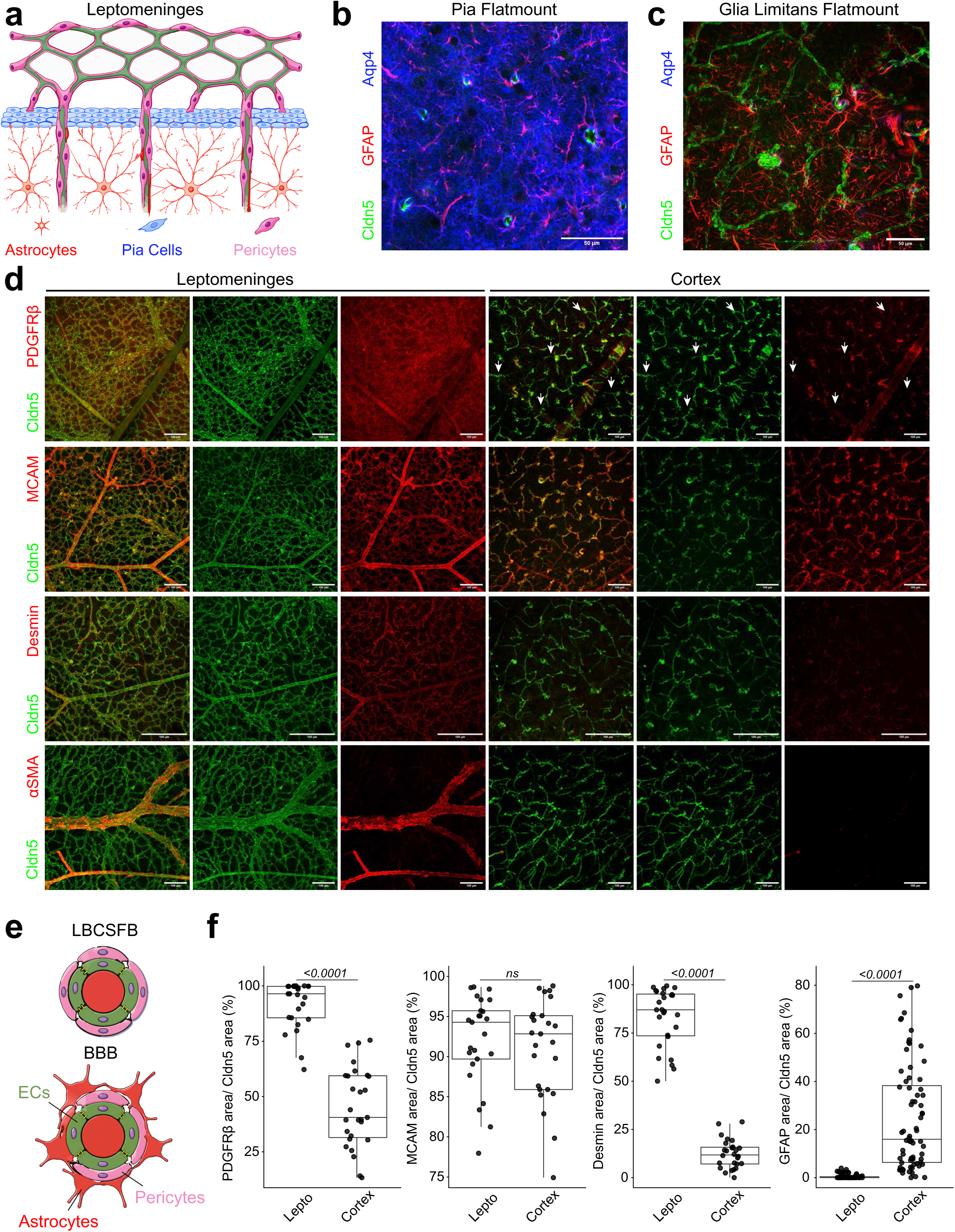
Structural organization of the leptomeningeal blood–CSF barrier and differential vascular coverage by pericytes and astrocytes across CNS barrier interfaces. (a) Schematic representation of the leptomeningeal architecture illustrating endothelial cells (green) forming a superficial vascular plexus within the subarachnoid space, associated with pial cells (blue) and extensive pericyte coverage (magenta). Astrocytes (red) are localized beneath the pia and extend projections toward penetrating vessels entering the cortex. (b) Representative flatmount immunofluorescence image of the pial surface stained for Cldn5 (green), GFAP (red), and Aqp4 (blue), demonstrating sparse astrocytic processes within the leptomeningeal compartment and broad Aqp4-positive pial coverage. (c) Representative flatmount image of the glial limitans/cortical surface showing dense GFAP-positive astrocytic endfeet surrounding cortical vessels. (d) Representative maximum-intensity projections of leptomeningeal and cortical vasculature stained for Cldn5 (green) together with PDGFRβ, MCAM, Desmin, or αSMA (red). Leptomeningeal vessels exhibit broad coverage by PDGFRβ-, Desmin-, and αSMA-positive mural cells, whereas cortical capillaries show reduced labeling for these markers. MCAM labeling remained highly associated with vessels in both compartments. White arrows indicate sparse marker-positive structures associated with cortical vessels. (e) Simplified schematic comparing mural and astrocytic organization at the LBCSFB versus the BBB. The LBCSFB is characterized by extensive pericyte coverage and minimal astrocytic association, whereas the BBB exhibits dense astrocytic endfoot coverage surrounding cortical vessels. (f) Quantification of marker overlap area normalized to total Cldn5-positive vascular area within leptomeningeal (SAS) and cortical compartments. PDGFRβ, Desmin, and GFAP coverage differed significantly between compartments, whereas MCAM coverage remained similar between the LBCSFB and BBB. Each point represents an individual ROI. Statistical comparisons were performed using two-sided Wilcoxon rank-sum tests after outlier removal using the 1.5× interquartile range method. Scale bars, 50 µm.

We next quantified vascular association with mural and glial markers by measuring the fraction of Cldn5-positive vascular area overlapping PDGFRβ, MCAM, Desmin, αSMA, or GFAP signal. Representative whole-mount images showed extensive PDGFRβ-, MCAM-, and Desmin-positive mural cell coverage along leptomeningeal vessels, whereas cortical vessels exhibited lower PDGFRβ and Desmin coverage and greater astroglial marker association **(Fig. 5D)**. αSMA signal was prominent along larger leptomeningeal vessels, consistent with smooth muscle investment of larger caliber vessels, but was not used as a global coverage metric across the vascular plexus **(Fig. 5D)**. A schematic summary highlights these compartment-specific structural differences: leptomeningeal vessels are closely associated with mural/perivascular cells and separated from astrocyte-rich glial structures, whereas cortical BBB vessels are surrounded by astrocytic endfeet in addition to pericytes **(Fig. 5E)**. Quantification confirmed that leptomeningeal vessels exhibited significantly greater PDGFRβ and Desmin coverage than cortical vessels, whereas MCAM coverage was high in both compartments and did not differ significantly **(Fig. 5F)**. In contrast, GFAP-positive astroglial coverage was abundant around cortical vessels but minimal around leptomeningeal vessels **(Fig. 5F)**.

Together, these findings indicate that the LBCSFB is a structurally distinct CNS vascular interface characterized by dense mural cell investment, limited astrocytic coverage, and spatial separation from the astrocyte-rich glia limitans. These architectural differences support the idea that leptomeningeal endothelial cells reside within a cellular microenvironment distinct from that of cortical BBB vessels.

### Leptomeningeal and cortical vessels exhibit compartment-specific responses during neonatal bacterial meningitis

The specialized molecular and structural organization of vascular barriers may shape how endothelial cells respond to pathological challenge. Although vascular dysfunction is a prominent feature of CNS infection, whether leptomeningeal vessels and the cortical blood–brain barrier (BBB) respond equivalently remains poorly defined. We therefore directly compared inflammatory activation, barrier dysfunction, and vascular remodeling between the leptomeningeal blood–CSF barrier (LBCSFB) and adjacent cortical vessels during neonatal bacterial meningitis **(Fig. 6)**.

**Fig. 6:**
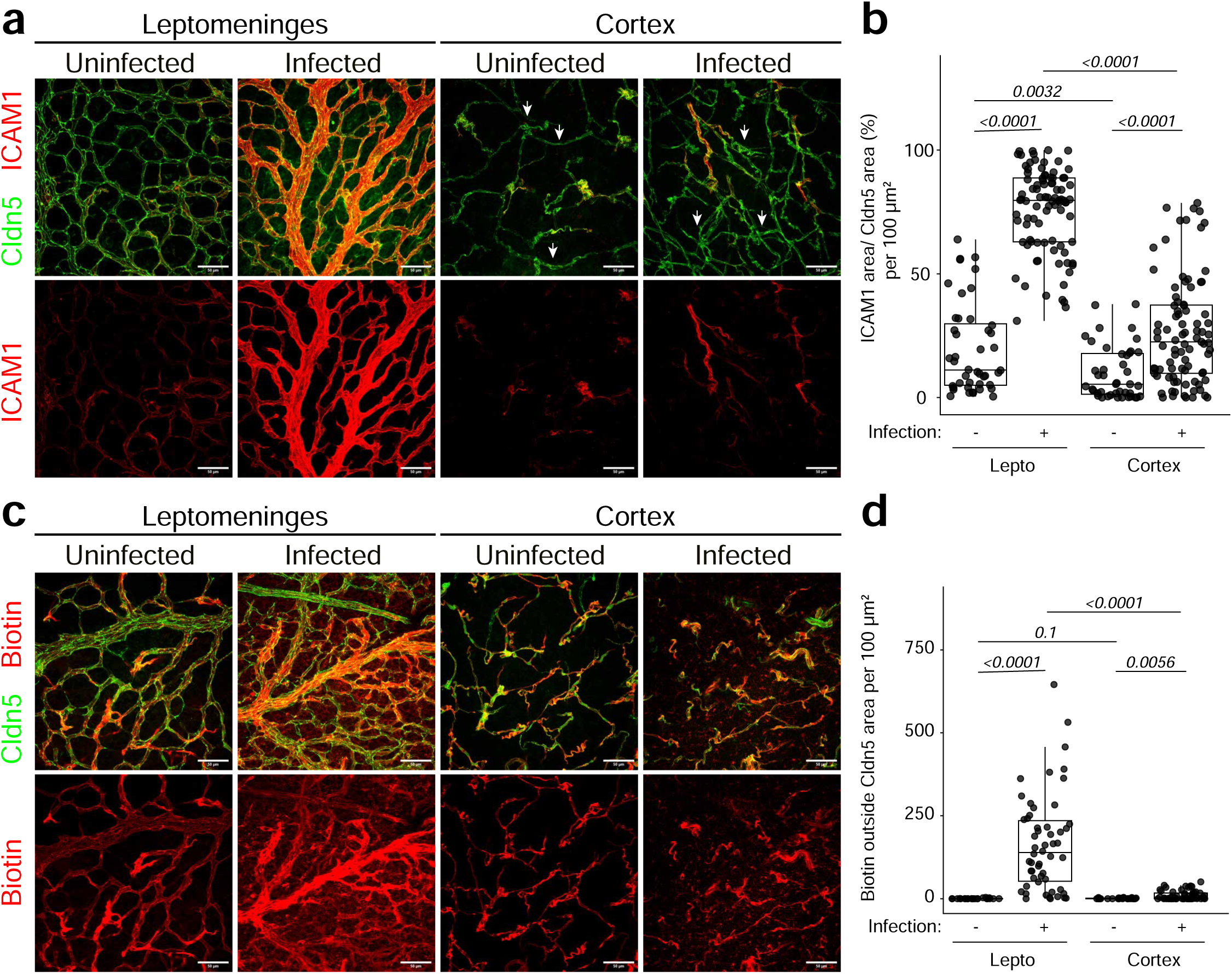
Compartment-specific vascular responses distinguish leptomeningeal and cortical barriers during neonatal meningitis. (a) Representative maximum-intensity projections of leptomeningeal and superficial cortical vessels from uninfected and *E. coli*–infected neonatal mice stained for Claudin-5 (Cldn5; green) and ICAM1 (red). Upper panels show merged images and lower panels show the ICAM1 channel alone. Leptomeningeal vessels displayed elevated baseline ICAM1 coverage compared to cortical vessels, with robust induction following infection. Cortical vessels exhibited comparatively lower baseline and infection-induced ICAM1 signal. Scale bars, 50 µm. (b) Quantification of ICAM1-positive vascular coverage across 100 × 100 µm ROIs. ICAM1 vascular coverage was quantified as the percentage of Cldn5-positive vessel area overlapping with ICAM1 signal within each ROI. Each dot represents one ROI. Boxplots show median and interquartile range. Statistical comparisons were performed using two-sided Wilcoxon rank-sum tests. (c) Representative maximum-intensity projections of sulfo-NHS-biotin tracer leakage in leptomeningeal and superficial cortical vessels from uninfected and *E. coli*–infected neonatal mice stained for Cldn5 (green) and biotin tracer (red). Upper panels show merged images and lower panels show the biotin channel alone. Infection induced prominent extravascular biotin accumulation surrounding leptomeningeal vessels, whereas cortical vessels exhibited substantially less tracer leakage. Scale bars, 50 µm. (d) Quantification of extravascular biotin leakage across 100 × 100 µm ROIs. Extravascular biotin signal was quantified as biotin-positive area located outside the Cldn5-positive vascular mask within each ROI. Each dot represents one ROI. Boxplots show median and interquartile range. Statistical comparisons were performed using two-sided Wilcoxon rank-sum tests.

Immunostaining for ICAM1 revealed a striking compartment-specific inflammatory response following infection **(Fig. 6A)**. ICAM1 expression was low under basal conditions but increased extensively throughout the leptomeningeal vascular network following infection, whereas vessels within the adjacent cortical parenchyma exhibited a more modest response. Quantification of ICAM1-positive area normalized to Cldn5-positive vascular area confirmed significantly greater inflammatory activation within leptomeningeal vessels **(Fig. 6B)**, which was maintained when measurements were averaged at the level of individual animals **(Extended Data Fig. 16A)**. Although infection increased Cldn5-positive vascular area within the leptomeningeal compartment **(Extended Data Fig. 16B)**, normalization relative to uninfected controls demonstrated that the greater ICAM1 response was not explained solely by altered vascular coverage **(Extended Data Fig. 16C)**. The masking workflow used for these measurements is shown in **Extended Data Fig. 16D**.

We next asked whether this enhanced inflammatory response was accompanied by preferential loss of barrier function. Infection produced extensive extravascular NHS-biotin accumulation throughout the leptomeningeal compartment, whereas leakage into the immediately underlying cortical parenchyma was substantially more limited **(Fig. 6C)**. Quantification confirmed a marked increase in extravascular biotin within infected leptomeninges compared with the cortex **(Fig. 6D)**, with the same pattern observed at the animal level **(Extended Data Fig. 16E)**. Measurements of Cldn5-positive vascular area and normalization of tracer leakage to available non-vascular area further demonstrated that this difference was not attributable to changes in vascular density or extravascular tissue area **(Extended Data Fig. 16F–G)**. The corresponding masking workflow is shown in **Extended Data Fig. 16H**. Together, these findings identify the LBCSFB as a preferential site of vascular inflammatory activation and barrier dysfunction during neonatal meningitis.

Barrier dysfunction was also accompanied by structural remodeling of the leptomeningeal vascular microenvironment. Laminin staining revealed disruption of the organized basement membrane network observed in uninfected vessels **(Extended Data Fig. 17A)**. Infection reduced laminin-positive area and mean intensity **(Extended Data Fig. 17B–C)** while increasing the number of discrete laminin-positive objects **(Extended Data Fig. 17D)**, consistent with fragmentation or loss of continuity within vascular basement membrane structures. Thus, acute LBCSFB dysfunction encompasses both endothelial barrier failure and remodeling of the surrounding extracellular matrix.

Together, these findings demonstrate that vascular responses to acute infection differ substantially across adjacent CNS vascular compartments. Relative to the cortical BBB, the LBCSFB exhibits greater inflammatory activation and vascular leakage, accompanied by remodeling of the leptomeningeal vascular basement membrane. These findings further support the LBCSFB as a functionally distinct vascular interface whose response to pathological challenge cannot be inferred directly from that of the cortical BBB.

### Human leptomeningeal endothelial cells exhibit transcriptional specialization distinct from the BBB

Our findings in mice identified the LBCSFB as a vascular compartment with molecular and functional properties distinct from the cortical BBB. We therefore asked whether this specialization was also evident in humans. We compared published human parenchymal brain and leptomeningeal single-nucleus RNA-sequencing datasets, restricting the analysis to donors without a clinical diagnosis of Alzheimer’s disease (AD)^13,31^. Both datasets contained arterial, capillary, and venous endothelial populations, with two transcriptionally distinct venous populations, vEndo_1 and vEndo_2, resolved in the leptomeninges **(Fig. 7A–B)**. Endothelial cells from both compartments retained canonical endothelial genes, including *PECAM1, VWF, KDR, FLT1,* and *CDH5*, together with the barrier-associated genes *CLDN5* and *SLC2A1* **(Fig. 7C)**, indicating preservation of core endothelial and barrier-associated programs.

**Fig. 7:**
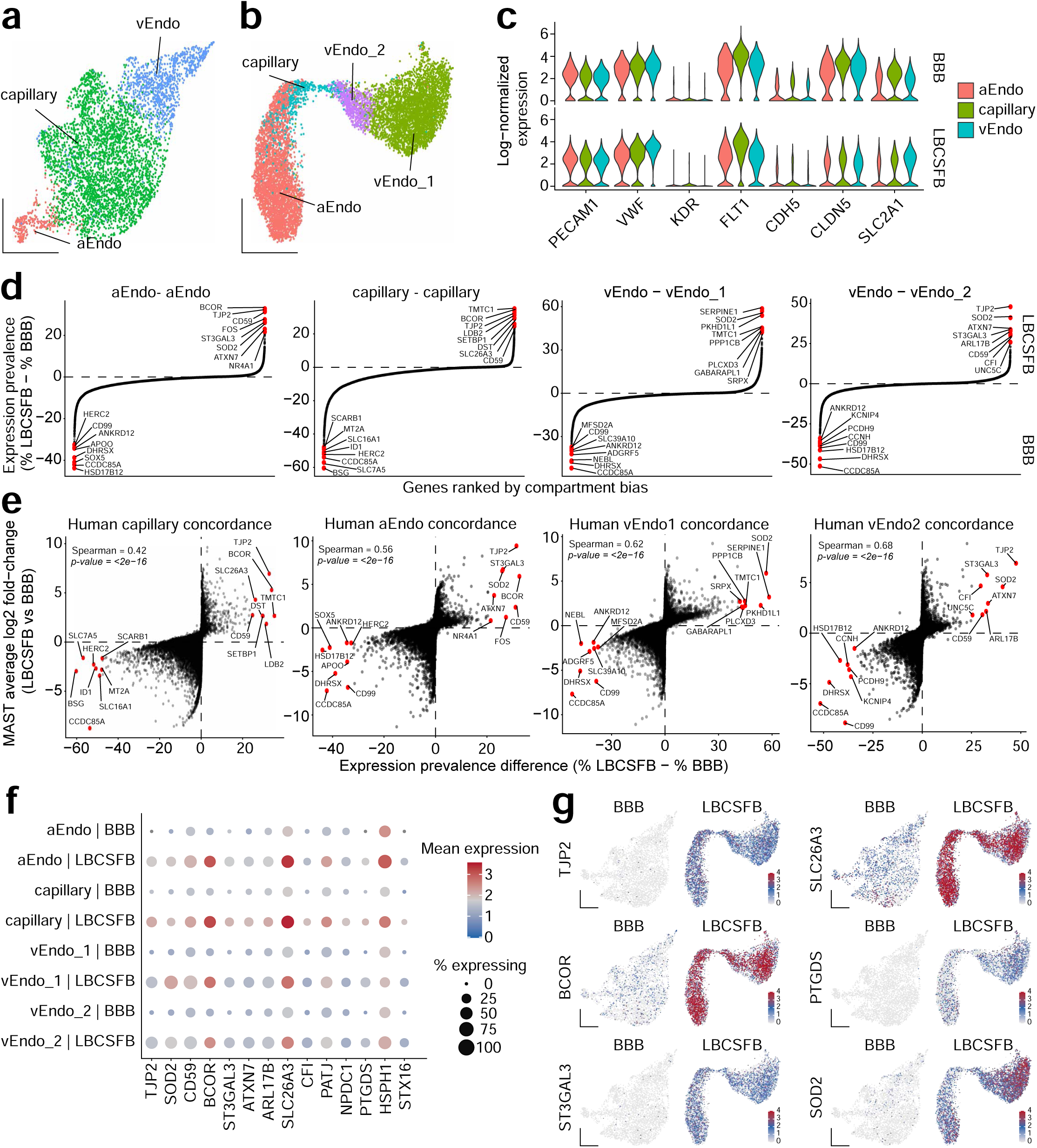
Human leptomeningeal endothelial cells exhibit transcriptional specialization distinct from the BBB. (a–b) UMAPs showing endothelial populations in the human cortical BBB dataset (a) and human leptomeningeal blood–cerebrospinal fluid barrier (LBCSFB) dataset (b). Endothelial cells were classified into arterial (aEndo), capillary, and venous (vEndo) populations. Two transcriptionally distinct venous populations, vEndo_1 and vEndo_2, were identified within the LBCSFB dataset. (c) Violin plots showing expression of canonical endothelial and blood–brain barrier-associated genes across matched arterial, capillary, and venous populations in the BBB and LBCSFB datasets. Expression is shown as log-normalized expression. (d) Ranked comparison of gene-expression prevalence between matched LBCSFB and BBB endothelial populations. Genes are ordered according to the difference in the percentage of cells expressing each gene (% LBCSFB − % BBB) for aEndo, capillary, vEndo_1, and vEndo_2 comparisons. Positive values indicate genes detected in a greater proportion of LBCSFB endothelial cells, whereas negative values indicate greater prevalence in the corresponding BBB population. Selected genes at the extremes of each distribution are labeled. (e) Concordance between expression-prevalence differences and differential-expression effect sizes for matched LBCSFB and BBB endothelial populations. Each point represents one gene. The x-axis indicates the difference in expression prevalence (% LBCSFB − % BBB), and the y-axis indicates the MAST average log2 fold change for LBCSFB relative to BBB. Spearman correlation coefficients and associated *P* values are shown for each comparison. Selected genes exhibiting strong compartment-associated differences are labeled. (f) Dot plot showing selected genes enriched across LBCSFB endothelial populations relative to their corresponding BBB populations. Genes were selected based on concordant evidence of LBCSFB enrichment, requiring a ≥10-percentage-point increase in expression prevalence, positive differential expression by MAST (average log2 fold change ≥ 1), and Benjamini–Hochberg-adjusted *P* < 0.05. Dot size indicates the percentage of cells expressing each gene and color indicates mean log-normalized expression among expressing cells. (g) Feature plots showing expression of *TJP2, SLC26A3, BCOR, PTGDS, ST3GAL3,* and *SOD2* across the complete BBB and LBCSFB endothelial manifolds. Expression is displayed on a common 0–4 log-normalized expression scale for each gene to facilitate direct comparison between compartments.

We next compared matched arterial, capillary, and venous populations to identify transcriptional features distinguishing the LBCSFB from the BBB. Genome-wide ranking by differences in expression prevalence revealed substantial compartment-associated transcriptional bias within each vascular class **(Fig. 7D)**. Numerous transcripts were preferentially detected in LBCSFB endothelial cells, with several LBCSFB-associated genes recurring across vascular populations. An independent MAST analysis produced concordant results: differences in expression prevalence correlated positively with MAST effect sizes in arterial (ρ = 0.56), capillary (ρ = 0.42), vEndo_1 (ρ = 0.62), and vEndo_2 (ρ = 0.68) comparisons **(Fig. 7E)**. Thus, compartment-associated differences were evident across complementary measures of transcript abundance.

We next identified genes recurrently enriched in LBCSFB endothelial cells across vascular classes. Applying combined thresholds for increased expression prevalence, MAST effect size, and statistical significance identified a shared LBCSFB-associated transcriptional signature across at least three of the four matched comparisons **(Fig. 7F)**. This signature included genes associated with junctional organization (*TJP2, PATJ*), transport (*SLC26A3*), complement regulation (*CD59, CFI*), and additional cellular programs, including *BCOR, SOD2, ST3GAL3,* and *PTGDS*. Visualization of representative genes across the complete endothelial manifolds confirmed their preferential expression within leptomeningeal endothelial cells **(Fig. 7G)**. Together, these analyses identify a human LBCSFB-associated transcriptional program that is superimposed on arterial, capillary, and venous endothelial identities.

Having established human LBCSFB specialization under non-AD conditions, we next asked whether leptomeningeal endothelial states were altered in neurodegenerative disease. Reanalysis of the leptomeningeal dataset across NCI/MCI and AD donors confirmed arterial, capillary, vEndo_1, and vEndo_2 populations, with both sexes and multiple donors represented across the endothelial populations and clinical groups **(Extended Data Fig. 18A–D)**. Donor-level pseudobulk analysis revealed subtype– and sex-dependent AD-associated transcriptional changes **(Extended Data Fig. 18E–F)**. Within vEndo_1 cells, female AD donors exhibited a prominent reduction in *KYNU*, whereas males showed a distinct response characterized by increased *F8, BMPR1B, PTPN13, NAV3, RDH10,* and *MEIS2* **(Extended Data Fig. 18G–H)**. Donor-level normalized expression confirmed reduced *KYNU* in female AD vEndo_1 cells (log2FC = −1.98, FDR = 2.8 × 10⁻⁵), with no corresponding change in males (log2FC = −0.06, FDR = 1.0) **(Extended Data Fig. 18I)**. Although exploratory, these findings indicate that human leptomeningeal endothelial populations undergo additional subtype– and sex-associated remodeling in AD.

Collectively, these human data extend LBCSFB specialization beyond the mouse, identifying a transcriptional program that distinguishes leptomeningeal endothelial cells from matched BBB populations while preserving core endothelial and barrier features. Moreover, this specialized endothelium remains responsive to disease, exhibiting additional transcriptional remodeling in AD.

## Discussion

Early studies demonstrated that the cerebrospinal fluid of healthy individuals is enriched for central memory CD4+ T cells and identified selective expression of P-selectin and ICAM-1 on leptomeningeal and choroid plexus vessels, but not parenchymal microvessels, suggesting that leptomeningeal vessels participate in physiological immune surveillance by regulating leukocyte entry into the CSF^32^. During autoimmune neuroinflammation, intravital two-photon microscopy showed that autoreactive T cells initially arrest on leptomeningeal post-capillary venules, crawl along these vessels before interacting with leptomeningeal antigen-presenting cells and subsequently invading the CNS parenchyma^33^. Subsequent work established the subarachnoid space as an active immunological compartment by showing that antigen-presenting cells within the leptomeninges locally reactivate encephalitogenic T cells, thereby licensing their entry into the brain^34^. More recently, single-cell studies have shifted attention toward defining the immune cell composition of the human leptomeninges, revealing specialized resident immune populations that undergo disease-associated remodeling across neurodegenerative disorders^35^. Collectively, these studies establish the leptomeninges as a highly specialized neuroimmune interface in which the blood vasculature serves as a portal for immune surveillance and leukocyte entry into the CNS. However, despite more than two decades of functional studies, the endothelial cells composing this vascular network have remained largely undefined. Our findings provide the missing molecular framework for these observations by demonstrating that leptomeningeal endothelial cells constitute a transcriptionally specialized vascular population distinct from cortical BBB endothelium. These findings suggest that the unique immunological functions of the leptomeningeal vasculature arise, at least in part, from intrinsic endothelial specialization rather than anatomical location alone.

Our findings suggest that the specialized molecular identity of leptomeningeal endothelial cells may reflect the markedly different cellular and signaling environment in which these vessels reside. Studies of the BBB and blood–retinal barrier have established that local ligand availability can directly shape endothelial barrier identity. Neural-derived Wnt ligands activate endothelial β-catenin signaling during CNS angiogenesis and induce BBB-associated programs, whereas Norrin/FZD4/LRP5 signaling controls a related but anatomically distinct endothelial program in the retina^9,24,36^. Genetic studies further demonstrated that endothelial responsiveness to these ligands depends on specialized receptor and co-receptor complexes. GPR124 and RECK selectively enable Wnt7a/Wnt7b signaling in CNS endothelium, whereas FZD4, LRP5 and TSPAN12 form the core signaling machinery for Norrin-dependent vascular development and barrier function^22,23,26^. Importantly, these signaling systems are not deployed uniformly across the CNS. Norrin and Wnt7a/Wnt7b signaling make overlapping but regionally distinct contributions to blood–brain and blood–retinal barrier development and maintenance, demonstrating that related endothelial barrier phenotypes can be generated through different combinations of extracellular ligands and endothelial signaling machinery^37^.

Against this framework, the leptomeningeal vascular niche is strikingly different. Leptomeningeal endothelial cells reside outside the neural parenchyma, lack the extensive astrocytic endfoot association characteristic of cortical BBB vessels, and are instead embedded within a fibroblast-rich stromal compartment with greater mural cell investment. Correspondingly, we find that this niche is depleted for transcriptional evidence of the canonical Wnt7a/Wnt7b ligand environment that characterizes the cortical neurovascular unit and is instead enriched for fibroblast– and barrier-cell-derived Wnt5-family ligands. Leptomeningeal endothelial cells themselves also express lower levels of multiple canonical BBB-associated Wnt receptors, co-receptors and downstream transcriptional targets, indicating that the difference extends beyond ligand availability to the signaling competence of the endothelium itself. Wnt5A can directly alter vascular endothelial behavior through non-canonical pathways, including cytoskeletal remodeling, migration, extracellular matrix-associated gene expression and changes in endothelial permeability^38,39^. These observations raise the possibility that a Wnt5-rich stromal environment contributes to the distinct endothelial state of the leptomeninges. Our data do not establish Wnt5-family signaling as a causal determinant of leptomeningeal endothelial identity, however. Rather, the coordinated differences in extracellular ligand availability, endothelial receptor machinery and surrounding cellular architecture support a broader model in which CNS vascular specialization emerges from interactions between local signaling niches and the intrinsic signaling capacity of regional endothelial populations.

Our findings also suggest that extracellular matrix organization is an important component of leptomeningeal vascular specialization. Leptomeningeal endothelial cells were enriched for adhesion– and extracellular matrix-associated transcriptional programs, including increased expression of Itga2, Mmrn1, and Mmp28, and ITGA2 protein was selectively enriched along leptomeningeal vessels relative to cortical vessels. These molecular differences were accompanied by greater mural cell association within the leptomeningeal vascular network, consistent with the broader concept that endothelial cells and perivascular cells jointly shape vascular basement membrane composition and organization^40,41^. Studies of the cortical BBB have similarly shown that endothelial cells, pericytes, and astrocytes contribute distinct extracellular matrix components to the vascular basement membrane and that laminin-rich matrix interactions support vascular stability and barrier integrity^19,42,43^. In this context, the distinct cellular architecture surrounding leptomeningeal vessels may be expected to generate a basement membrane environment that differs from that of the cortical neurovascular unit. Consistent with this possibility, we observed substantial remodeling of laminin-positive vascular structures during neonatal meningitis, including reduced laminin area and intensity together with increased fragmentation. These findings do not establish whether extracellular matrix remodeling is a cause or consequence of leptomeningeal barrier dysfunction, but they indicate that vascular injury at this interface involves coordinated changes in endothelial state and basement membrane organization. Together, the transcriptional, structural, and infection-associated data support a model in which endothelial–matrix interactions represent an additional axis of specialization at the leptomeningeal vascular interface, alongside differences in canonical barrier signaling and cellular composition.

Our findings further demonstrate that specialization of the leptomeningeal vasculature extends to its response to inflammatory challenge. Early studies of human immune surveillance identified expression of endothelial adhesion molecules, including ICAM1, along leptomeningeal and choroid plexus vessels relative to parenchymal microvessels^32^, while intravital imaging during autoimmune neuroinflammation subsequently demonstrated that autoreactive T cells preferentially arrest and crawl along leptomeningeal post-capillary venules before entering the CNS^33^. Local antigen presentation within the subarachnoid space further promotes reactivation of encephalitogenic T cells following their entry into the leptomeningeal compartment, emphasizing that vascular recruitment occurs within a broader tissue environment specialized for immune surveillance and inflammatory responses^34^. Consistent with the importance of endothelial activation in human disease, increased endothelial adhesion molecules, including ICAM1, have also been associated with intrathecal inflammation and leukocyte recruitment during bacterial meningitis^44^. Our previous studies established that neonatal bacterial meningitis produces widespread inflammatory activation within the leptomeninges^10^ and that endothelial signaling contributes directly to leptomeningeal inflammation, Claudin-5 internalization, and vascular barrier dysfunction during infection (Seegren et al., 2026). The present findings extend these observations by directly comparing leptomeningeal and cortical vessels within the same infectious challenge. Despite exposure to the same systemic infection, the two vascular compartments exhibited markedly different responses, with substantially greater ICAM1 induction and vascular permeability in leptomeningeal vessels, accompanied by remodeling of their vascular basement membrane. Thus, the distinction between the LBCSFB and cortical BBB is not restricted to their basal transcriptional and structural properties but extends to how these vascular interfaces respond to inflammatory challenge. Together with the preferential involvement of leptomeningeal vessels in leukocyte trafficking during autoimmune neuroinflammation, these findings suggest that regional endothelial specialization may contribute to the distinct susceptibility and function of CNS vascular interfaces during disease.

The disease-associated remodeling observed in human leptomeningeal endothelial cells suggests that this vascular interface may also be altered in chronic neurodegenerative disease. Particularly notable was the selective reduction of *KYNU* in vEndo_1 endothelial cells from female AD donors. *KYNU* encodes kynureninase, a key enzyme in the kynurenine pathway of tryptophan metabolism, which has been extensively implicated in AD and neuroinflammation^45^. Human AD studies have identified altered circulating kynurenine-pathway metabolites^46^, while amyloid-β induces *KYNU* and other pathway enzymes in human microglia^47^. More recently, integrated human CSF, blood, and brain analyses linked tryptophan-kynurenine metabolism with amyloid-β and tau pathology in AD^48^. Our findings extend these observations to a defined leptomeningeal endothelial population, suggesting that disease-associated remodeling of this vascular interface may involve metabolic pathways already implicated in AD pathophysiology. Given the small cohort and sex specificity of this effect, its functional significance remains to be determined.

Several questions remain regarding how the specialized identity of the LBCSFB is established and maintained. Although our data identify coordinated differences in the leptomeningeal Wnt ligand environment and endothelial Wnt signaling machinery, they do not establish whether altered Wnt signaling is responsible for the distinct leptomeningeal endothelial state. In particular, the relative contributions of reduced Wnt7a/Wnt7b signaling, increased exposure to non-canonical Wnt ligands, and other stromal signals will require direct genetic and functional testing. Similarly, the enrichment of ITGA2 and other extracellular matrix-associated programs identifies endothelial–matrix interactions as a prominent feature of leptomeningeal endothelial cells but does not establish whether these programs instruct vascular identity or instead reflect adaptation to the surrounding stromal and basement membrane environment. Our functional analyses are also limited to vascular permeability during neonatal bacterial meningitis and therefore do not define the basal permeability or molecular selectivity of the LBCSFB, nor whether its response differs from the cortical BBB across other inflammatory, developmental, or age-associated conditions. Finally, although human leptomeningeal atlases contain endothelial populations with transcriptional features consistent with those identified here, direct cross-species and spatial validation will be necessary to determine the extent to which the molecular organization defined in mice is conserved in humans. These limitations highlight a broader need to understand when and how leptomeningeal endothelial identity emerges during development, which local signals maintain this state in adulthood, and how these programs are remodeled during aging and disease. Nevertheless, our findings demonstrate that endothelial cells with canonical CNS barrier features should not necessarily be considered a functionally uniform vascular population. By defining the LBCSFB as a molecularly, structurally, and functionally specialized vascular interface, this study expands the organization of CNS barriers beyond the classical cortical BBB and provides a framework for investigating how regional endothelial specialization shapes CNS homeostasis and disease.

## Supporting information

GSE_Annotation_Table

## Acknowledgements

Supported by the Howard Hughes Medical Institute. This work was initiated in the laboratory of Jeremy Nathans. I thank Jeremy Nathans for his mentorship, scientific insight, continued encouragement, and support throughout the development of this project. I thank Zhongming Li and Amir Rattner for providing the mouse snRNA-seq datasets.

The results published here are in whole or in part based on data obtained from the AD Knowledge Portal Community Data Contribution Program. We are grateful to those who agreed to donate their brains for research. We thank all the employees at RADC for their support and assistance. The original study was supported by NIA grants R01AG074082, R01AG079223, P30AG10161, P30AG72975, R01AG015819, R01AG017917, U01AG61356, and R01AG061798.

## Data Availability

Mouse snRNA-seq objects for leptomeninges can be obtained from Seegren, et al 2026 – eLife. https://doi.org/10.7554/eLife.110458.3

Mouse snRNA-seq objects for leptomeninges/dura can be obtained form Wang, et al 2023 – eLife. https://doi.org/10.7554/eLife.86130

Mouse snRNA-seq objects for cortical ECs can be obtained upon reasonable request

All GSE studies used from GRIEN Database^49^ – List of GSE Annotations can be found in Supplemental File 1

Human leptomeningeal single-nucleus RNA-sequencing data were obtained from Kearns et al. (2023) through the AD Knowledge Portal Community Data Contribution Program (Synapse: syn54847635; https://doi.org/10.1038/s41467-023-42825-y) under the applicable controlled-access data-use terms.

Human cortical single-nucleus RNA-sequencing count matrices and metadata were obtained from the publicly available dataset provided by Sun et al. (2023) through the study website (http://compbio.mit.edu/scADbbb/) and used as the reference dataset for comparison with human leptomeningeal endothelial cells.

## Methods

### Construction of the cross-study endothelial RNA-seq atlas (Bulk RNA-seq Meta-analysis)

Publicly available bulk RNA-sequencing datasets were identified through the GREIN database and manually curated to generate a cross-study atlas of endothelial and CNS-associated cell populations. Studies were included if raw gene-level count matrices and sufficient accompanying metadata were available to assign tissue and cell identity. Metadata from each study were manually harmonized and annotated with simplified cell-class labels (simpleID), age category, experimental methodology, and study identifier. Individual count matrices were merged into a unified gene-by-sample expression matrix for downstream analysis.

Gene expression normalization was performed using DESeq2 size-factor normalization. Variance-stabilized expression values were generated using the DESeq2 variance stabilizing transformation and used for principal component analysis (PCA). PCA was performed using the most variable genes within the dataset and visualized using ggplot2. Metadata were further used to summarize sample composition, age distribution, study representation, and experimental capture methodology across the atlas.

### Validation of sample identity and marker gene expression (Bulk RNA-seq Meta-analysis)

To evaluate sample purity and validate cell-type annotations, expression of established endothelial, pericyte, microglial, and parenchymal marker genes was examined across all sample groups. Normalized expression values were calculated from DESeq2 size-factor–normalized counts and transformed as log2(normalized counts + 1). Expression distributions of canonical endothelial markers (Pecam1, Cdh5, Kdr, Esam, and Robo4) were visualized using boxplots across all sample groups.

To provide a global assessment of marker enrichment, dot plots were generated for representative endothelial (Pecam1, Cdh5, Kdr, Esam, Robo4), pericyte (Pdgfrb, Cspg4, Des, Notch3, Rgs5), microglial (Cx3cr1, P2ry12, Tmem119, Aif1, Hexb), and parenchymal (Rbfox3, Slc17a7, Aqp4, Aldh1l1, Mbp) marker genes. Dot color represents the mean log2(normalized counts + 1) expression within each sample group. Dot size represents the percentage of samples within a group whose expression met or exceeded a reference threshold calculated from the cognate cell type as the mean expression minus one-half standard deviation. This approach provides a combined measure of marker abundance and consistency across independently generated datasets.

### Differential expression analysis of brain endothelial cells (Bulk RNA-seq Meta-analysis)

To identify genes enriched in brain endothelial cells relative to other endothelial populations, samples annotated as brain endothelial cells were compared with pooled peripheral endothelial samples (lung, liver, heart, kidney, intestinal, and bone marrow endothelial cells) using DESeq2. Samples were subset from the pooled RNA-seq atlas, genes with fewer than 10 total counts across all samples were excluded, and differential expression was performed using a negative binomial generalized linear model with cell group specified as the experimental design. Size-factor normalization and dispersion estimation were performed using the standard DESeq2 workflow.

Log2 fold-change estimates were shrunk using the adaptive Bayesian shrinkage estimator implemented in the apeglm package (lfcShrink, type = “apeglm”) to improve effect-size estimation for low-count and highly variable genes while preserving large, well-supported expression differences. Statistical significance was determined using Wald tests with Benjamini–Hochberg correction for multiple testing. Volcano plots display shrunken log2 fold changes together with adjusted P values. For visualization, genes with a normalized mean expression (baseMean) <10 or absolute shrunken log2 fold change <0.1 were excluded from plotting, and genes were considered significantly differentially expressed when the adjusted P value was <0.05 and the absolute shrunken log2 fold change exceeded 1.

To further evaluate brain endothelial specificity within the CNS, a second DESeq2 analysis compared brain endothelial samples with pooled CNS non-endothelial samples (microglia, pericytes, and whole-brain tissue) using the same normalization, statistical framework, fold-change shrinkage, and significance thresholds. Shrunken log2 fold changes from both comparisons were subsequently integrated with cross-study reproducibility metrics to prioritize robust brain endothelial-enriched candidate genes.

### Identification of sample group–enriched transcriptional signatures (Bulk RNA-seq Meta-analysis)

To identify genes enriched within each sample group, mean normalized expression was calculated across all samples belonging to a given group and compared with the mean expression across all remaining groups. Gene enrichment was defined as the difference between mean expression within the focal group and mean expression across all other groups. Candidate marker genes were required to exhibit a mean expression greater than 1 and an enrichment score greater than 1.

To improve interpretability, genes corresponding to predicted transcripts (Gm family), RIKEN cDNA clones, LOC-designated genes, ribosomal genes, mitochondrial genes, and hemoglobin genes were excluded prior to marker selection. For each sample group, the five genes with the highest enrichment scores were selected. Mean expression values for these genes were then calculated across all sample groups and visualized as a heatmap.

For visualization, expression values were row-scaled by z-score transformation to emphasize relative enrichment patterns across tissues and cell types. Heatmap rows and columns were manually ordered to preserve the relationship between each sample group and its corresponding enriched transcriptional signature.

### Brain endothelial study reproducibility analysis (Bulk RNA-seq Meta-analysis)

To evaluate the consistency of BBB-associated transcriptional programs across independent studies, samples annotated as brain endothelial cells were extracted from the pooled endothelial RNA-seq atlas. Gene-level counts were normalized using DESeq2 size-factor normalization and transformed as log2(normalized counts + 1) for visualization.

Principal component analysis (PCA) was performed using DESeq2 variance-stabilized expression values and the 500 most variable genes. PCA plots were used to assess whether transcriptomic variation among brain endothelial samples was associated with study origin, animal age, or sample isolation methodology.

To evaluate preservation of the canonical BBB transcriptional program, expression of established BBB marker genes (*Slco1c1, Mfsd2a, Slc2a1, Cldn5, Abcb1a,* and *Abcg2*) and BBB-associated Wnt signaling components (*Lrp5, Lrp6, Reck, Tspan12, Ctnnb1, Lef1, Tcf7, Apcdd1, Nkd1, Notum,* and *Spock2*) was quantified across individual studies. Expression values are displayed as log2(normalized counts + 1) for each sample, with boxplots summarizing study-level distributions.

### Non-endothelial lineage and endothelial subtype/state program scoring (Bulk RNA-seq Meta-analysis)

To determine whether transcriptomic heterogeneity among independent brain endothelial cell (Brain EC) studies could be explained by variable contributions of non-endothelial transcripts, we quantified lineage-specific expression programs within each Brain EC sample. Brain endothelial samples were extracted from the pooled atlas and analyzed separately using DESeq2-normalized expression values. For lineage scoring, normalized expression values were transformed as log2(normalized counts + 1) and subsequently standardized by gene using z-score normalization across Brain EC samples.

Canonical endothelial identity was assessed using the genes *Pecam1, Cdh5, Kdr, Flt1, Tek, Esam, Robo4, Erg, Slco1c1, Mfsd2a, Slc2a1, Cldn5, Abcb1a,* and *Abcg2*. Non-endothelial lineage programs were defined using curated marker sets representing neuronal (*Rbfox3, Snap25, Syp, Map2, Tubb3, Slc17a7, Slc17a6, Gad1,* and *Gad2*), astroglial (*Aqp4, Aldh1l1, Gfap, Slc1a2, Slc1a3, Gja1, S100b,* and *Sox9*), oligodendrocyte (*Mbp, Mog, Plp1, Mobp, Mag, Cnp, Myrf, Olig1,* and *Olig2*), mural (*Pdgfrb, Rgs5, Cspg4, Notch3, Acta2, Tagln, Myh11, Myl9, Des, Kcnj8,* and *Abcc9*), and myeloid (*P2ry12, Cx3cr1, Tmem119, Aif1, Ctss, C1qa, C1qb, C1qc, Hexb, Trem2, Tyrobp,* and *Laptm5*) cell populations.

For each lineage, a lineage score was calculated as the mean z-score of all genes within the corresponding marker set:

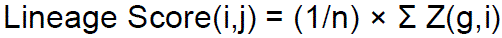

where *Z(g,i)* is the z-score normalized expression of gene *g* in sample *i*, and *n* is the number of genes within lineage program *j*.

A total non-endothelial lineage score was then calculated for each sample as the arithmetic mean of the neuronal, astroglial, oligodendrocyte, mural, and myeloid lineage scores:

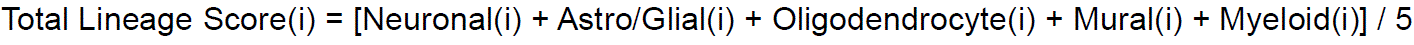

Study-level lineage scores were generated by averaging sample-level scores within each GEO study. Heatmaps display study-level mean lineage z-scores. For visualization, study-level lineage scores were row-scaled by z-score transformation across studies.

To compare endothelial and non-endothelial transcriptional contributions directly, absolute endothelial and non-endothelial expression scores were additionally calculated using log2(normalized counts + 1) expression values without z-score transformation. Endothelial scores were calculated using the canonical endothelial marker set described above. Non-endothelial scores were calculated by combining all genes from the neuronal, astroglial, oligodendrocyte, mural, and myeloid marker sets. Mean expression values were calculated for each study and displayed as bar plots of log2(normalized counts + 1) expression.

To determine whether non-endothelial lineage signatures explained variability among Brain EC studies, principal component analysis was performed using variance-stabilized expression values from Brain EC samples only. Sample-level lineage scores were mapped onto PCA coordinates, and associations between lineage scores and principal components were assessed using Spearman rank correlation.

To investigate whether residual transcriptomic heterogeneity reflected endothelial subtype composition rather than non-endothelial transcript contribution, additional endothelial subtype and state programs were quantified. Capillary markers included *Mfsd2a, Slco1c1, Abcb1a, Abcg2, Cldn5, Ocln, Slc2a1, Tfrc, Spock2, Apcdd1, Reck,* and *Tspan12*. Arterial markers included *Gja5, Efnb2, Sox17, Bmx, Fbln5, Cxcl12, Hey1, Hey2, Jag1, Notch4,* and *Cxcr4*. Venous markers included *Nr2f2, Vwf, Slc38a5, Ackr1, Emcn, Aplnr, Ephb4, Flt4,* and *Plvap*. Angiogenic markers included *Apln, Angpt2, Kdr, Adm, Esm1, Nrp1, Dll4,* and *Vegfc*. Activated/inflammatory endothelial markers included *Icam1, Vcam1, Sele, Selp, Ccl2, Cxcl10, Nfkbia, Socs3, Stat1, Irf1, Ifit1,* and *Isg15*.

For each sample, endothelial subtype and state scores were calculated as the mean log2(normalized counts + 1) expression of genes within each program:

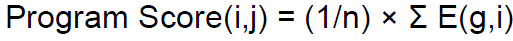

where *E(g,i)* is the log2(normalized counts + 1) expression value of gene *g* in sample *i*, and *n* is the number of genes within endothelial program *j*.

A total endothelial program score was additionally calculated as the arithmetic mean of the capillary, arterial, venous, angiogenic, and activated/inflammatory program scores:

Total Endothelial Program(i) = [Capillary(i) + Arterial(i) + Venous(i) + Angiogenic(i) + Activated/Inflammatory(i)] / 5

Study-level endothelial subtype and state scores were generated by averaging sample-level scores within each GEO study. Heatmaps display study-level mean program scores following row-wise z-score scaling across studies. Absolute program expression values were visualized as study-level mean log2(normalized counts + 1) expression values.

Associations between subtype/state scores and PCA coordinates were evaluated using Spearman rank correlation. To compare the relative explanatory power of lineage and endothelial programs, squared Spearman correlation coefficients were calculated as:

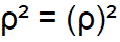

where *ρ* represents the Spearman rank correlation coefficient between a lineage or endothelial program score and the corresponding PCA coordinate. Values of ρ² were used as a measure of the relative association between individual programs and the major axes of variation identified by PCA.

Total endothelial program scores were used as a summary measure of endothelial-state variation and were not incorporated into PCA construction.

### Study-correction sensitivity analysis (Bulk RNA-seq Meta-analysis)

Brain endothelial samples were analyzed separately using the DESeq2 variance-stabilized expression matrix. Raw PCA was performed on the 500 genes with the highest variance across Brain EC samples. PCA was performed with prcomp on the transposed expression matrix, so that samples were treated as observations and genes as variables.

To assess study-associated variation, metadata factors were tested against PC1 and PC2 using one-way ANOVA. Separate models were fit for each metadata factor:

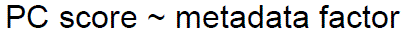

where the metadata factor was study identity, age category, or isolation method. For each model, η² was calculated as:

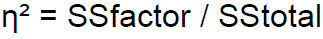

where SSfactor is the sum of squares explained by the metadata factor and SStotal is the total sum of squares for that PC axis.

To remove study-associated variation, the Brain EC variance-stabilized expression matrix was corrected using limma::removeBatchEffect, with GEO study identity supplied as the batch variable. No additional design matrix was included, so no age or isolation-method structure was explicitly protected during correction. PCA was then repeated on the study-corrected matrix using the same procedure: the 500 most variable genes were selected, and PCA was performed using prcomp.

To determine whether gene-level PC-associated structure persisted after study correction, Spearman correlations were calculated between each expressed gene and PC1 or PC2 in the raw PCA and again between each gene and PC1 or PC2 in the study-corrected PCA. For each gene:

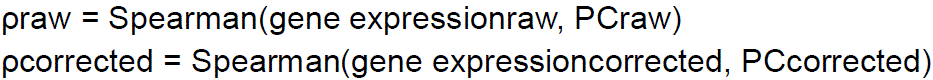

Gene-level retention was assessed by comparing ρraw and ρcorrected. For genes with strong raw PC association, defined as |ρraw| ≥ 0.5, we calculated the percentage of genes that retained corrected |ρ| ≥ 0.3, the percentage that retained corrected |ρ| ≥ 0.5, and the percentage that preserved the same direction of association after correction. We also calculated the fraction of gene-level association retained as:

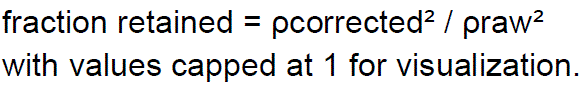

### Functional enrichment analysis of retained PC2-associated genes (Bulk RNA-seq Meta-analysis)

Genes exhibiting retained positive or retained negative associations with corrected PC2 were analyzed separately for functional enrichment. Gene Ontology (GO) Biological Process enrichment analysis was performed using the clusterProfiler package with the expressed Brain EC gene set used as the background universe. Significance was assessed using Benjamini–Hochberg multiple-testing correction. Enriched biological processes were ranked according to adjusted p-value and visualized using dot plots generated with enrichplot.

### Heatmap generation (Bulk RNA-seq Meta-analysis)

For visualization of retained PC2-associated genes, genes were ordered according to their corrected PC2 correlation coefficient, with retained positive genes ranked from highest to lowest positive correlation and retained negative genes ranked from strongest to weakest negative correlation. Samples were ordered according to either raw PC2 coordinates or study-corrected PC2 coordinates. Expression values were row-wise z-score normalized across samples prior to heatmap visualization.

### Single-nucleus RNA-seq analysis of leptomeningeal and cortical Wnt signaling programs (snRNA-seq)

A previously generated WT leptomeningeal single-nucleus RNA-seq dataset was used to define major leptomeningeal cell populations, including endothelial cells, macrophages, pia-associated fibroblasts, arachnoid-associated fibroblasts, dural border cells, and arachnoid barrier cells. Cells were visualized using Uniform Manifold Approximation and Projection (UMAP) and annotated based on established marker genes. Cell-type assignments were validated using curated marker-gene dot plots.

To provide a cortical comparison dataset, an annotated cortical single-cell RNA-seq atlas containing endothelial, mural, glial, epithelial, and neuronal populations was analyzed using an identical workflow. UMAP visualizations and curated marker-gene dot plots were generated to confirm cell-type identities.

Expression of Wnt pathway components was surveyed across both datasets using Seurat FeaturePlot and DotPlot functions. Genes examined included Wnt ligands (Wnt1–Wnt16 and Ndp), Frizzled receptors (Fzd1–Fzd10), canonical BBB Wnt signaling co-receptors and pathway components (Lrp5, Lrp6, Gpr124, Reck, Tspan12, Ctnnb1, Axin2, Lef1, and Tcf7), and canonical BBB-associated Wnt target genes (Apcdd1, Nkd1, Notum, and Spock2). Feature plots were generated using standardized expression limits (0–2 normalized expression units) to facilitate comparisons across genes and cell populations.

For endothelial-specific analyses, leptomeningeal and cortical endothelial cells were subset from their respective datasets and visualized using standardized dot plots. Dot size represents the percentage of expressing cells, whereas color intensity represents average normalized expression.

To independently evaluate Wnt ligand expression across CNS cell classes, publicly available Allen Brain Cell Atlas data were analyzed. Rare vascular-associated populations (endothelial cells, mural cells, vascular leptomeningeal cells, and microglia/perivascular macrophages) were retained in full, whereas abundant neuronal subclasses were downsampled to improve computational tractability while preserving cellular diversity. Expression matrices were queried for Wnt ligands, Frizzled receptors, BBB-associated Wnt signaling genes, and canonical BBB markers. UMAP visualizations and summary dot plots were generated using standardized expression scaling.

### Integration and analysis of cortical and leptomeningeal endothelial cells (snRNA-seq)

Single-nucleus RNA-seq datasets generated from cortical and leptomeningeal tissues were integrated using Seurat. Endothelial cells were identified based on expression of canonical endothelial markers including *Pecam1, Cldn5, Flt1,* and *Kdr* and subset for downstream analysis. Integrated endothelial nuclei were clustered using graph-based clustering and visualized using Uniform Manifold Approximation and Projection (UMAP).

Endothelial identity was validated by expression of established endothelial and blood–brain barrier markers and by the absence of enrichment for mural cell, fibroblast, neuronal, glial, and myeloid marker genes. Classical vascular zonation markers representing arterial, capillary, and venous endothelial populations were examined to assess the relationship between endothelial origin and vascular subtype identity.

### Quality control and doublet identification (snRNA-seq)

Standard quality-control metrics, including total transcript counts (nCount_RNA), detected genes (nFeature_RNA), and mitochondrial transcript percentage (percent.mt), were calculated for all endothelial nuclei. Doublets were identified using scDblFinder and visualized across clusters and on the integrated UMAP embedding. Sample-to-sample relationships were assessed using pseudobulk variance-stabilized expression profiles generated with DESeq2. Pairwise sample distances were calculated from variance-stabilized transformed (VST) expression values and displayed as a distance heatmap.

### Pseudobulk differential expression analysis (snRNA-seq)

To compare endothelial transcriptional programs across anatomical origins, endothelial nuclei were aggregated into pseudobulk samples according to biological replicate and tissue origin. Differential expression analysis was performed using DESeq2. The design formula included sample origin as the primary variable while accounting for variation in the number of nuclei contributing to each pseudobulk sample.

Genes were ranked according to DESeq2 test statistics. Differentially expressed genes were identified using an adjusted *P* value threshold of 0.05 and a minimum absolute log2 fold change of 0.5. Volcano plots and heatmaps were generated from DESeq2 results. For heatmap visualization, variance-stabilized expression values were row-scaled (Z-score transformed) to highlight relative expression differences between cortical and leptomeningeal samples. Prior to heatmap generation, ribosomal genes, mitochondrial genes, predicted genes (Gm), and RIKEN transcripts were excluded from candidate marker lists.

### Gene ontology enrichment analysis (snRNA-seq)

Gene Set Enrichment Analysis (GSEA) was performed using clusterProfiler with Gene Ontology Biological Process annotations. Genes were ranked according to DESeq2 statistics from the cortex-versus-leptomeninges comparison. Normalized enrichment scores (NES) were used to identify pathways enriched in either cortical or leptomeningeal endothelial populations.

### Module score analysis (snRNA-seq)

Curated endothelial functional modules were generated to summarize biological programs identified from the cortex versus leptomeninges pseudobulk differential expression and pathway analyses. Modules were intentionally defined as targeted, biologically interpretable gene sets rather than unbiased pathway discovery sets.

Module scores were calculated from pseudobulk variance-stabilized transformed (VST) expression values generated with DESeq2. For each module, genes present in the pseudobulk expression matrix were extracted, expression values were scaled by gene across samples, and the module score for each sample was calculated as the mean row-scaled VST expression of all detected genes in that module. Module scores were compared between cortical and leptomeningeal endothelial pseudobulk samples using Wilcoxon rank-sum tests, with Benjamini–Hochberg adjustment applied across modules.

The curated modules were defined as follows: (1) BBB transporters: Mfsd2a, Slc2a1, Abcb1a, Abcb1b, Abcg2, Slco1a4, Slc7a5, Lrp1, Lrp8, Slc22a27. (2) Wnt-BBB specification: Gpr124, Reck, Fzd4, Lef1, Apcdd1, Foxf2, Ctnnb1, Axin2, Tcf7l2, Wnt7a, Wnt7b. (3) Immune-related/interface: Ly96, Csf1, Lyn, Lgals9, Cd200, Tmem176b, Ifitm2, Mafb, S100a8, Lcp1, Tox, Klf10, Tgm2. (4) Endothelial adhesion: Icam1, Icam2, Vcam1, Sele, Selp, Itga2, Itga5, Mmrn1, Myzap, Adgrg6, Nectin2, Tspan18, Fblim1. (5) Extracellular matrix remodeling: Sparc, Fn1, Dcn, Col1a1, Col1a2, Col3a1, Col18a1, Bgn, Mmp14, Mmp28, Adamts9, Igfbp4. (6) Vesicular transport/transcytosis: Plvap, Cav1, Cavin1, Cavin2, Dnm2, Rab5a, Rab7a, Ehd2, Pacsin2. (7) Neurovascular interaction: Grin2b, Nlgn1, Cntnap2, Lrrtm4, Dlgap1, Cntn1, Grid2, Astn2, Gabrb3, Cacna2d3, Dpp10, Camk4. Representative genes from selected modules were visualized on the integrated endothelial UMAP using FeaturePlot.

### Pericyte and Astrocyte Coverage Analysis

Leptomeningeal and cortical vascular coverage analyses were performed on confocal Z-stack images acquired from neonatal mouse brain flatmount preparations stained for Cldn5 together with PDGFRβ, MCAM, Desmin, αSMA, GFAP, or Aqp4. For each image, the subarachnoid space (SAS) and cortical/glial limitans compartments were manually defined by selecting Z-ranges corresponding to the superficial leptomeningeal vasculature or underlying cortical vessels, respectively. Selected Z-ranges were duplicated and converted into maximum-intensity projections in Fiji/ImageJ. Images were Gaussian blurred (σ = 2), converted to 8-bit, and thresholded using marker– and layer-specific intensity thresholds to generate binary masks for Cldn5 and the corresponding marker channel. Overlap masks were generated using logical AND operations between Cldn5 and marker masks. Quantification was performed using a fixed ROI set (200 × 200 µm regions) applied identically across all images. For each ROI, raw fluorescence intensity, total mask area, and overlap area were measured for Cldn5 and the corresponding marker. Marker coverage was calculated as overlap area normalized to total Cldn5-positive area within each ROI. ROI-level measurements were analyzed in R using ggplot2 and ggpubr. Outliers were removed independently within each group using the 1.5× interquartile range method, and statistical comparisons between SAS and cortical compartments were performed using two-sided Wilcoxon rank-sum tests.

### Immunofluorescence Imaging and Quantification of ICAM1 Vascular Coverage

Flat-mount leptomeningeal and superficial cortical preparations were stained for Claudin-5 (Cldn5) and ICAM1 and imaged by confocal microscopy using identical acquisition settings across experimental groups. Z-stacks were collected through the leptomeningeal and superficial cortical vascular compartments.

For quantitative analysis, Cldn5 and ICAM1 channels were separated and processed independently in Fiji/ImageJ. Anatomical Z-ranges corresponding to the subarachnoid space (SAS/leptomeninges) and superficial cortex were manually annotated for each image stack based on vascular morphology and tissue depth. Maximum-intensity projections were generated separately for leptomeningeal and cortical compartments for both channels. Projected Z-depths were comparable across experimental groups and typically ranged from ∼15–35 µm total projected depth (data not shown).

Binary masks were generated following Gaussian smoothing and intensity thresholding of the Cldn5 and ICAM1 channels. Overlap masks were generated using logical AND operations between thresholded Cldn5 and ICAM1 masks. Quantification was performed using a fixed ROI set consisting of 100 × 100 µm regions sampled across each image. For each ROI, total Cldn5-positive area, ICAM1-positive area, and overlap area were measured. ICAM1 vascular coverage was calculated as:

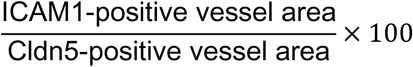

ROI-level analyses were used to assess local spatial heterogeneity in vascular activation, whereas mouse-level analyses were generated by averaging all ROIs from each animal prior to statistical comparison. Additional control analyses demonstrated that the observed increase in leptomeningeal ICAM1 coverage was not solely explained by differences in total Cldn5-positive vessel area. Outliers were identified independently within each experimental group using the 1.5× interquartile range (IQR) rule and excluded prior to plotting and statistical analysis.

### Quantification of Extravascular Biotin Leakage

For vascular leak experiments, neonatal mice were intravenously injected with sulfo-NHS-biotin tracer 30 minutes prior to tissue collection. Flat-mount leptomeningeal and superficial cortical preparations were stained for Cldn5 and biotin tracer and imaged using identical confocal acquisition settings across experimental groups.

Image stacks were processed identically to the ICAM1 analysis workflow. Separate maximum-intensity projections corresponding to leptomeningeal and cortical compartments were generated for the Cldn5 and biotin channels. Binary masks were generated following Gaussian smoothing and intensity thresholding. Cldn5 masks were used to define vascular regions, and inverse Cldn5 masks were generated to define non-vascular tissue space within each ROI. Biotin-positive signal located outside the Cldn5-positive vascular mask was identified using logical AND operations between the biotin mask and inverse Cldn5 mask. Projected Z-depths were comparable across experimental groups and generally ranged from ∼15–35 µm total projected depth (data not shown).

Quantification was performed using fixed 100 × 100 µm ROIs sampled across each image. For each ROI, total Cldn5-positive vessel area, total biotin-positive area, and extravascular biotin-positive area were measured. Extravascular leak was quantified both as absolute extravascular biotin-positive area per ROI and as normalized leak relative to available non-vascular tissue space:

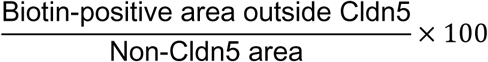

where non-Cldn5 area was defined as the total ROI area minus the Cldn5-positive vessel area. ROI-level analyses were used to assess local heterogeneity in vascular permeability, whereas mouse-level analyses were generated by averaging all ROIs from each animal prior to statistical comparison. Additional control analyses demonstrated that the observed increase in leptomeningeal tracer leakage was not solely explained by differences in total Cldn5-positive vessel area, total biotin signal, or available non-vascular tissue area. Outliers were identified independently within each experimental group using the 1.5× IQR rule and excluded prior to plotting and statistical analysis.

### Quantification of Laminin Remodeling and Fragmentation

Flat-mount leptomeningeal preparations were stained for Cldn5 and Laminin and imaged by confocal microscopy using identical acquisition settings across experimental groups. Maximum-intensity projections corresponding to the leptomeningeal vascular compartment were generated following manual annotation of anatomical Z-ranges based on tissue morphology and vascular organization.

For quantitative analysis, laminin images were processed in Fiji/ImageJ following background subtraction and Gaussian smoothing. Fixed intensity thresholding was applied to generate binary laminin masks. Quantification was performed using fixed 100 × 100 µm ROIs sampled across each image.

Total laminin-positive area and mean laminin intensity were measured within each ROI. To quantify laminin fragmentation, contiguous laminin-positive objects were identified using Fiji/ImageJ particle analysis on thresholded laminin masks. The number of laminin-positive objects within each ROI was quantified as a measure of structural discontinuity and fragmentation of the vascular basement membrane.

ROI-level analyses were used to assess local heterogeneity in laminin organization, whereas mouse-level analyses were generated by averaging all ROIs from each animal prior to statistical comparison. Outliers were identified independently within each experimental group using the 1.5× IQR rule and excluded prior to plotting and statistical analysis.

### Comparison of human leptomeningeal and blood–brain barrier endothelial cells (Human snRNA-seq)

Published human parenchymal brain vascular single-nucleus RNA-sequencing data were obtained from Sun et al. (2023)^31^, who profiled the cerebrovasculature across six brain regions from ROSMAP donors. Human leptomeningeal single-nucleus RNA-sequencing data were obtained from Kearns et al. (2023), who profiled postmortem leptomeninges from aged ROSMAP participants. Both datasets were reanalyzed in R using Seurat. For the parenchymal dataset, nuclei carrying the published endothelial annotation were extracted from the original count matrix and metadata to generate an endothelial-only reference. Published endothelial subtype annotations were retained, comprising arterial endothelial (aEndo), capillary endothelial (capEndo), and venous endothelial (vEndo) populations. For the leptomeningeal dataset, published endothelial annotations were retained, comprising aEndo, capillary, vEndo_1, and vEndo_2 populations. Thus, vascular identities assigned independently in the original studies were preserved without reclustering the datasets together or transferring subtype labels between datasets.

Because both datasets included aged individuals with and without Alzheimer’s disease (AD), the cross-compartment analyses shown in Fig. 7 were restricted to donors without a clinical AD diagnosis to minimize disease-associated transcriptional remodeling as a source of variation. Parenchymal donors annotated as non-AD were retained, while leptomeningeal donors classified as no cognitive impairment (NCI) or mild cognitive impairment (MCI) were grouped as non-AD; donors classified as AD were excluded. Donor identifiers, sex annotations, age information, and clinical classifications were retained from the source metadata and harmonized before subsetting. The resulting comparison therefore used aged non-AD endothelial populations derived from the broader ROSMAP resource while preserving the independently generated anatomical and transcriptional annotations of each dataset.

RNA expression was independently normalized within each dataset using Seurat NormalizeData with LogNormalize and a scale factor of 10,000, and downstream comparisons were restricted to genes represented in both datasets. Endothelial populations were compared within matched vascular classes rather than across the complete endothelial populations: parenchymal aEndo versus leptomeningeal aEndo, parenchymal capEndo versus leptomeningeal capillary cells, and parenchymal vEndo independently versus leptomeningeal vEndo_1 and vEndo_2 cells. The datasets were not integrated or batch-corrected before these comparisons, thereby avoiding forced alignment of transcriptionally distinct endothelial states across anatomical compartments.

Endothelial population structure was visualized independently within each dataset using UMAP. For the parenchymal endothelial reference, variable features were identified using the variance-stabilizing transformation method, followed by data scaling, principal-component analysis, and UMAP using the first 30 principal components. The published leptomeningeal endothelial UMAP and subtype annotations were retained. Correspondence of endothelial identities between datasets was assessed using the canonical endothelial genes *PECAM1, VWF, KDR, FLT1,* and *CDH5* and barrier-associated genes *CLDN5* and *SLC2A1*. For this descriptive comparison, leptomeningeal vEndo_1 and vEndo_2 nuclei were combined into a single venous category to permit direct arterial–capillary–venous visualization between datasets.

To quantify compartment-associated transcriptional specialization, expression prevalence was calculated from raw counts for each shared gene as the percentage of nuclei containing at least one detected transcript. Compartment bias was defined as the percentage of LBCSFB nuclei expressing a gene minus the percentage of matched BBB nuclei expressing that gene. Genes were ranked by this prevalence difference to visualize the distribution of BBB– and LBCSFB-biased transcripts within each vascular class. Mitochondrial genes and incompletely annotated transcripts beginning with AC, AL, AP, or LINC were excluded from ranked visualization and subsequent concordance analyses.

As a complementary measure of expression magnitude, cell-level differential-expression analysis was performed independently for each matched vascular comparison using MAST implemented through Seurat FindMarkers. Raw counts from the relevant BBB and LBCSFB populations were combined and renormalized using LogNormalize with a scale factor of 10,000. LBCSFB was specified as the numerator and BBB as the reference, such that positive average log2 fold changes indicate higher expression in LBCSFB endothelial cells. The number of detected genes per nucleus (nFeature_RNA) was included as a latent variable, and no minimum expression-frequency or fold-change threshold was imposed during testing. Benjamini–Hochberg-adjusted *P* values were calculated across tested genes. Because these analyses treated individual nuclei rather than donors as observations, MAST statistics were used as a complementary measure of transcriptional effect size and not as donor-level inference.

Genome-wide agreement between the two measures of compartment bias was assessed independently for each matched vascular population using Spearman rank correlation between the LBCSFB-minus-BBB expression-prevalence difference and MAST average log2 fold change. Correlations were positive for aEndo (ρ = 0.56), capillary (ρ = 0.42), vEndo_1 (ρ = 0.62), and vEndo_2 (ρ = 0.68) comparisons (all *P* < 2 × 10⁻¹⁶), indicating concordant compartment-associated differences across the two measures of transcript abundance.

To identify a recurrent LBCSFB-associated endothelial signature, genes were required within each matched comparison to exhibit a ≥10-percentage-point increase in expression prevalence in LBCSFB endothelial cells, a MAST average log2 fold change ≥1, and a Benjamini–Hochberg-adjusted *P* < 0.05. Genes satisfying all three criteria in at least three of the four vascular comparisons were considered recurrently LBCSFB-biased. Candidate genes were ranked first by the number of comparisons in which they met all criteria, then by median qualifying prevalence difference and median qualifying MAST effect size; the top 15 ranked genes were displayed. In dot plots, dot size represents the percentage of nuclei with detectable expression and color represents mean log-normalized expression among expressing nuclei. Expression of selected representative genes (*TJP2, SLC26A3, BCOR, PTGDS, ST3GAL3,* and *SOD2*) was additionally visualized across the complete BBB and LBCSFB endothelial UMAPs using a common log-normalized expression scale of 0–4, with values >4 saturated at the upper limit.

### Reanalysis of human leptomeningeal endothelial cells (Human snRNA-seq)

Human leptomeningeal single-nucleus RNA-sequencing data were obtained from Kearns et al. (2023) through Synapse under the applicable data-use terms and analyzed in R using Seurat and DESeq2^13^. The published endothelial annotations were retained, comprising arterial endothelial (aEndo), capillary, vEndo_1, and vEndo_2 populations. The endothelial dataset contained 16,525 nuclei derived from 18 biological donors. Donor metadata provided with the dataset were matched to individual nuclei using the ROSMAP project identifier and published clinical and sex annotations were retained for downstream analyses. Clinical diagnoses were grouped as NCI/MCI or Alzheimer’s disease (AD) for disease-associated comparisons.

For differential-expression analyses, raw transcript counts were aggregated at the donor level within each endothelial population to generate donor × endothelial-subtype pseudobulk profiles, thereby treating biological donors rather than individual nuclei as the unit of replication. Differential expression analysis between AD and NCI/MCI donors was performed using DESeq2. Analyses were performed across the complete donor cohort and independently within female and male donors for each endothelial population. Genes displayed in subtype-level heatmaps were selected using a nominal DESeq2 Wald-test *P* < 0.01 and absolute log2 fold change ≥ 0.5; Benjamini–Hochberg-adjusted *P* values were additionally calculated to identify genes meeting a false-discovery rate (FDR) < 0.05. Heatmaps display DESeq2 log2 fold changes for AD relative to NCI/MCI, with nominally significant and FDR-significant genes indicated separately.

Sex-stratified disease-associated transcriptional changes within the vEndo_1 population were further visualized using volcano plots generated from the corresponding female– and male-specific donor-level DESeq2 analyses. To directly compare the magnitude and direction of disease-associated transcriptional changes between sexes, female and male AD-versus-NCI/MCI log2 fold changes were compared for genes represented in the sex-stratified analyses. KYNU expression was examined at the individual-donor level using DESeq2-normalized pseudobulk counts from vEndo_1 endothelial cells, with donors stratified by sex and clinical diagnosis. Statistical significance and effect sizes for KYNU were derived from the corresponding sex-specific DESeq2 models.

## Figure Legends

**Extended Data Fig. 1:**
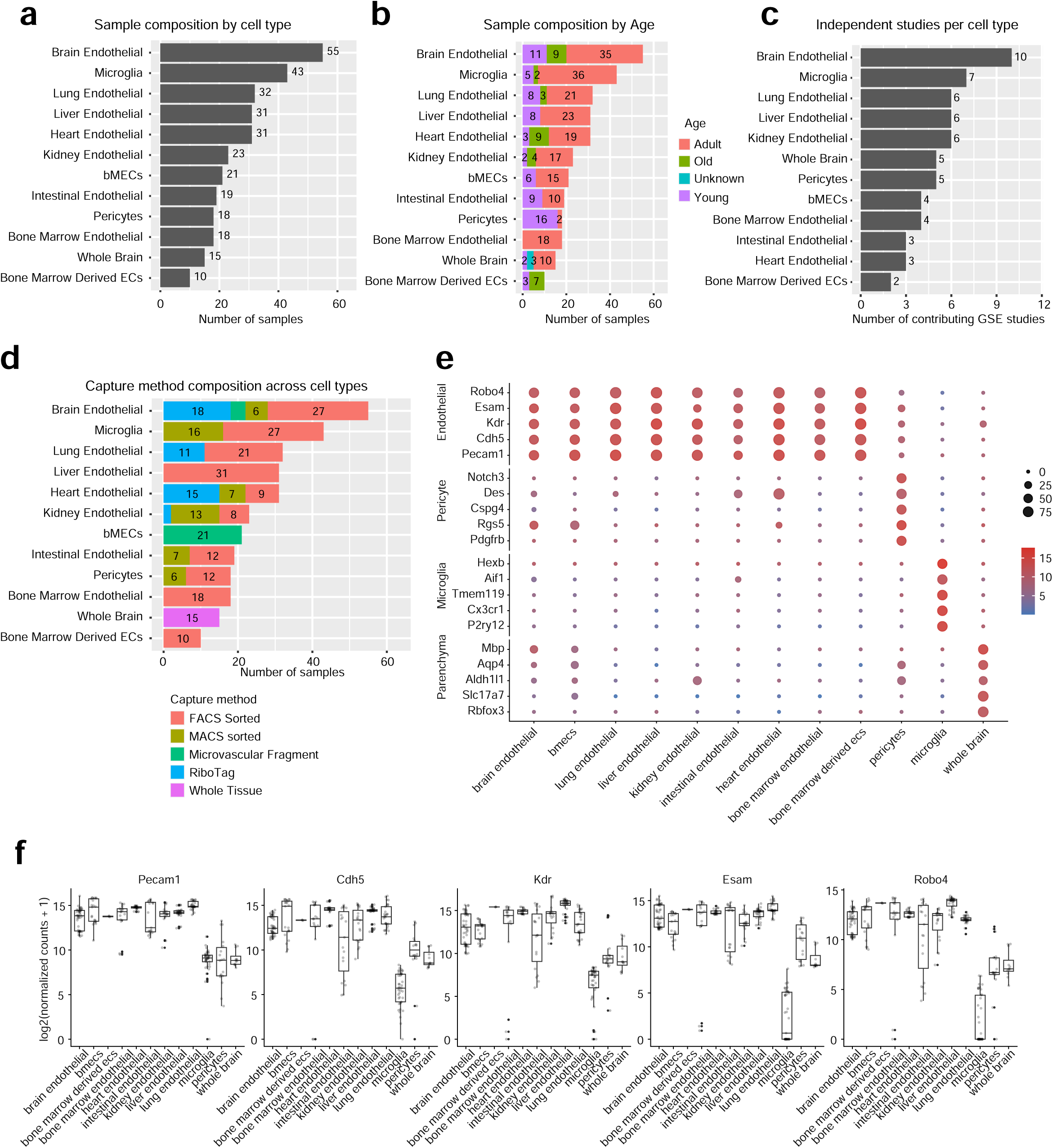
Metadata composition of Brain Endothelial RNA-seq Atlas and validation of sample purity. (a) Distribution of samples across major cell and tissue classes included in the atlas. (b) Age composition of samples within each cell class. Samples were classified as young, adult, old, or unknown age based on original study metadata. (c) Number of independent studies contributing samples to each cell class. (d) Distribution of sample isolation methodologies across cell classes, including fluorescence-activated cell sorting (FACS), magnetic-activated cell sorting (MACS), microvascular fragment isolation, RiboTag-based approaches, and whole-tissue sequencing. (e) Dot plot showing expression of canonical endothelial, pericyte, microglial, and parenchymal marker genes across sample groups. Dot color indicates mean log2(normalized counts + 1) expression and dot size indicates the percentage of studies exceeding the reference expression threshold defined for each marker. (f) Expression of canonical endothelial marker genes (*Pecam1, Cdh5, Kdr, Esam,* and *Robo4*) across sample groups. Boxes indicate the interquartile range with the median shown as the center line; whiskers extend to 1.5× the interquartile range and individual points represent individual RNA-seq samples.

**Extended Data Fig. 2:**
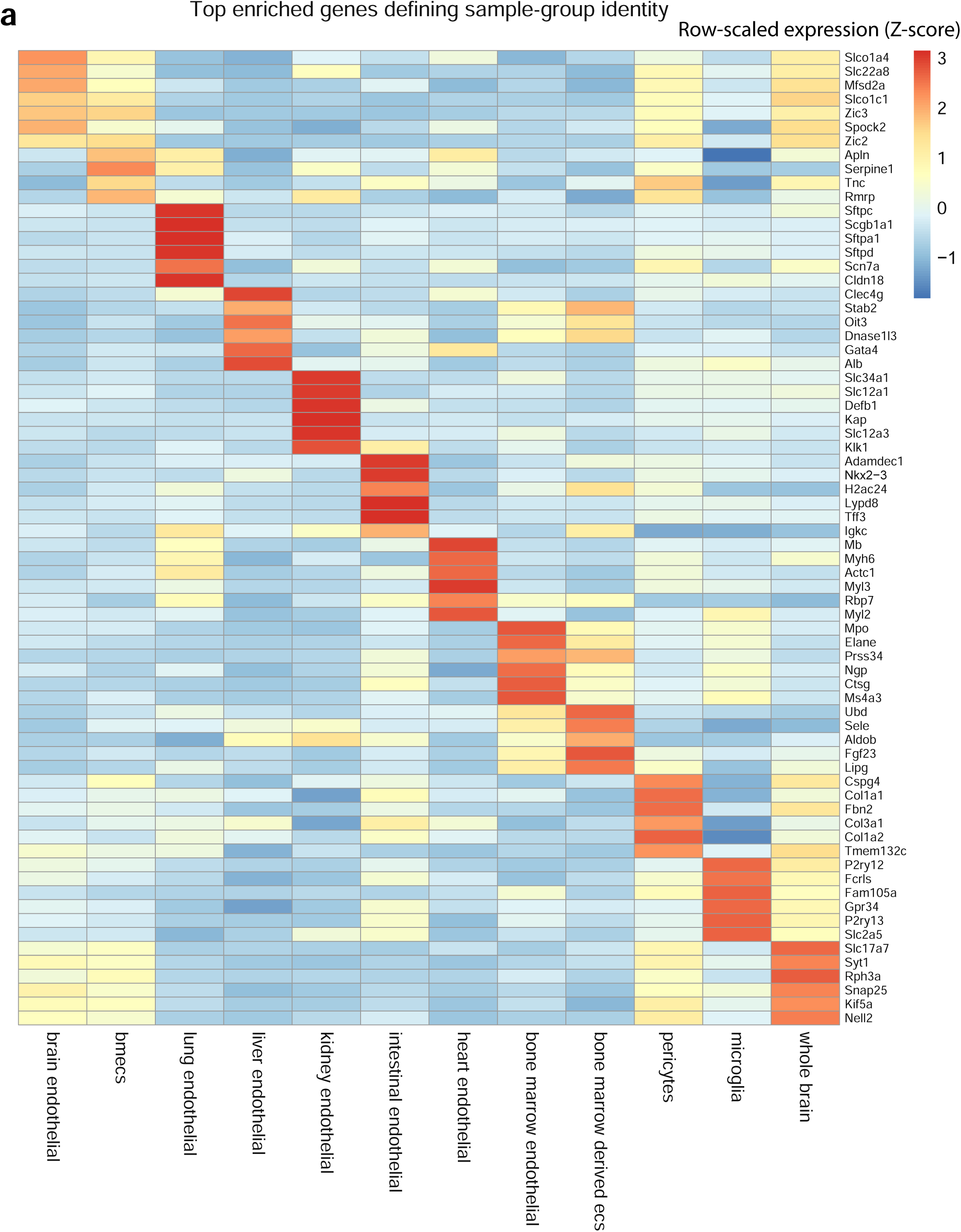
Top enriched transcriptional signatures across endothelial and CNS-associated sample groups. (a) Heatmap showing the five most enriched genes for each sample group within the cross-study atlas. Genes were ranked according to the difference between mean expression within the focal group and mean expression across all remaining groups. Heatmap colors represent row-scaled expression (z-scores).

**Extended Data Fig. 3:**
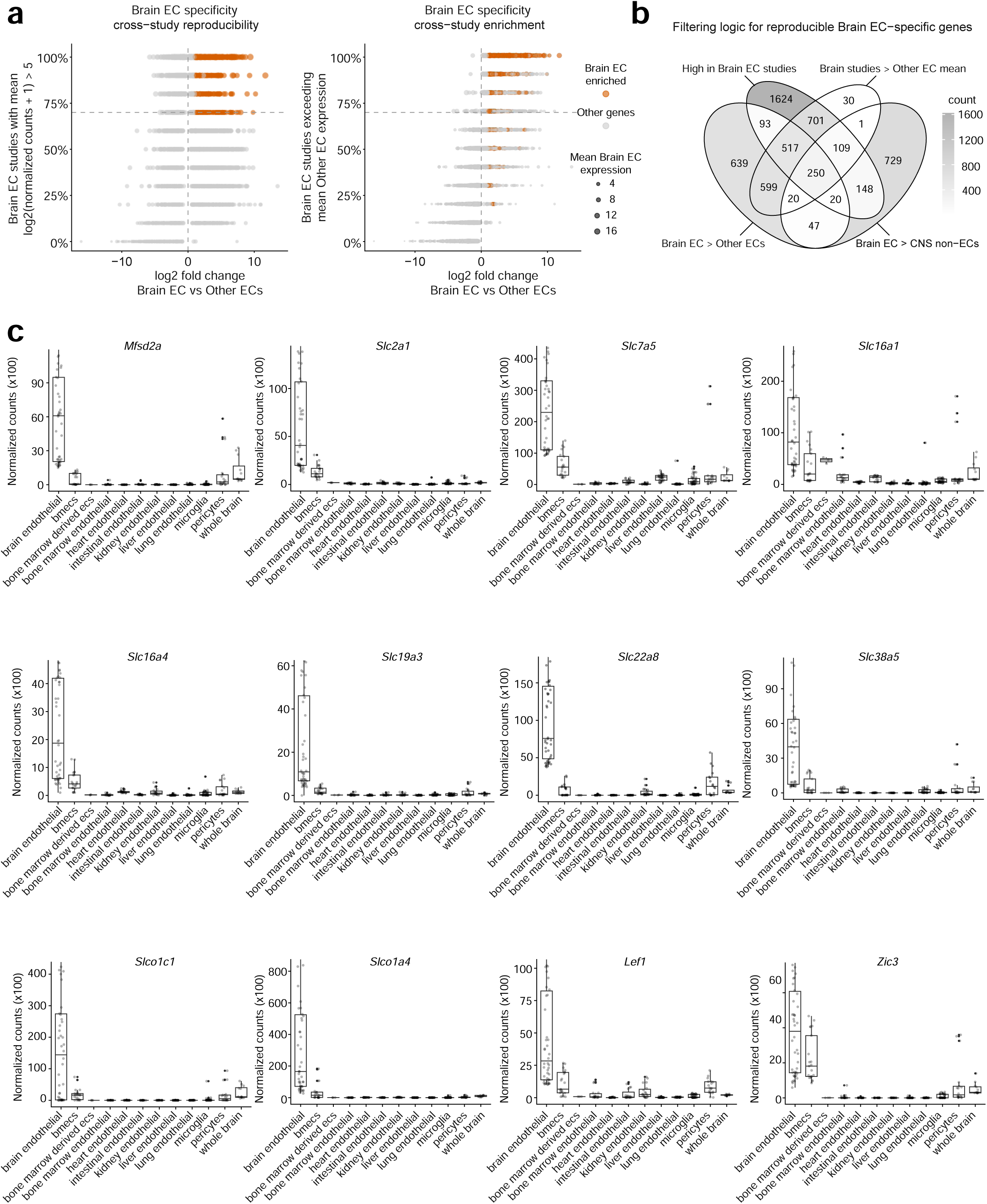
Sequential filtering for endothelial specificity, cross-study reproducibility, and CNS enrichment defines a robust Brain endothelial cell gene signature. (a) Cross-study reproducibility analysis of Brain EC-enriched genes. Left, genes are plotted by Brain EC enrichment relative to other endothelial populations and the fraction of independent Brain EC studies with mean log2(normalized counts + 1) ≥ 5. Right, genes are plotted by Brain EC enrichment and the fraction of Brain EC studies exceeding the mean expression observed in other endothelial samples. Brain EC-enriched genes are highlighted in orange. (b) Four-way Venn diagram showing the filtering strategy used to define reproducible Brain EC-specific genes. Candidate genes were filtered based on enrichment in Brain ECs relative to other endothelial populations, high expression across Brain EC studies, expression above the mean of other endothelial samples, and enrichment relative to CNS non-endothelial populations. The central intersection identifies 250 genes meeting all four criteria. (c) Normalized count distributions for representative Brain EC-enriched genes across endothelial and CNS-associated sample groups. Genes shown include canonical BBB-associated markers and candidate Brain EC-enriched genes identified through the filtering pipeline, including Mfsd2a, Slc2a1, Slc7a5, Slc16a1, Slc16a4, Slc19a3, Slc22a8, Slc38a5, Slco1c1, Slco1a4, Lef1, and Zic3. Boxes indicate the interquartile range with the median shown as the center line; whiskers extend to 1.5× the interquartile range and individual points represent individual RNA-seq samples.

**Extended Data Fig. 4:**
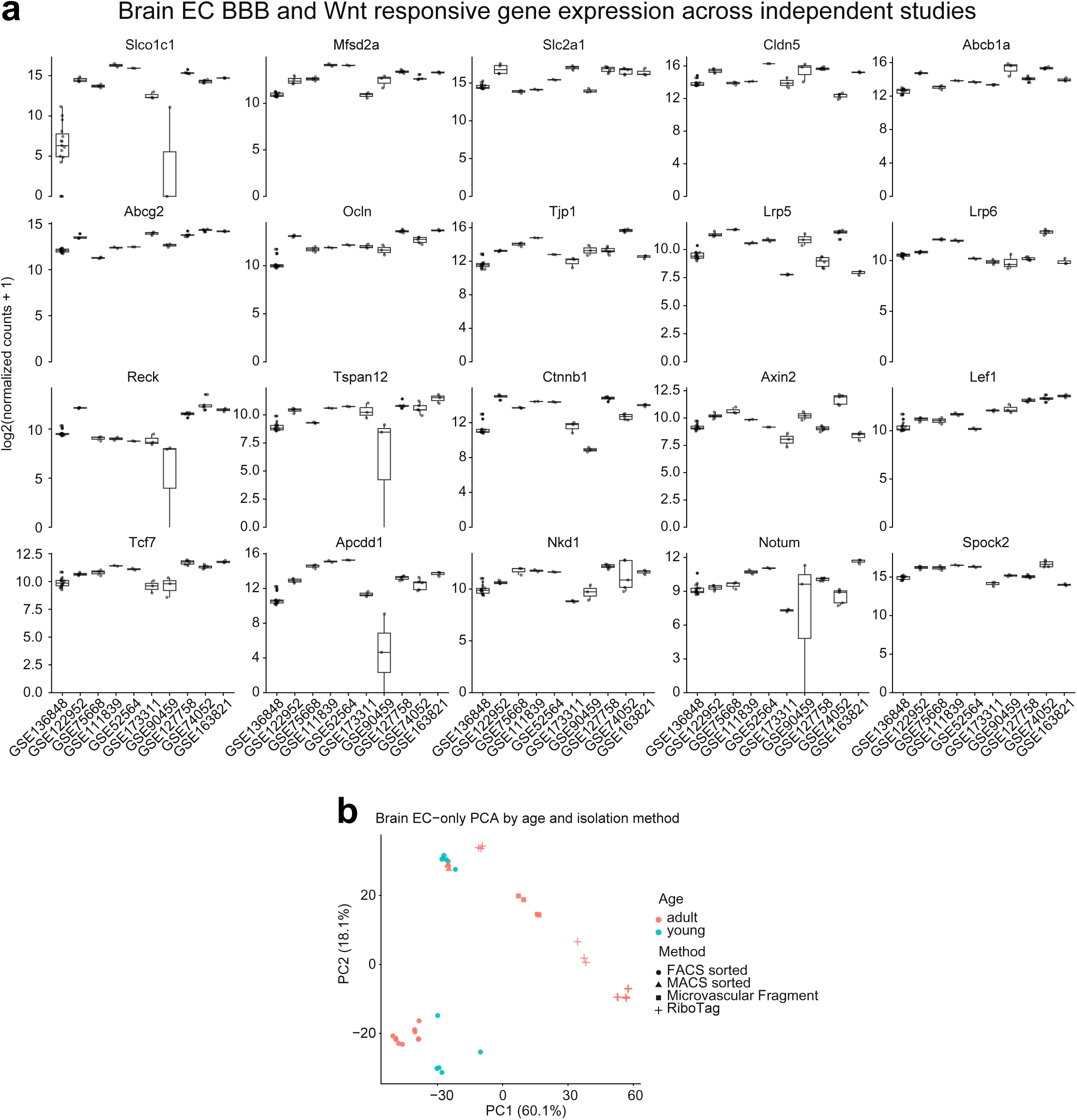
Independent Brain EC RNA-seq studies exhibit a conserved BBB signature despite substantial transcriptomic heterogeneity. (a) Expression of BBB-associated genes and Wnt-responsive components across independent Brain EC studies. Boxplots show log2(normalized counts + 1) expression for canonical BBB transporters and junctional genes, including Slco1c1, Mfsd2a, Slc2a1, Cldn5, Abcb1a, Abcg2, Ocln, and Tjp1, as well as BBB/Wnt-associated genes including Lrp5, Lrp6, Reck, Tspan12, Ctnnb1, Axin2, Lef1, Tcf7, Apcdd1, Nkd1, Notum, and Spock2. Boxes indicate the interquartile range with the median shown as the center line; whiskers extend to 1.5× the interquartile range and individual points represent individual RNA-seq samples. (b) Brain EC-only PCA colored by age and shaped by isolation method. Samples were generated using FACS sorting, MACS sorting, microvascular fragment isolation, or RiboTag-based approaches.

**Extended Data Fig. 5:**
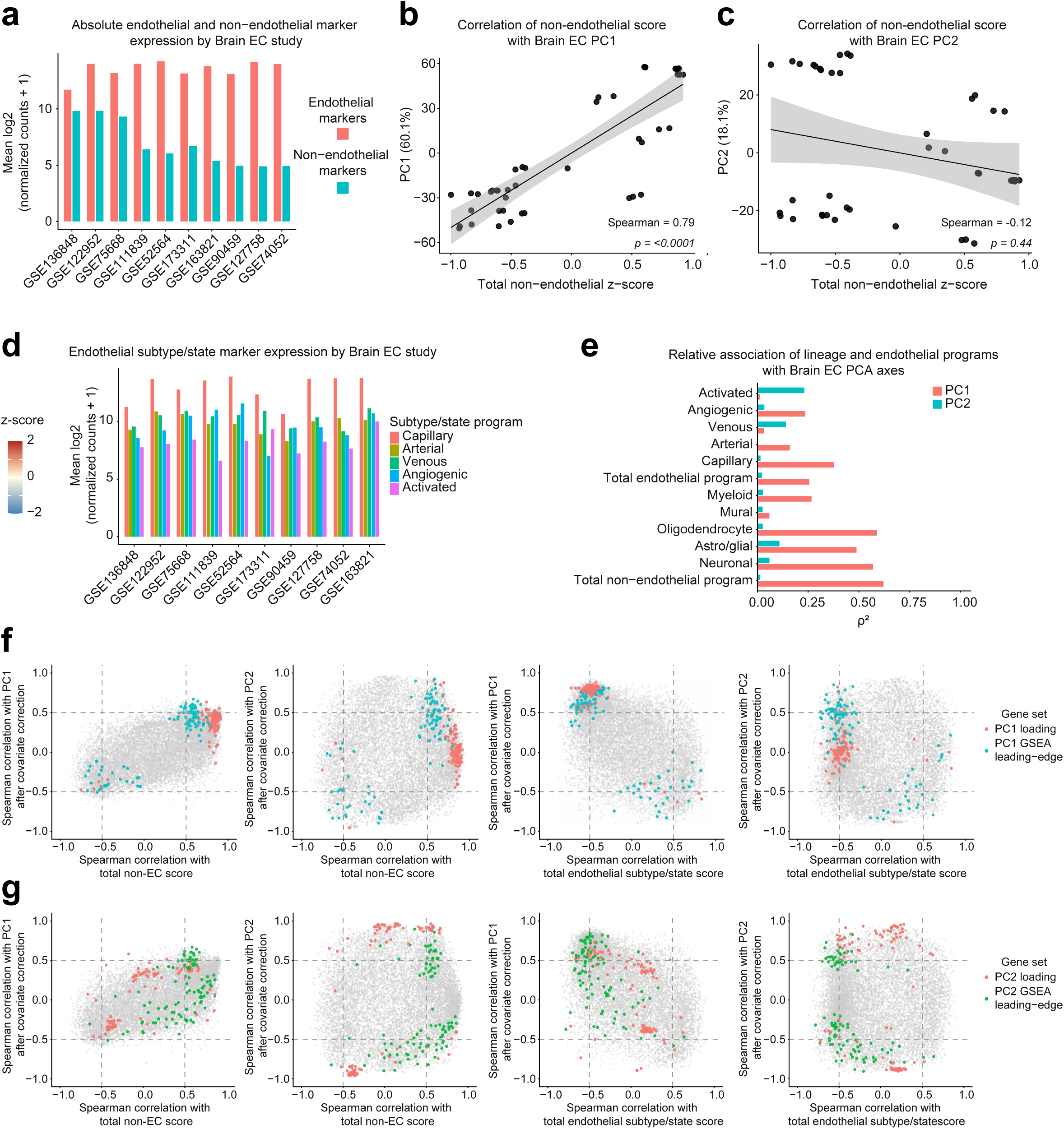
Brain endothelial heterogeneity is not completely explained by non-endothelial markers or endothelial subtype/state programs. (a) Mean expression of endothelial markers and non-endothelial markers across Brain EC studies. (b) Correlation between total non-endothelial score and Brain EC PC1. (c) Correlation between total non-endothelial score and Brain EC PC2. (d) Mean endothelial subtype/state marker expression across Brain EC studies, including capillary, arterial, venous, angiogenic, and activated programs. (e) Relative association of lineage and endothelial subtype/state programs with Brain EC PC1 and PC2. Bars show squared Spearman correlation coefficients. (f) Relationship between gene-level PC1 associations after covariate correction and correlation with total non-endothelial score or total endothelial program score. PC1 loading genes and PC1 GSEA leading-edge genes are highlighted. (g) Relationship between gene-level PC2 associations after covariate correction and correlation with total non-endothelial score or total endothelial program score. PC2 loading genes and PC2 GSEA leading-edge genes are highlighted.

**Extended Data Fig. 6:**
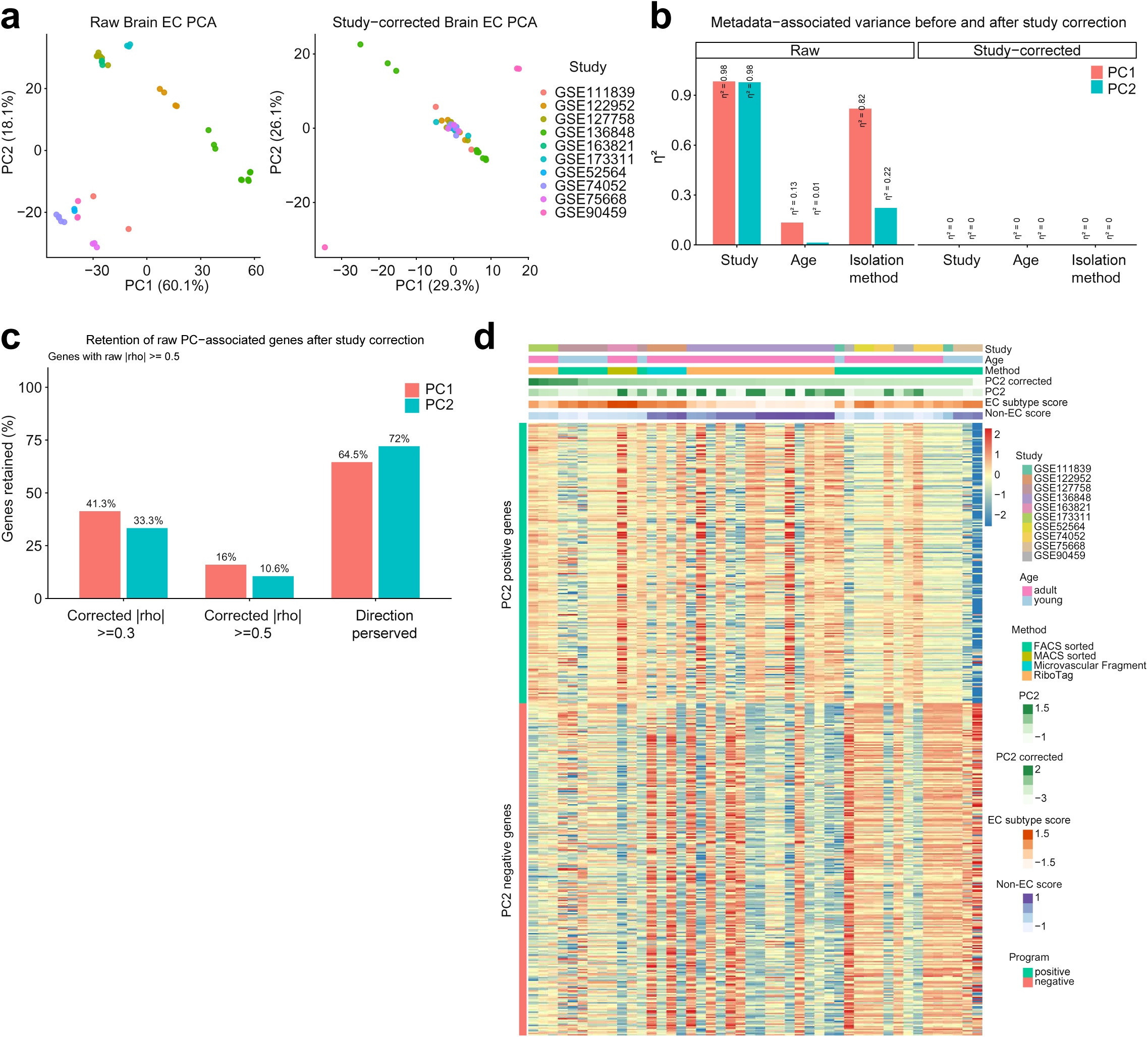
Study-associated variation contributes to Brain EC PCA structure, but gene-level transcriptional associations persist after correction. (a) Brain EC PCA before and after study correction. Samples are colored by GEO study identity. (b) Metadata-associated variance before and after study correction. Bars show η² values for the association of PC1 and PC2 with study identity, age, and isolation method. (c) Retention of raw PC-associated genes after study correction. Genes with raw |ρ| ≥ 0.5 were assessed for corrected |ρ| ≥ 0.3, corrected |ρ| ≥ 0.5, and preservation of association direction. (d) Heatmap of retained PC2-associated genes ordered by study-corrected PC2. Annotations indicate study, age, isolation method, raw PC2, corrected PC2, endothelial subtype/state score, non-endothelial score, and PC2 program membership.

**Extended Data Fig. 7:**
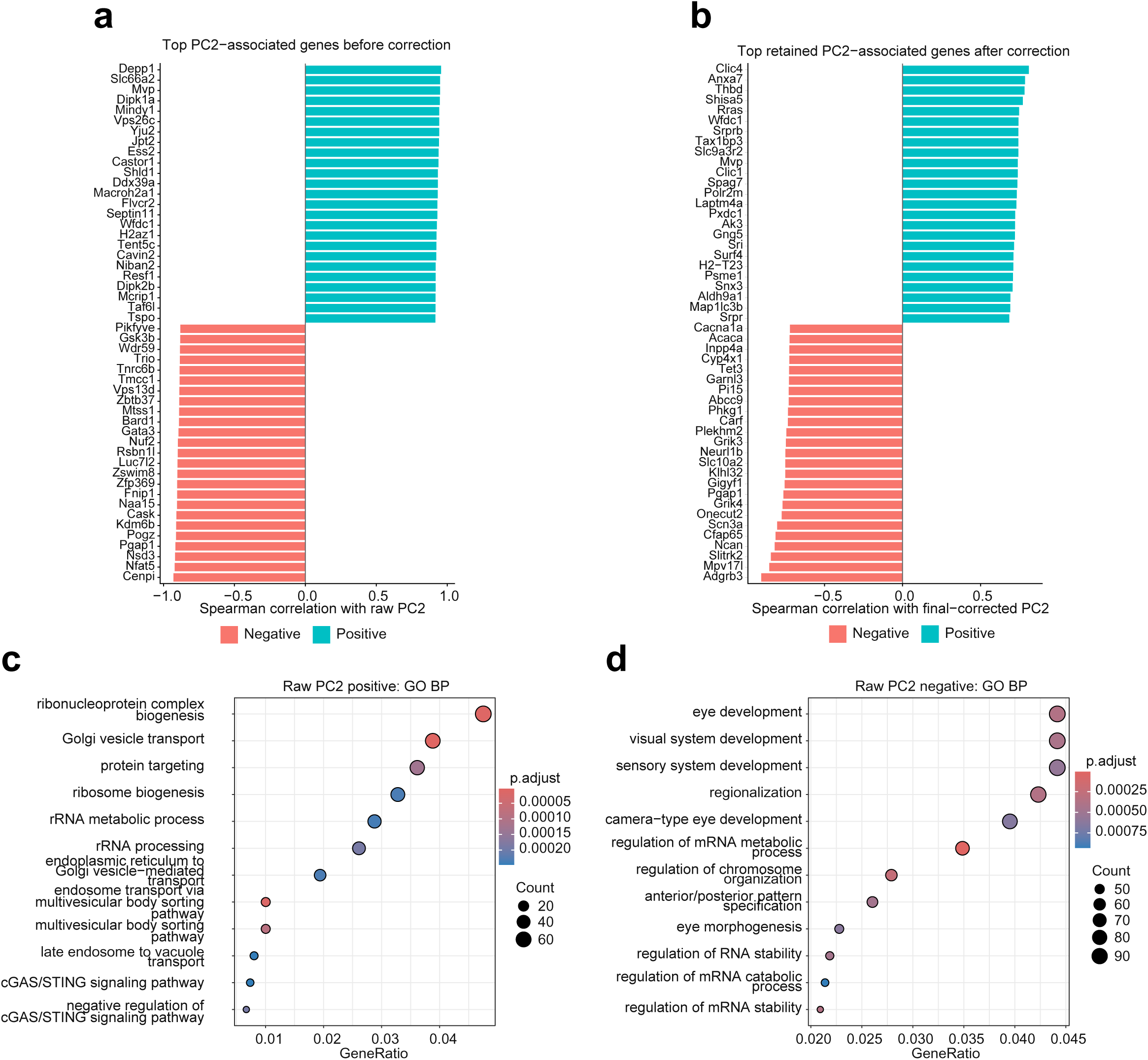
Residual transcriptomic heterogeneity among Brain EC studies is associated with a transcriptional continuum. (a) Top PC2-associated genes before correction, ranked by Spearman correlation with raw PC2. (b) Top retained PC2-associated genes after correction for study identity, non-endothelial score, and endothelial subtype/state score, ranked by Spearman correlation with final-corrected PC2. (c) Gene Ontology Biological Process enrichment of raw PC2-positive genes. GeneRatio denotes the proportion of genes from the corresponding PC2-associated gene program that are annotated to a given GO term (number of overlapping genes divided by the total number of genes submitted for enrichment analysis). Point size indicates the number of genes contributing to the GO term, and point color indicates the Benjamini–Hochberg adjusted *P* value. (d) Gene Ontology Biological Process enrichment of raw PC2-negative genes. GeneRatio denotes the proportion of genes from the corresponding PC2-associated gene program that are annotated to a given GO term (number of overlapping genes divided by the total number of genes submitted for enrichment analysis). Point size indicates the number of genes contributing to the GO term, and point color indicates the Benjamini–Hochberg adjusted *P* value.

**Extended Data Fig. 8:**
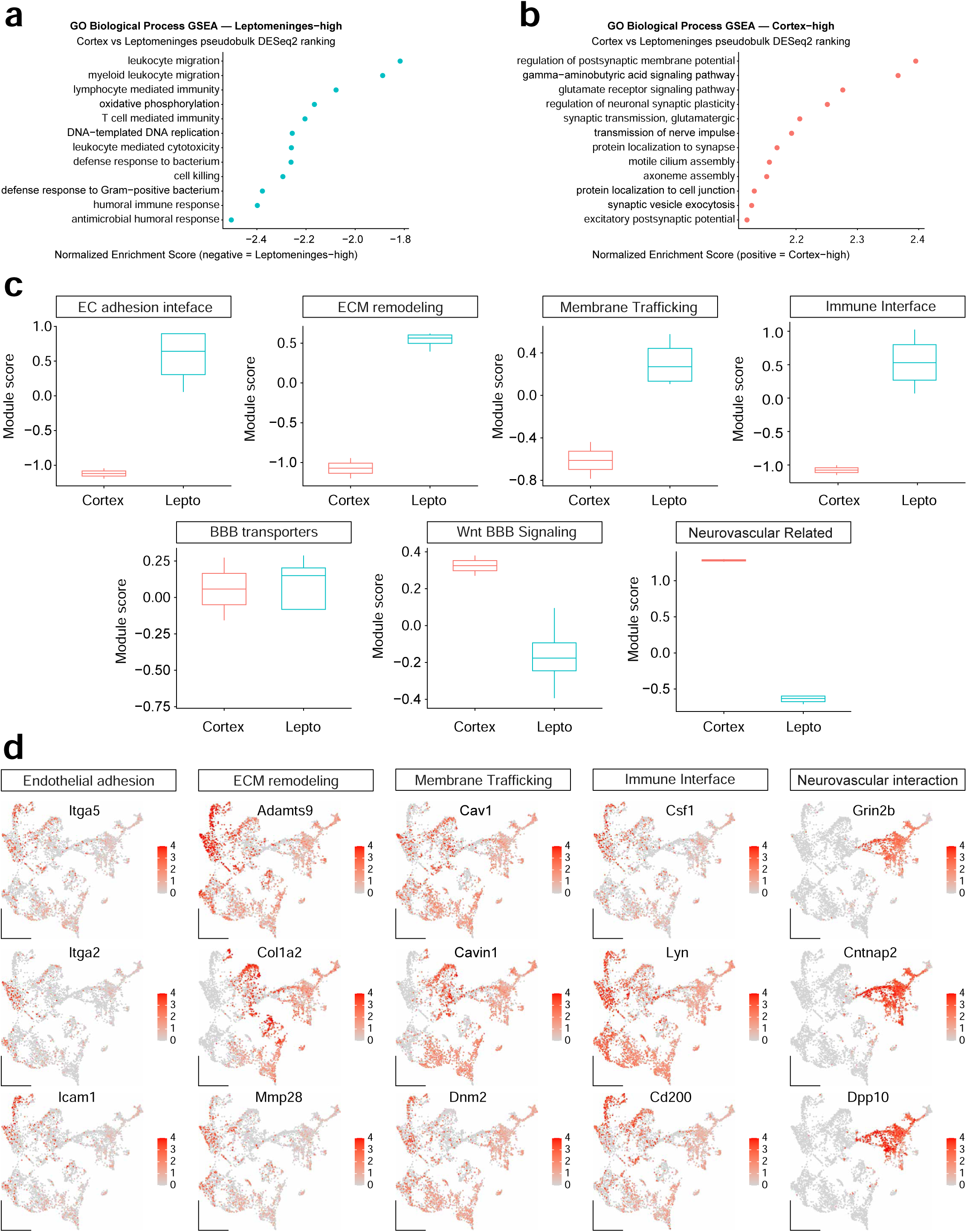
Functional pathway analysis reveals immune-interacting and neurovascular endothelial programs. (a) Gene ontology (GO) biological process gene set enrichment analysis (GSEA) of genes enriched in leptomeningeal endothelial cells. Enriched pathways include leukocyte migration, lymphocyte-mediated immunity, antimicrobial responses, and bacterial defense programs. (b) GO biological process GSEA of genes enriched in cortical endothelial cells. Enriched pathways include synaptic transmission, neuronal signaling, glutamate receptor signaling, and regulation of postsynaptic membrane potential. (c) Comparison of module scores between cortical and leptomeningeal endothelial cells. Modules represent endothelial adhesion, extracellular matrix remodeling, membrane trafficking, immune interface, BBB transporter, Wnt-BBB signaling, and neurovascular interaction programs. (d) Representative feature plots showing genes associated with endothelial adhesion (Itga5, Itga2, Icam1), extracellular matrix remodeling (Adamts9, Col1a2, Mmp28), membrane trafficking (Cav1, Cavin1, Dnm2), immune interface (Csf1, Lyn, Cd200), and neurovascular interaction (Grin2b, Cntnap2, Dpp10) programs.

**Extended Data Fig. 9:**
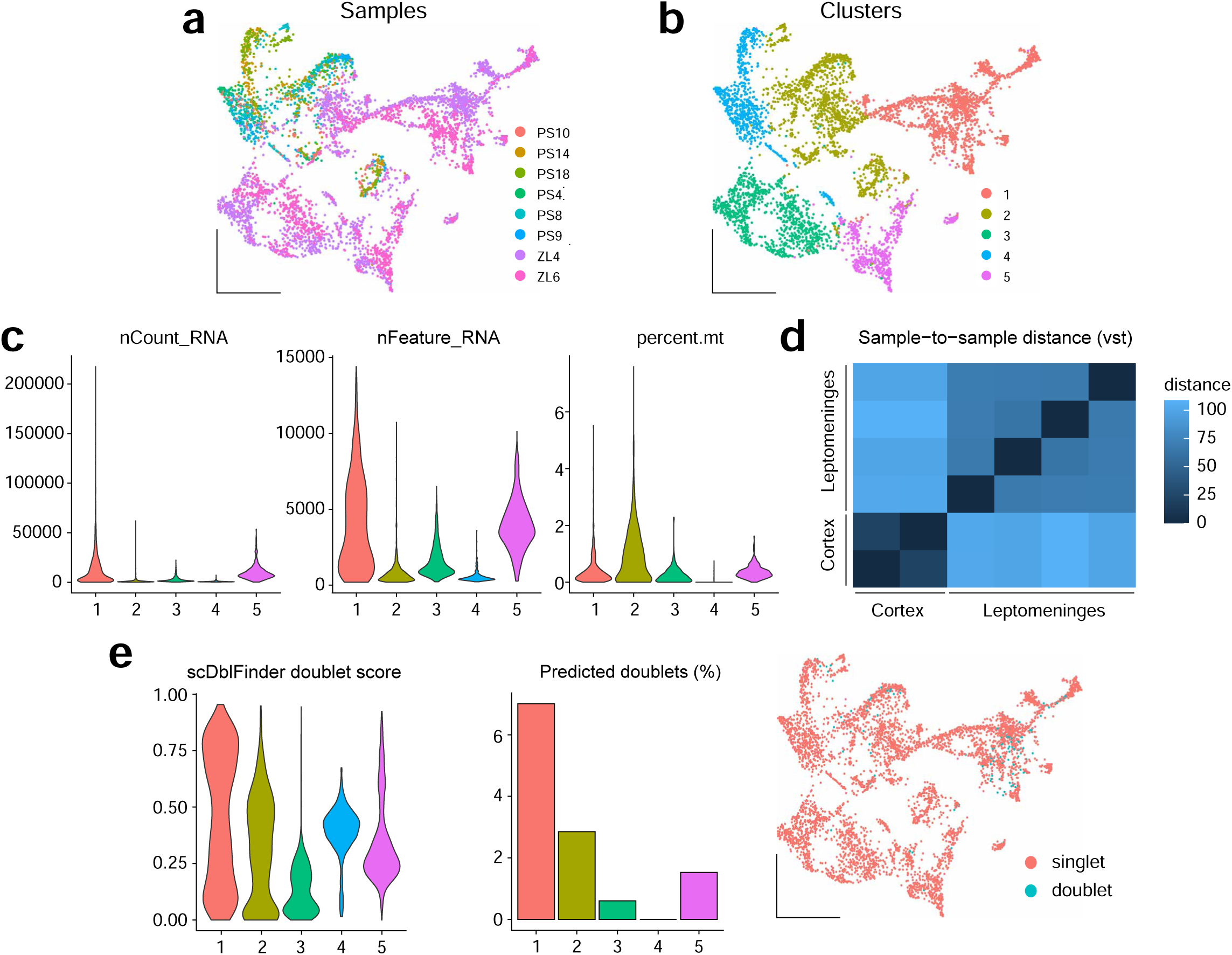
Quality control and integration of cortical and leptomeningeal endothelial nuclei. (a) UMAP of endothelial nuclei colored by sample identity, demonstrating contribution of multiple independent cortical and leptomeningeal samples to the integrated endothelial dataset. (b) UMAP of endothelial nuclei colored by unsupervised cluster assignment. (c) Violin plots showing quality-control metrics across endothelial clusters, including total RNA counts (nCount_RNA), detected genes (nFeature_RNA), and mitochondrial transcript percentage (percent.mt). (d) Sample-to-sample distance heatmap generated from variance-stabilized pseudobulk expression values, demonstrating segregation primarily by anatomical origin. (e) Doublet detection analysis using scDblFinder showing cluster-level doublet scores, predicted doublet percentages, and UMAP localization of predicted singlets and doublets.

**Extended Data Fig. 10:**
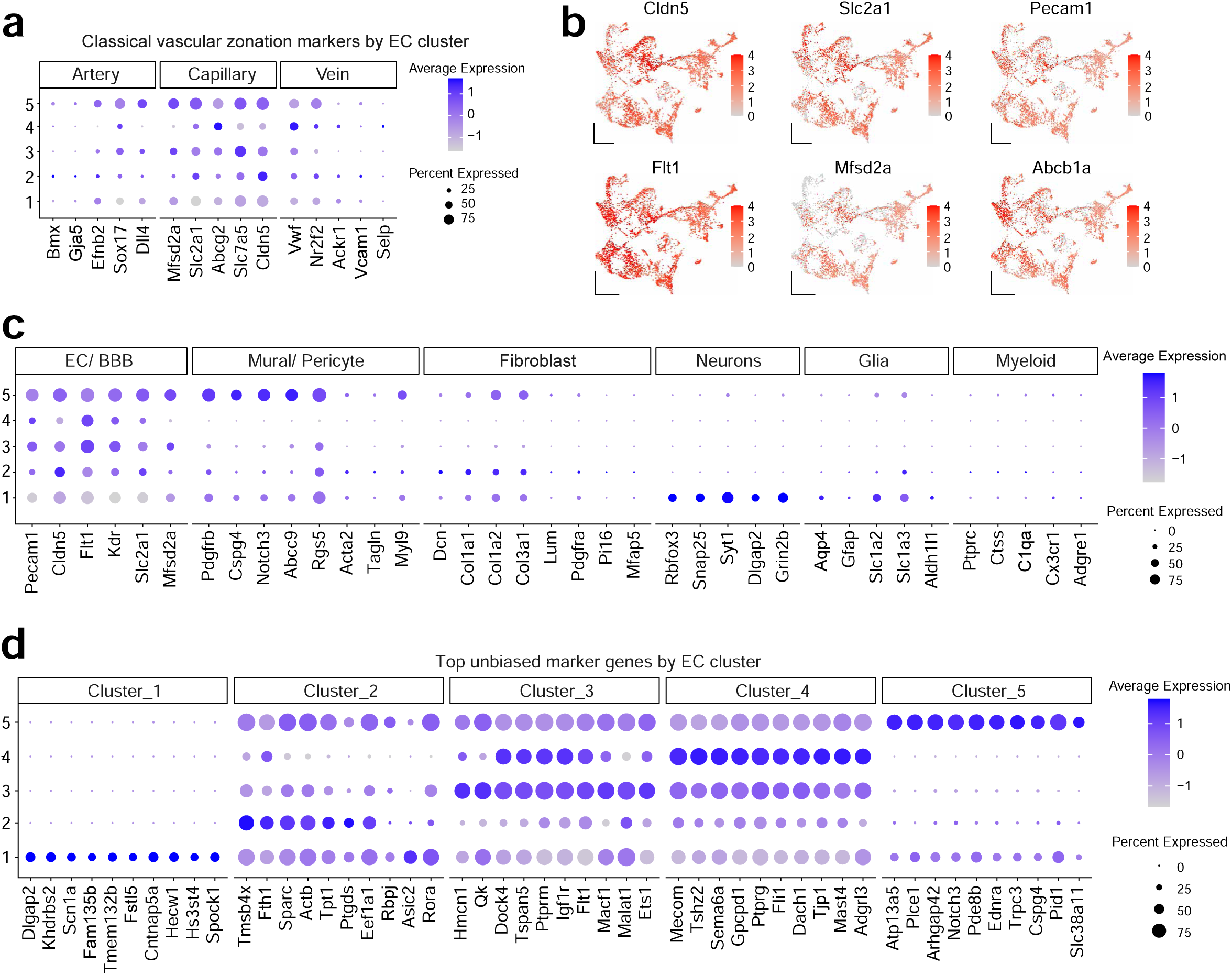
Validation of endothelial identity and cluster annotation. (a) Dot plot showing expression of classical arterial, capillary, and venous endothelial zonation markers across endothelial clusters. (b) Feature plots showing expression of canonical endothelial and BBB-associated markers including Cldn5, Slc2a1, Pecam1, Flt1, Mfsd2a, and Abcb1a. (c) Dot plot demonstrating enrichment of endothelial markers and minimal contamination by mural/pericyte, fibroblast, neuronal, glial, and myeloid gene signatures. (d) Top marker genes defining each endothelial cluster identified by unbiased differential expression analysis.

**Extended Data Fig. 11:**
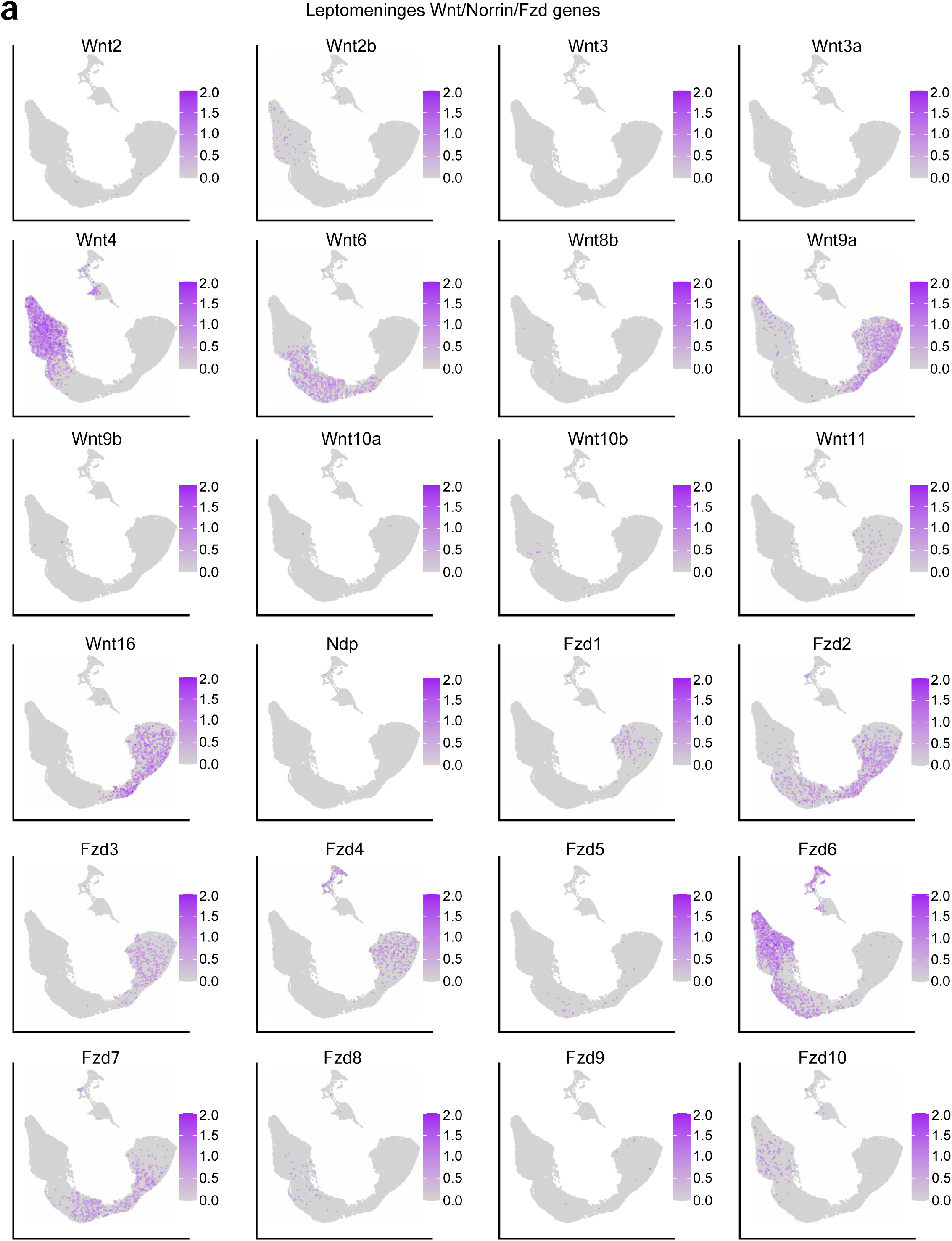
Comprehensive survey of Wnt ligands, Frizzled receptors, and Norrin signaling components in the leptomeninges. (a) Feature plots showing expression of Wnt ligands, Ndp (Norrin), and Frizzled receptor family members (Fzd1–Fzd10) across leptomeningeal cell populations from WT uninfected mice. Wnt4 and Wnt5a were enriched in fibroblast populations, whereas Wnt5b, Wnt9a, and Wnt16 were associated with arachnoid barrier-related populations. Multiple Frizzled receptors were broadly distributed among fibroblast and barrier cell populations. Canonical BBB-associated ligands Wnt7a and Wnt7b were largely absent from leptomeningeal populations. Expression values are shown on a standardized scale (0–2 normalized expression units) to facilitate comparison across genes.

**Extended Data Fig. 12:**
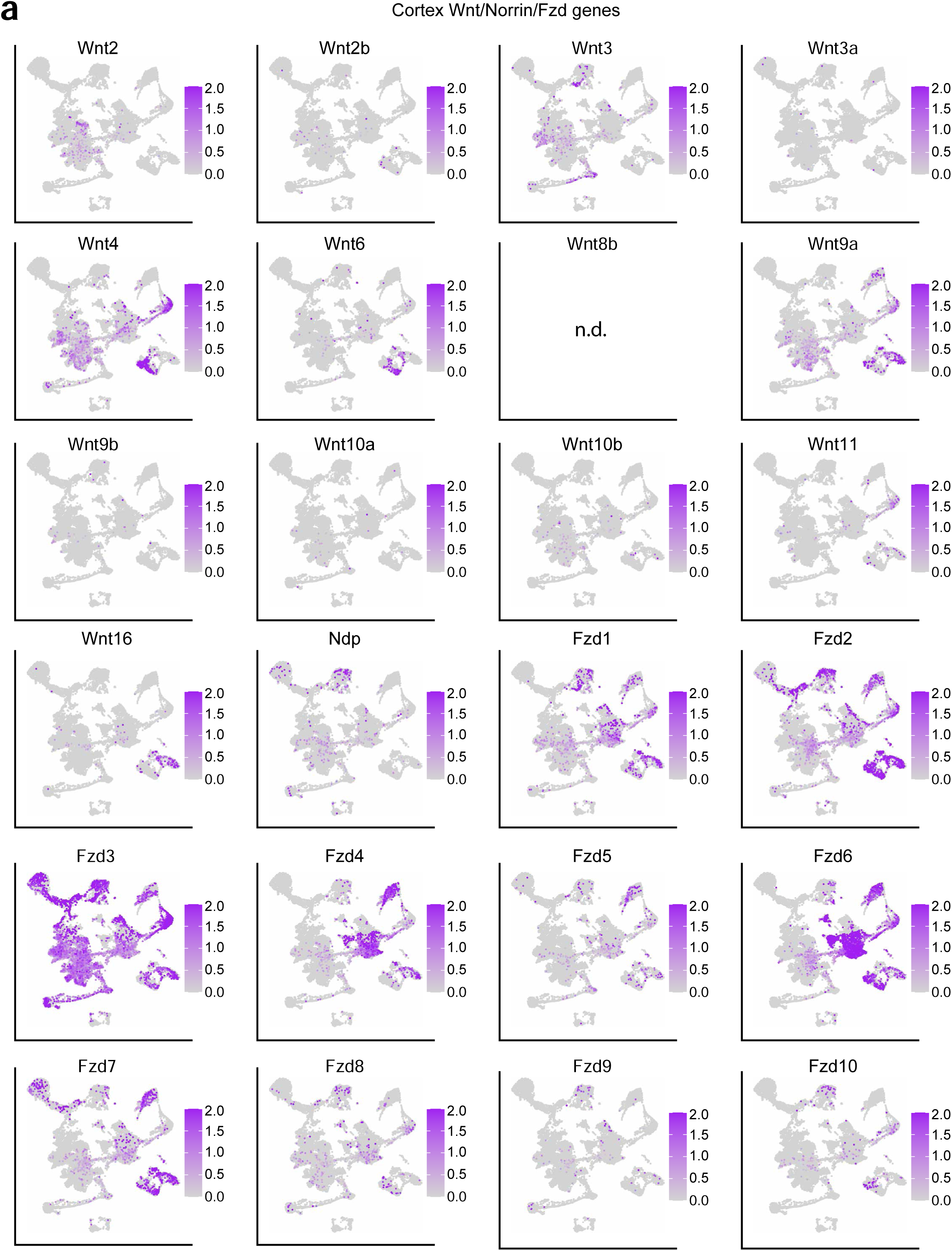
Comprehensive survey of Wnt ligands, Frizzled receptors, and Norrin signaling components in the cortex. (a) Feature plots showing expression of Wnt ligands, Ndp (Norrin), and Frizzled receptor family members (Fzd1–Fzd10) across cortical cell populations from control samples. Wnt7a and Wnt7b were detected across cortical parenchymal populations, consistent with the established cortical BBB Wnt ligand environment. Several additional Wnt ligands, including Wnt4, Wnt5a, Wnt5b, Wnt6, Wnt9a, and Wnt11, were also detected in subsets of cortical cell populations. Frizzled receptors were broadly expressed across cortical cell classes, suggesting widespread competence for Wnt pathway responsiveness. Expression values are shown on a standardized scale (0–2 normalized expression units) to facilitate comparison across genes. n.d., not detected.

**Extended Data Fig. 13:**
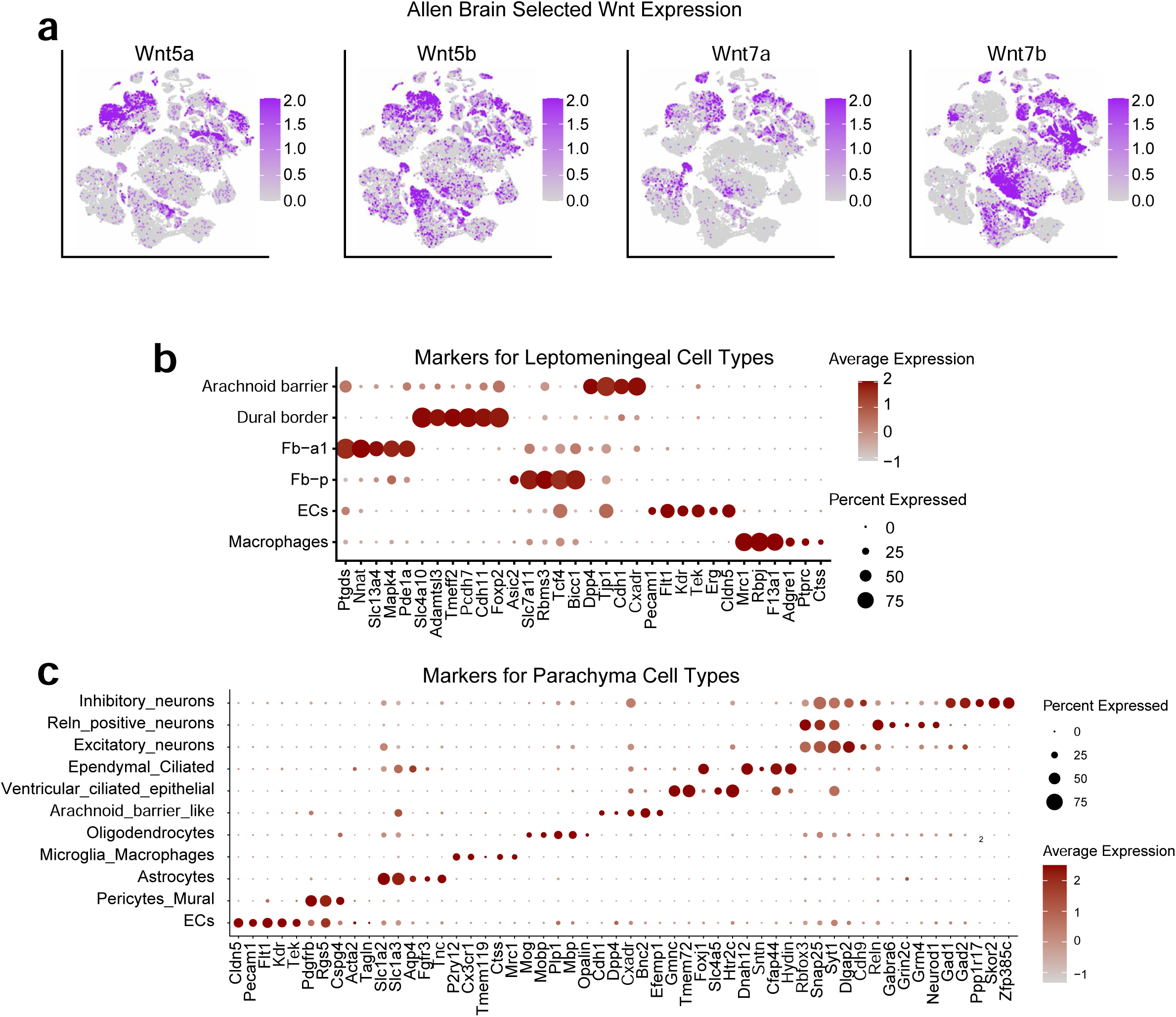
Validation of cell-type annotations and comparison of Wnt pathway components across leptomeningeal, cortical, and Allen Brain datasets. (a) Independent validation using the Allen Brain Cell Atlas. Feature plots show expression of Wnt5a, Wnt5b, Wnt7a, and Wnt7b across major CNS cell classes. Wnt7a and Wnt7b expression was enriched in parenchymal neural populations, whereas Wnt5-family ligands were broadly distributed across multiple CNS cell populations, supporting distinct Wnt signaling environments between meningeal and cortical compartments. (b) Dot plot demonstrating expression of established marker genes used to identify leptomeningeal cell populations, including endothelial cells, macrophages, pia fibroblasts, arachnoid fibroblasts, dural border cells, and arachnoid barrier cells. (c) Dot plot demonstrating expression of established marker genes used to identify cortical cell populations, including endothelial cells, mural cells, astrocytes, oligodendrocytes, microglia/macrophages, ciliated epithelial cells, and neuronal populations.

**Extended Data Fig. 14:**
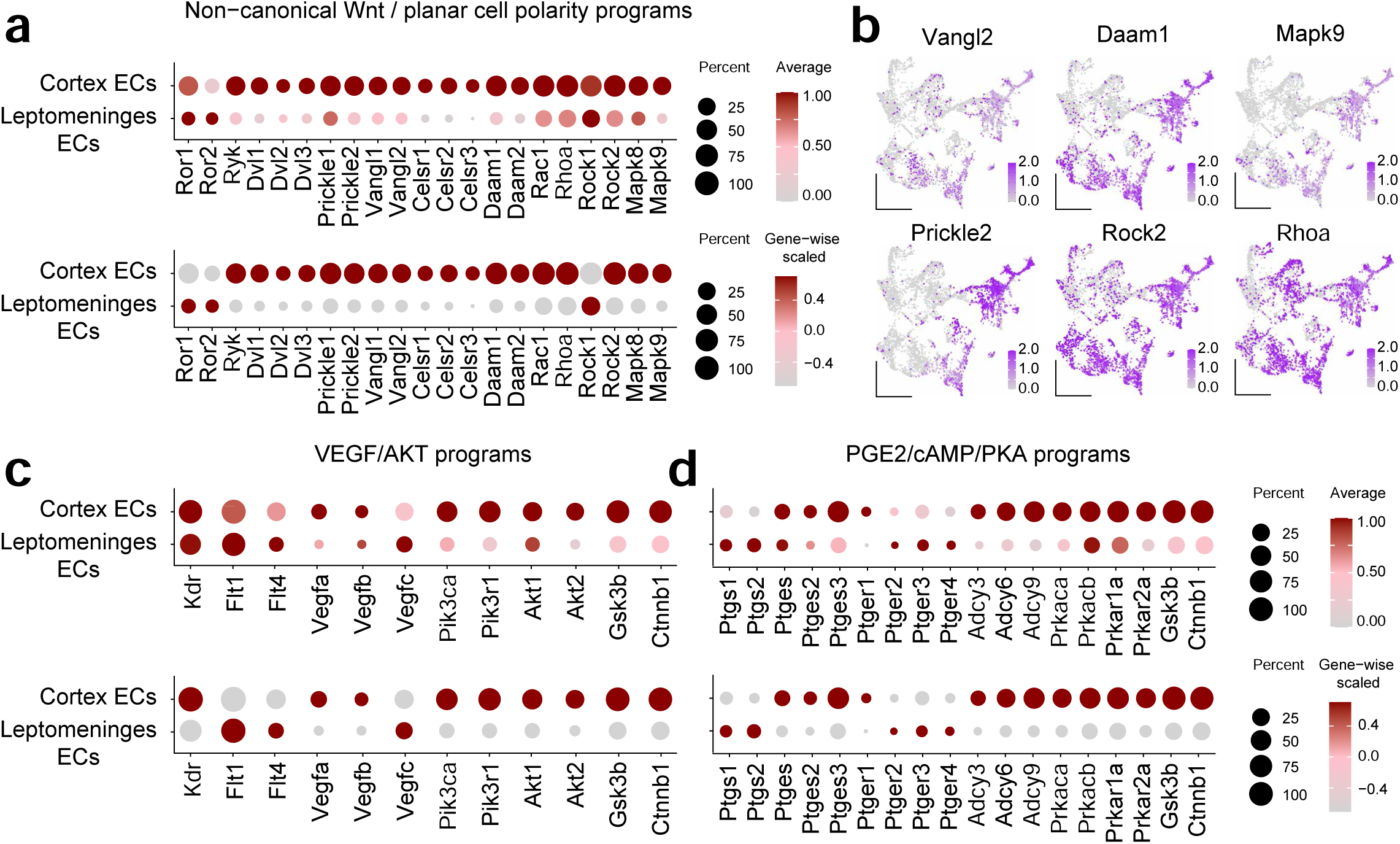
Survey of alternative β-catenin regulatory and non-canonical Wnt signaling pathways in cortical and leptomeningeal ECs. (a) Dot plots showing expression of non-canonical Wnt/planar cell polarity (PCP) signaling genes in cortical and leptomeningeal endothelial cells (ECs). Upper panel shows log1p-transformed average expression; lower panel shows gene-wise scaled expression. Dot size indicates the percentage of expressing cells and color indicates expression level. (b) Representative UMAP feature plots of selected PCP pathway genes. Vangl2, Prickle2, Daam1, and Mapk9 exhibited enrichment within cortical EC populations, whereas Rhoa and Rock2 were broadly expressed across both endothelial compartments. Feature plots are displayed on a common expression scale (0–2 normalized expression units). (c) Dot plots showing expression of genes associated with VEGF/PI3K/AKT signaling. Upper panel shows log1p-transformed average expression; lower panel shows gene-wise scaled expression. (d) Dot plots showing expression of genes associated with prostaglandin E2 (PGE2)/cAMP/PKA signaling. Upper panel shows log1p-transformed average expression; lower panel shows gene-wise scaled expression.

**Extended Data Fig. 15:**
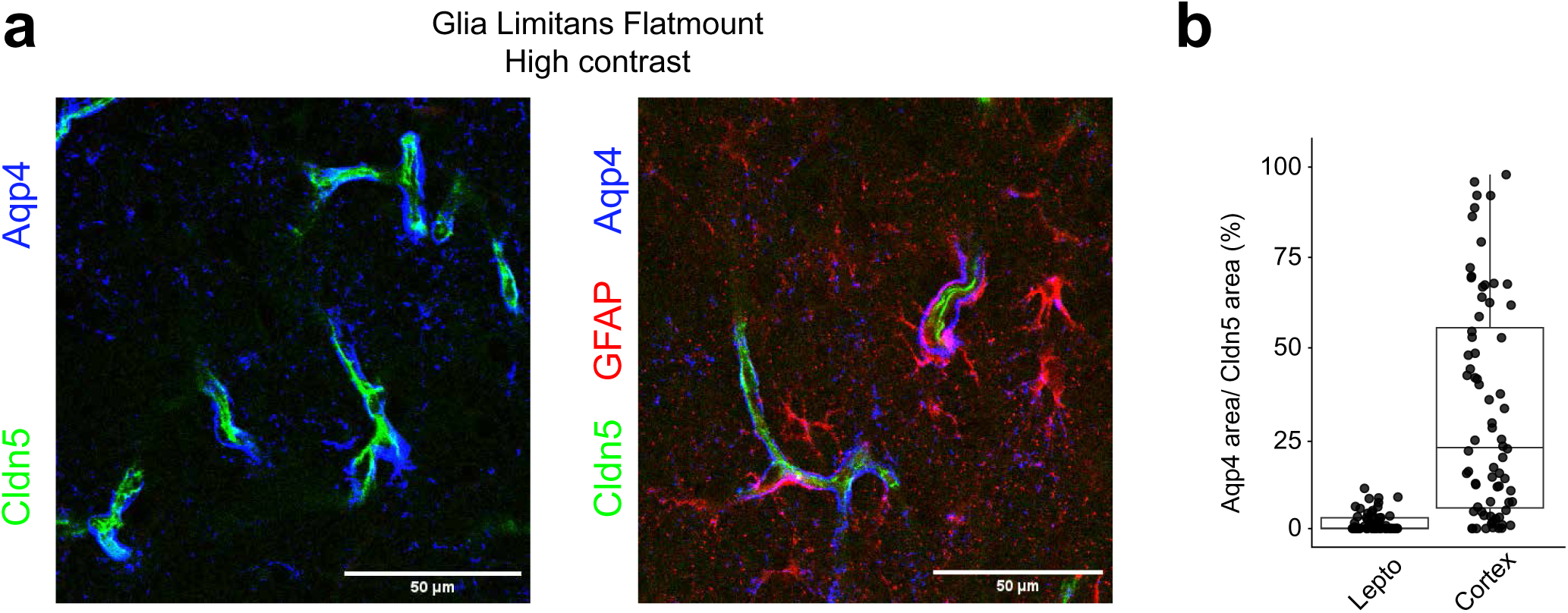
Aqp4 localizes to distinct compartment-specific structures across the leptomeningeal surface and cortical glial limitans. (a) Contrast-enhanced flatmount images of the cortical glial limitans stained for Cldn5 (green), Aqp4 (blue), and GFAP (red). Left panel shows Aqp4 and Cldn5 channels demonstrating that Aqp4-positive structures are spatially associated with cortical vessels. Right panel includes the GFAP channel, revealing that vascular-associated Aqp4 signal colocalizes with GFAP-positive astrocytic endfeet surrounding cortical vessels. (b) Quantification of Aqp4 overlap normalized to total Cldn5-positive vascular area within subarachnoid space (SAS) and cortical compartments. Aqp4 overlap with Cldn5-positive vessels was minimal in the SAS but significantly increased in cortical vessels, with some cortical ROIs demonstrating 80–90% overlap. Each point represents an individual ROI. Statistical comparisons were performed using a two-sided Wilcoxon rank-sum test following outlier removal using the 1.5× interquartile range method. Scale bars, 50 µm.

**Extended Data Fig. 16:**
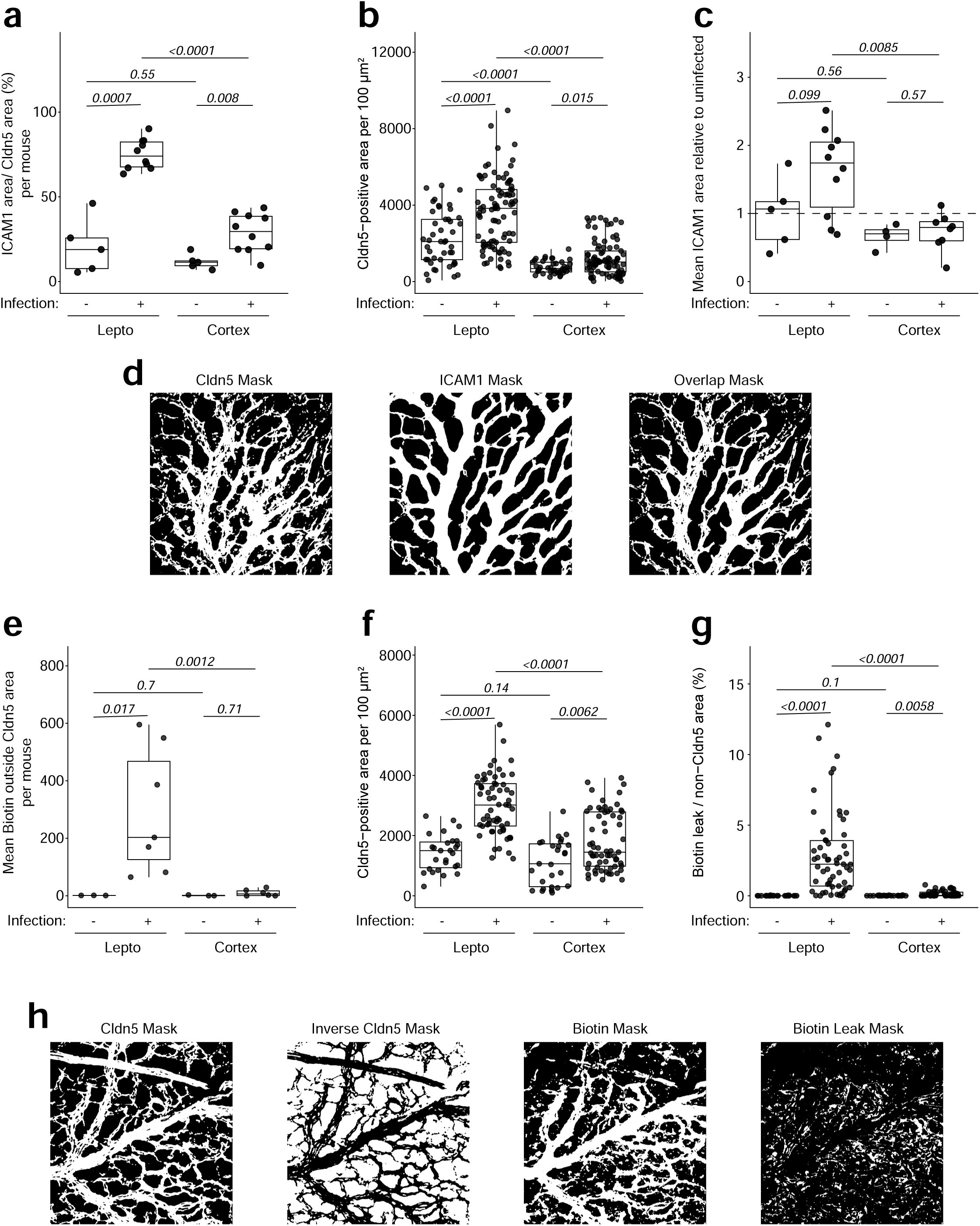
Quantification workflow and compartment-specific inflammatory and permeability responses across leptomeningeal and cortical vascular interfaces. (a) Mouse-level quantification of ICAM1-positive vascular coverage. ROI-level measurements were averaged for each mouse prior to statistical analysis. Each dot represents one mouse. (b) Absolute Cldn5-positive vessel area quantified per 100 × 100 µm ROI across leptomeningeal and cortical compartments in uninfected and infected mice. (c) Relative ICAM1 vascular coverage normalized to the mean uninfected value within each anatomical compartment (leptomeninges or cortex). Dashed line indicates the compartment-specific uninfected baseline. These analyses demonstrate that differences in ICAM1 vascular coverage are not solely explained by changes in total Cldn5-positive vessel area. (d) Representative binary masks used for ICAM1 overlap analysis, including thresholded Cldn5 mask, ICAM1 mask, and overlap mask generated using logical AND operations. (e) Mouse-level quantification of extravascular biotin leakage. ROI-level measurements were averaged for each mouse prior to statistical analysis. Each dot represents one mouse. (f) Total Cldn5-positive vessel area quantified per 100 × 100 µm ROI in biotin tracer experiments. (g) Normalized extravascular biotin leakage quantified as biotin-positive area outside the Cldn5 mask divided by total non-Cldn5 area within each ROI. These analyses demonstrate that elevated leptomeningeal tracer leakage is not solely attributable to differences in vessel density or available extravascular tissue area. (h) Representative binary masks used for biotin leak analysis, including thresholded Cldn5 mask, inverse Cldn5 mask defining non-vascular tissue space, biotin mask, and final biotin leak mask identifying biotin-positive signal outside the vascular compartment.

**Extended Data Fig. 17:**
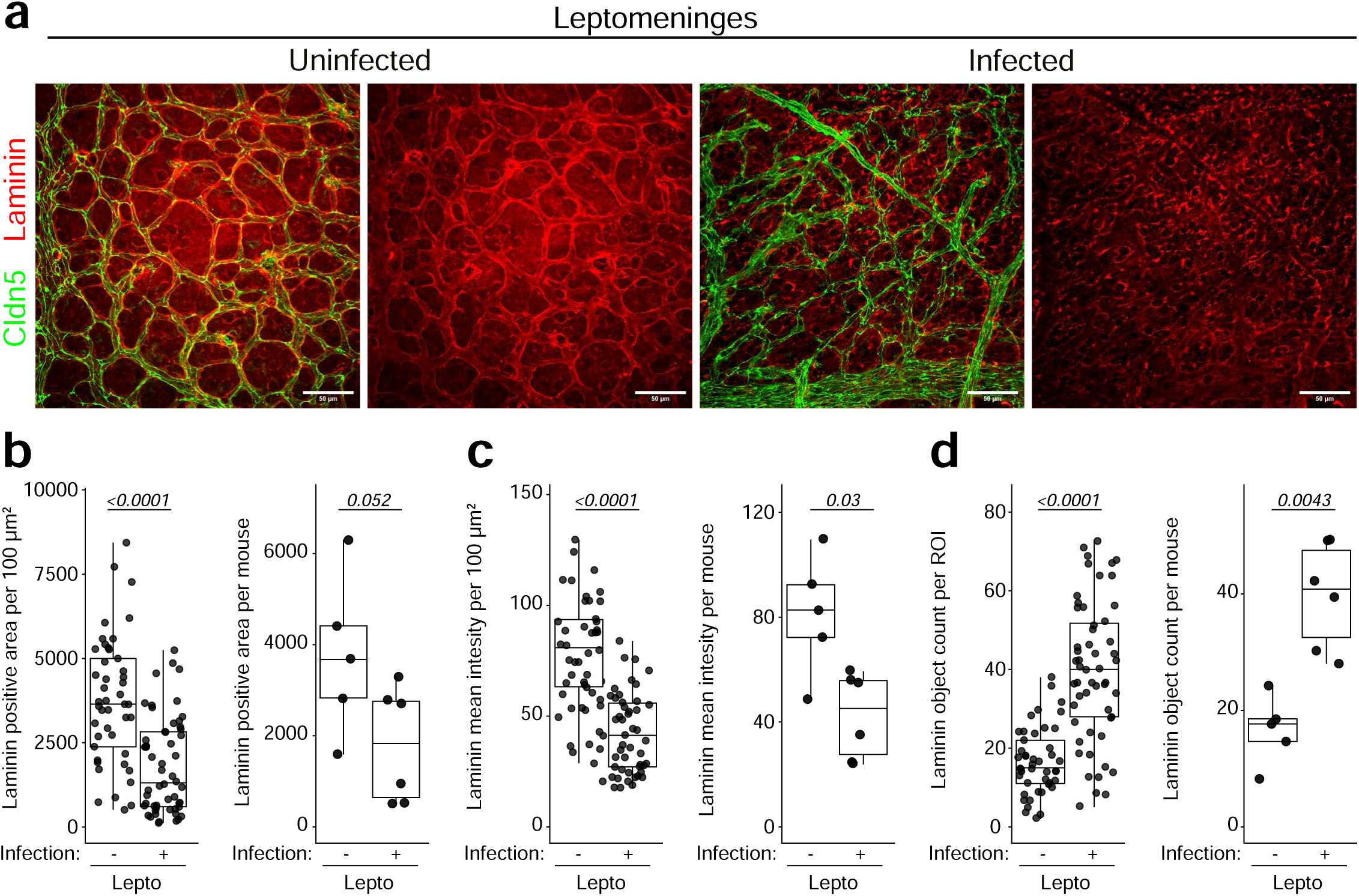
Infection induces loss and fragmentation of laminin-positive vascular structures in the leptomeninges. (a) Representative maximum-intensity projections of leptomeningeal vessels from uninfected and *E. coli*– infected neonatal mice stained for Claudin-5 (Cldn5; green) and Laminin (red). Upper panels show merged images and lower panels show the Laminin channel alone. Infection induced marked disruption and fragmentation of laminin-positive vascular structures within the leptomeningeal compartment. Scale bars, 50 µm. (b) ROI-level and mouse-level quantification of laminin-positive area. Laminin-positive area was quantified from thresholded laminin masks within fixed 100 × 100 µm ROIs. Mouse-level analyses were generated by averaging ROI-level measurements for each animal prior to statistical analysis. (c) ROI-level and mouse-level quantification of mean laminin intensity. Mean laminin intensity was quantified within fixed 100 × 100 µm ROIs across leptomeningeal vascular regions. (d) ROI-level and mouse-level quantification of laminin object count. Thresholded laminin-positive objects were identified using particle analysis in Fiji/ImageJ, and the number of contiguous laminin-positive objects within each ROI was quantified as a measure of laminin fragmentation and structural discontinuity. Each dot represents one ROI or one mouse, as indicated. Boxplots show median and interquartile range. Statistical comparisons were performed using two-sided Wilcoxon rank-sum tests.

**Extended Data Fig. 18:**
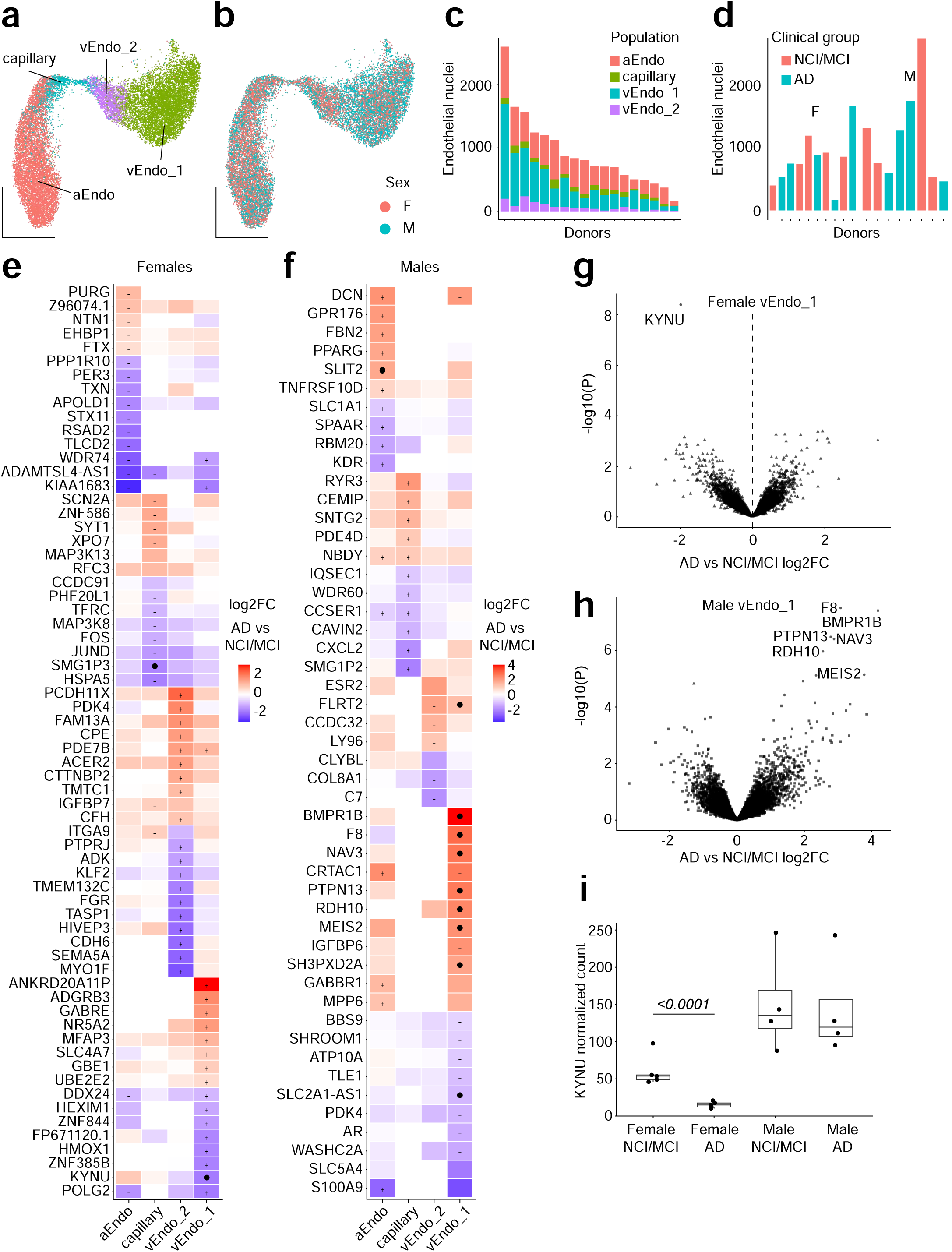
Human leptomeningeal endothelial cells exhibit disease-associated transcriptional remodeling in Alzheimer’s disease. (a) UMAP of human leptomeningeal endothelial nuclei showing the four endothelial populations identified in the reanalysis: arterial endothelial (aEndo), capillary, vEndo_1, and vEndo_2 cells. (b) UMAP of the same endothelial nuclei colored by donor sex. Female (F) and male (M) nuclei were distributed throughout the endothelial manifold without overt segregation by sex. (c) Endothelial nuclei contributed by individual donors, stratified by endothelial population. Each bar represents one donor and colors indicate endothelial subtype. (d) Endothelial nuclei contributed by individual donors according to clinical group and sex. Each bar represents one donor and colors indicate NCI/MCI or Alzheimer’s disease (AD) clinical classification. Female (F) and male (M) donors are indicated above the corresponding groups. (e–f) Heatmaps showing sex-stratified transcriptional changes associated with clinical AD across aEndo, capillary, vEndo_2, and vEndo_1 populations in female (e) and male (f) donors. Color indicates the donor-level pseudobulk DESeq2 log2 fold change for AD relative to NCI/MCI, with red indicating increased and blue indicating decreased expression in AD. Genes represent the union of genes meeting the indicated differential-expression thresholds across endothelial populations. Plus signs (+) denote genes with nominal DESeq2 Wald-test *P* < 0.01, and filled black circles denote genes significant following Benjamini–Hochberg multiple-testing correction (FDR < 0.05). Differential-expression analyses were performed independently within each sex using donor-level pseudobulk counts. (g–h) Volcano plots showing donor-level pseudobulk differential expression between AD and NCI/MCI donors within the vEndo_1 population in females (g) and males (h). The x-axis indicates DESeq2 log2 fold change for AD relative to NCI/MCI and the y-axis indicates −log10(*P*) from the DESeq2 Wald test. Selected genes exhibiting prominent disease-associated changes are labeled. KYNU was strongly reduced in female AD vEndo_1 endothelial cells, whereas male vEndo_1 cells exhibited a distinct transcriptional response including increased expression of *F8*, *BMPR1B*, *PTPN13*, *NAV3*, *RDH10*, and *MEIS2*. (i) Donor-level DESeq2-normalized KYNU counts in vEndo_1 endothelial cells stratified by sex and clinical group. Each point represents one donor; boxplots show the median and interquartile range. KYNU expression was significantly reduced in female AD compared with female NCI/MCI donors (log2FC = −1.98, DESeq2 Wald-test *P* = 3.95 × 10⁻⁹, Benjamini–Hochberg FDR = 2.8 × 10⁻⁵), whereas no corresponding AD-associated difference was detected in males (log2FC = −0.06, *P* = 0.892, FDR = 1.0).

## References

1 Daneman, R. & Prat, A. The blood-brain barrier. Cold Spring Harb Perspect Biol 7, a020412 (2015). 10.1101/cshperspect.a020412

2 Saunders, N. R., Dziegielewska, K. M., Fame, R. M., Lehtinen, M. K. & Liddelow, S. A. The choroid plexus: a missing link in our understanding of brain development and function. Physiol Rev 103, 919–956 (2023). 10.1152/physrev.00060.2021

3 O’Leary, F. & Campbell, M. The blood-retina barrier in health and disease. FEBS J 290, 878–891 (2023). 10.1111/febs.16330

4 Smyth, L. C. D. & Kipnis, J. Redefining CNS immune privilege. Nat Rev Immunol 25, 766–775 (2025). 10.1038/s41577-025-01175-0

5 Barnett, S. N., et al. An organotypic atlas of human vascular cells. Nat Med 30, 3468–3481 (2024). 10.1038/s41591-024-03376-x

6 Vanlandewijck, M. et al. A molecular atlas of cell types and zonation in the brain vasculature. Nature 554, 475-480 (2018). 10.1038/nature25739

7 Pfau, S. J., et al. Characteristics of blood-brain barrier heterogeneity between brain regions revealed by profiling vascular and perivascular cells. Nat Neurosci 27, 1892–1903 (2024). 10.1038/s41593-024-01743-y

8 Dani, N., et al. A cellular and spatial map of the choroid plexus across brain ventricles and ages. Cell 184, 3056–3074 e3021 (2021). 10.1016/j.cell.2021.04.003

9 Wang, Y., et al. Norrin/Frizzled4 signaling in retinal vascular development and blood brain barrier plasticity. Cell 151, 1332–1344 (2012). 10.1016/j.cell.2012.10.042

10 Wang, J., Rattner, A. & Nathans, J. Bacterial meningitis in the early postnatal mouse studied at single-cell resolution. Elife 12 (2023). https://doi.org/ARTN e86130, 10.7554/eLife.86130

11 Mapunda, J. A., et al. VE-cadherin in arachnoid and pia mater cells serves as a suitable landmark for in vivo imaging of CNS immune surveillance and inflammation. Nat Commun 14, 5837 (2023). 10.1038/s41467-023-41580-4

12 Pietila, R., et al. Molecular anatomy of adult mouse leptomeninges. Neuron 111, 3745–3764 e3747 (2023). 10.1016/j.neuron.2023.09.002

13 Kearns, N. A., et al. Dissecting the human leptomeninges at single-cell resolution. Nat Commun 14, 7036 (2023). 10.1038/s41467-023-42825-y

14 Li, Y., et al. Decoding the spatiotemporal development of human meninges. Cell (2026). 10.1016/j.cell.2026.04.040

15 Smyth, L. C. D. et al. Identification of direct connections between the dura and the brain. Nature 627, 165-173 (2024). 10.1038/s41586-023-06993-7

16 Iadecola, C. The Neurovascular Unit Coming of Age: A Journey through Neurovascular Coupling in Health and Disease. Neuron **96**, 17–42 (2017). 10.1016/j.neuron.2017.07.030

17 Armulik, A., et al. Pericytes regulate the blood-brain barrier. Nature 468, 557–561 (2010). 10.1038/nature09522

18 Abbott, N. J., Ronnback, L. & Hansson, E. Astrocyte-endothelial interactions at the blood-brain barrier. Nat Rev Neurosci 7, 41–53 (2006). 10.1038/nrn1824

19 Yao, Y., Chen, Z. L., Norris, E. H. & Strickland, S. Astrocytic laminin regulates pericyte differentiation and maintains blood brain barrier integrity. Nat Commun 5, 3413 (2014). 10.1038/ncomms4413

20 Engelhardt, B. & Ransohoff, R. M. Capture, crawl, cross: the T cell code to breach the blood-brain barriers. Trends Immunol 33, 579–589 (2012). 10.1016/j.it.2012.07.004

21 Obermeier, B., Daneman, R. & Ransohoff, R. M. Development, maintenance and disruption of the blood-brain barrier. Nat Med 19, 1584-1596 (2013). 10.1038/nm.3407

22 Zhou, Y., et al. Canonical WNT signaling components in vascular development and barrier formation. Journal of Clinical Investigation 124, 3825–3846 (2014). 10.1172/JCI76431

23 Cho, C., Smallwood, P. M. & Nathans, J. Reck and Gpr124 Are Essential Receptor Cofactors for Wnt7a/Wnt7b-Specific Signaling in Mammalian CNS Angiogenesis and Blood–Brain Barrier Regulation. Neuron 95, 1056–1073.e1055 (2017). 10.1016/j.neuron.2017.08.004

24 Daneman, R., et al. Wnt/beta-catenin signaling is required for CNS, but not non-CNS, angiogenesis. Proc Natl Acad Sci U S A 106, 641–646 (2009). 10.1073/pnas.0805165106

25 Vanhollebeke, B., et al. Tip cell-specific requirement for an atypical Gpr124– and Reck-dependent Wnt/β-catenin pathway during brain angiogenesis. eLife 4, e06489 (2015). 10.7554/eLife.06489

26 Junge, H. J., et al. TSPAN12 regulates retinal vascular development by promoting Norrin-but not Wnt-induced FZD4/beta-catenin signaling. Cell 139, 299–311 (2009). 10.1016/j.cell.2009.07.048

27 Montagne, A., et al. Blood–brain barrier breakdown in the aging human hippocampus. Neuron 85, 296-302 (2015). 10.1016/j.neuron.2014.12.032

28 Shlosberg, D., Benifla, M., Kaufer, D. & Friedman, A. Blood-Brain Barrier Breakdown as a Therapeutic Target in Traumatic Brain Injury. Nature Reviews Neurology 6, 393–403 (2010). 10.1038/nrneurol.2010.74

29 Sweeney, M. D., Sagare, A. P. & Zlokovic, B. V. Blood–brain barrier breakdown in Alzheimer disease and other neurodegenerative disorders. Nature Reviews Neurology 14, 133–150 (2018). 10.1038/nrneurol.2017.188

30 Yang, A. C. et al. A human brain vascular atlas reveals diverse mediators of Alzheimer’s risk. Nature 603, 885-892 (2022). 10.1038/s41586-021-04369-3

31 Sun, N., et al. Single-nucleus multiregion transcriptomic analysis of brain vasculature in Alzheimer’s disease. Nat Neurosci 26, 970–982 (2023). 10.1038/s41593-023-01334-3

32 Kivisakk, P., et al. Human cerebrospinal fluid central memory CD4+ T cells: evidence for trafficking through choroid plexus and meninges via P-selectin. Proc Natl Acad Sci U S A 100, 8389–8394 (2003). 10.1073/pnas.1433000100

33 Bartholomaus, I., et al. Effector T cell interactions with meningeal vascular structures in nascent autoimmune CNS lesions. Nature 462, 94–98 (2009). 10.1038/nature08478

34 Kivisakk, P., et al. Localizing central nervous system immune surveillance: meningeal antigen-presenting cells activate T cells during experimental autoimmune encephalomyelitis. Ann Neurol 65, 457–469 (2009). 10.1002/ana.21379

35 Hobson, R., et al. Clonal CD8(+) T cells populate the leptomeninges and coordinate with immune cells in human degenerative brain diseases. Nat Immunol 27, 323–335 (2026). 10.1038/s41590-025-02401-6

36 Ye, X., et al. Norrin, frizzled-4, and Lrp5 signaling in endothelial cells controls a genetic program for retinal vascularization. Cell 139, 285–298 (2009). 10.1016/j.cell.2009.07.047

37 Wang, Y., et al. Interplay of the Norrin and Wnt7a/Wnt7b signaling systems in blood-brain barrier and blood-retina barrier development and maintenance. Proc Natl Acad Sci U S A 115, E11827–E11836 (2018). 10.1073/pnas.1813217115

38 Masckauchan, T. N., et al. Wnt5a signaling induces proliferation and survival of endothelial cells in vitro and expression of MMP-1 and Tie-2. Mol Biol Cell 17, 5163–5172 (2006). 10.1091/mbc.e06-04-0320

39 Skaria, T., Bachli, E. & Schoedon, G. Wnt5A/Ryk signaling critically affects barrier function in human vascular endothelial cells. Cell Adh Migr 11, 24–38 (2017). 10.1080/19336918.2016.1178449

40 Davis, G. E. & Senger, D. R. Endothelial extracellular matrix: biosynthesis, remodeling, and functions during vascular morphogenesis and neovessel stabilization. Circ Res 97, 1093–1107 (2005). 10.1161/01.RES.0000191547.64391.e3

41 Stratman, A. N., Malotte, K. M., Mahan, R. D., Davis, M. J. & Davis, G. E. Pericyte recruitment during vasculogenic tube assembly stimulates endothelial basement membrane matrix formation. Blood 114, 5091–5101 (2009). 10.1182/blood-2009-05-222364

42 Gautam, J., Zhang, X. & Yao, Y. The role of pericytic laminin in blood brain barrier integrity maintenance. Sci Rep 6, 36450 (2016). 10.1038/srep36450

43 Thomsen, M. S., Birkelund, S., Burkhart, A., Stensballe, A. & Moos, T. Synthesis and deposition of basement membrane proteins by primary brain capillary endothelial cells in a murine model of the blood-brain barrier. J Neurochem 140, 741–754 (2017). 10.1111/jnc.13747

44 Fassbender, K., et al. Endothelial-derived adhesion molecules in bacterial meningitis: association to cytokine release and intrathecal leukocyte-recruitment. J Neuroimmunol 74, 130–134 (1997). 10.1016/s0165-5728(96)00214-7

45 Liang, Y., et al. Kynurenine Pathway Metabolites as Biomarkers in Alzheimer’s Disease. Dis Markers 2022, 9484217 (2022). 10.1155/2022/9484217

46 Giil, L. M., et al. Kynurenine Pathway Metabolites in Alzheimer’s Disease. J Alzheimers Dis 60, 495–504 (2017). 10.3233/JAD-170485

47 Walker, D. G., Link, J., Lue, L. F., Dalsing-Hernandez, J. E. & Boyes, B. E. Gene expression changes by amyloid beta peptide-stimulated human postmortem brain microglia identify activation of multiple inflammatory processes. J Leukoc Biol 79, 596–610 (2006). 10.1189/jlb.0705377

48 Wang, M., et al. Immune-proteo-metabolomic changes link to Abeta and tau pathology in Alzheimer disease. Alzheimers Dement 22, e71359 (2026). 10.1002/alz.71359

49 Mahi, N. A., Najafabadi, M. F., Pilarczyk, M., Kouril, M. & Medvedovic, M. GREIN: An Interactive Web Platform for Re-analyzing GEO RNA-seq Data. Sci Rep 9, 7580 (2019). 10.1038/s41598-019-43935-8

