## Supplementary material for "Defining a leptomeningeal blood–cerebrospinal fluid barrier as a specialized vascular interface": GSE_Annotation_Table

| GSE_ID | n_samples | cell_types | ages | PMID | Hyperlink | Journal | Year | Title |
|---|---|---|---|---|---|---|---|---|
| GSE136848 | 41 | Brain Endothelial; Heart Endothelial; Lung Endothelial | Adult | 31944177 | https://www.ncbi.nlm.nih.gov/pubmed/31944177 | Elife | 2020 | Endothelial heterogeneity across distinct vascular beds during homeostasis and inflammation |
| GSE163821 | 20 | Brain Endothelial; Heart Endothelial; Kidney Endothelial | Adult; Old | 34097533 | https://europepmc.org/article/MED/34097533 | Physiological Genomics | 2021 | Age−dependent transcriptional alterations in cardiac endothelial cells |
| GSE155797 | 16 | Liver Endothelial | Adult | 34038707 | https://www.ncbi.nlm.nih.gov/pubmed/34038707 | Dev Cell | 2021 | A spatial vascular transcriptomic, proteomic, and phosphoproteomic atlas unveils an angiocrine Tie−Wnt signaling axis in the liver |
| GSE123021 | 12 | Microglia | Adult | 30606613 | https://www.ncbi.nlm.nih.gov/pubmed/30606613 | Neuron | 2019 | Developmental Heterogeneity of Microglia and Brain Myeloid Cells Revealed by Deep Single−Cell RNA Sequencing |
| GSE127758 | 12 | Brain Endothelial | Old; Young | 31086348 | https://pmc.ncbi.nlm.nih.gov/articles/PMC6642642/ | Nat Med | 2019 | Aged blood impairs hippocampal neural precursor activity and activates microglia via brain endothelial cell VCAM1 |
| GSE111839 | 10 | Brain Endothelial; Kidney Endothelial; Liver Endothelial; Lung Endothelial; Whole Brain | Young | 31913116 | https://www.ncbi.nlm.nih.gov/pubmed/31913116 | Elife | 2020 | A genome−wide view of the de−differentiation of central nervous system endothelial cells in culture |
| GSE134005 | 10 | bMECs | Adult; Young | 31426861 | https://www.ncbi.nlm.nih.gov/pubmed/31426861 | Acta Neuropathol Commun | 2019 | Transcriptome clarifies mechanisms of lesion genesis versus progression in models of Ccm3 cerebral cavernous malformations |
| GSE104530 | 9 | Heart Endothelial | Old; Young | 29212850 | https://www.ncbi.nlm.nih.gov/pubmed/29212850 | Physiol Genomics | 2018 | Endothelial transcriptomics reveals activation of fibrosis−related pathways in hypertension |
| GSE104701 | 9 | Bone Marrow Endothelial | Adult | 29934585 | https://www.ncbi.nlm.nih.gov/pubmed/29934585 | Nat Commun | 2018 | Stem cell factor is selectively secreted by arterial endothelial cells in bone marrow |
| GSE102562 | 8 | Microglia | Adult | 28930663 | https://www.ncbi.nlm.nih.gov/pubmed/28930663 | Immunity | 2017 | The TREM2−APOE Pathway Drives the Transcriptional Phenotype of Dysfunctional Microglia in Neurodegenerative Diseases |
| GSE181508 | 8 | Lung Endothelial | Old; Young | 35879310 | https://www.ncbi.nlm.nih.gov/pubmed/35879310 | Nat Commun | 2022 | Dysfunctional ERG signaling drives pulmonary vascular aging and persistent fibrosis |
| GSE95201 | 8 | Liver Endothelial; Lung Endothelial | Adult | 31611708 | https://www.ncbi.nlm.nih.gov/pubmed/31611708 | Nat Neurosci | 2019 | Profiling the mouse brain endothelial transcriptome in health and disease models reveals a core blood−brain barrier dysfunction module |
| GSE124868 | 7 | Microglia | Adult; Young | 30846482 | https://www.ncbi.nlm.nih.gov/pubmed/30846482 | J Exp Med | 2019 | Impaired ?V?8 and TGF? signaling lead to microglial dysmaturation and neuromotor dysfunction |
| GSE52564 | 7 | Brain Endothelial; Microglia; Whole Brain | Adult | 25186741 | https://www.ncbi.nlm.nih.gov/pubmed/25186741 | J Neurosci | 2014 | An RNA−sequencing transcriptome and splicing database of glia, neurons, and vascular cells of the cerebral cortex |
| GSE102563 | 6 | Microglia | Adult | 28930663 | https://www.ncbi.nlm.nih.gov/pubmed/28930663 | Immunity | 2017 | The TREM2−APOE Pathway Drives the Transcriptional Phenotype of Dysfunctional Microglia in Neurodegenerative Diseases |
| GSE117083 | 6 | Pericytes | Young | 31249304 | https://www.ncbi.nlm.nih.gov/pubmed/31249304 | Nat Commun | 2019 | Loss of the transcription factor RBPJ induces disease−promoting properties in brain pericytes |
| GSE135442 | 6 | Kidney Endothelial | Adult | 32651395 | https://www.ncbi.nlm.nih.gov/pubmed/32651395 | Sci Rep | 2020 | Glomerular endothelial cell heterogeneity in Alport syndrome |
| GSE227145 | 6 | Bone Marrow Derived ECs | Old; Young | 37037837 | https://www.ncbi.nlm.nih.gov/pubmed/37037837 | Nat Commun | 2023 | Restoring bone marrow niche function rejuvenates aged hematopoietic stem cells by reactivating the DNA Damage Response |
| GSE75668 | 6 | Brain Endothelial; Pericytes | Young | 27725773 | https://www.ncbi.nlm.nih.gov/pubmed/27725773 | Sci Rep | 2016 | Analysis of the brain mural cell transcriptome |
| GSE180169 | 5 | Lung Endothelial | Adult | 34528097 | https://www.ncbi.nlm.nih.gov/pubmed/34528097 | Cardiovasc Res | 2022 | Single−cell RNA sequencing profiling of mouse endothelial cells in response to pulmonary arterial hypertension |
| GSE74052 | 5 | Brain Endothelial | Adult | 28288111 | https://pubmed.ncbi.nlm.nih.gov/28288111/ | Nat Med | 2017 | Gpr124 is essential for blood−brain barrier integrity in central nervous system disease |
| GSE106692 | 4 | Microglia | Adult | 29268096 | https://www.ncbi.nlm.nih.gov/pubmed/29268096 | Neuron | 2017 | Activation of the STING−Dependent Type I Interferon Response Reduces Microglial Reactivity and Neuroinflammation |
| GSE115188 | 4 | Whole Brain | Adult | 29912970 | https://www.ncbi.nlm.nih.gov/pubmed/29912970 | PLoS One | 2018 | The influence of adolescent nicotine exposure on ethanol intake and brain gene expression |
| GSE122952 | 4 | Brain Endothelial | Adult | 32529267 | https://www.ncbi.nlm.nih.gov/pubmed/32529267 | Acta Neuropathol | 2020 | HIF−1? is involved in blood−brain barrier dysfunction and paracellular migration of bacteria in pneumococcal meningitis |
| GSE142361 | 4 | Microglia | Old; Young | 32065074 | https://www.ncbi.nlm.nih.gov/pubmed/32065074 | J Cereb Blood Flow Metab | 2020 | Transcriptomic and functional studies reveal undermined chemotactic and angiostimulatory properties of aged microglia during stroke recovery |
| GSE163561 | 4 | bMECs | Adult | 34903569 | https://www.ncbi.nlm.nih.gov/pubmed/34903569 | J Neurosci | 2022 | Endothelial Sphingosine−1−Phosphate Receptor 4 Regulates Blood−Brain Barrier Permeability and Promotes a Homeostatic Endothelial Phenotype |
| GSE227147 | 4 | Bone Marrow Derived ECs | Old | 37037837 | https://www.ncbi.nlm.nih.gov/pubmed/37037837 | Nat Commun | 2023 | Restoring bone marrow niche function rejuvenates aged hematopoietic stem cells by reactivating the DNA Damage Response |
| GSE268823 | 4 | bMECs | Adult | 38957986 | https://www.ncbi.nlm.nih.gov/pubmed/38957986 | Arterioscler Thromb Vasc Biol | 2024 | Cav?3 Contributes to the Maintenance of the Blood−Brain Barrier and Alleviates Symptoms of Experimental Autoimmune Encephalomyelitis |
| GSE72341 | 4 | Pericytes | Adult; Young | 26634440 | https://www.ncbi.nlm.nih.gov/pubmed/26634440 | Science | 2016 | Fetal liver hematopoietic stem cell niches associate with portal vessels |
| GSE126127 | 3 | Whole Brain | Young | 33460120 | https://www.ncbi.nlm.nih.gov/pubmed/33460120 | J Neurochem | 2021 | Loss of PRMT1 in the central nervous system (CNS) induces reactive astrocytes and microglia during postnatal brain development |
| GSE149776 | 3 | Bone Marrow Endothelial | Adult | 34848712 | https://www.ncbi.nlm.nih.gov/pubmed/34848712 | Nat Commun | 2021 | Neuropilin 1 regulates bone marrow vascular regeneration and hematopoietic reconstitution |
| GSE159754 | 3 | Kidney Endothelial | Adult | 34301760 | https://www.ncbi.nlm.nih.gov/pubmed/34301760 | Cancer Res | 2021 | Endothelial Reprogramming Stimulated by Oncostatin M Promotes Inflammation and Tumorigenesis in VHL−Deficient Kidney Tissue |
| GSE169267 | 3 | Bone Marrow Endothelial | Adult | 38344689 | https://www.ncbi.nlm.nih.gov/pubmed/38344689 | Nat Cardiovasc Res | 2023 | Bone marrow adipocytes fuel emergency hematopoiesis after myocardial infarction |
| GSE172360 | 3 | Liver Endothelial | Adult | 35364013 | https://pubmed.ncbi.nlm.nih.gov/35364013/ | Cell Stem Cell | 2022 | Specification of fetal liver endothelial progenitors to functional zonated adult sinusoids requires c−Maf induction |
| GSE173311 | 3 | Brain Endothelial | Adult | 36170290 | https://pubmed.ncbi.nlm.nih.gov/36170290/ | PLoS One | 2022 | Optimized protocol for translatome analysis of mouse brain endothelial cells |
| GSE193544 | 3 | Kidney Endothelial | Adult | 35440634 | https://www.ncbi.nlm.nih.gov/pubmed/35440634 | Nat Commun | 2022 | Loss of vascular endothelial notch signaling promotes spontaneous formation of tertiary lymphoid structures |
| GSE223871 | 3 | Liver Endothelial | Young | 38125029 | https://www.ncbi.nlm.nih.gov/pubmed/38125029 | iScience | 2023 | Quantitative proteomics and RNA−sequencing of mouse liver endothelial cells identify novel regulators of BMP6 by iron |
| GSE233052 | 3 | Liver Endothelial | Young | 37563497 | https://www.ncbi.nlm.nih.gov/pubmed/37563497 | Angiogenesis | 2023 | Aging impairs the ability of vascular endothelial stem cells to generate endothelial cells in mice. |
| GSE61636 | 3 | Bone Marrow Endothelial | Adult | 26441307 | https://www.ncbi.nlm.nih.gov/pubmed/26441307 | Stem Cell Reports | 2015 | Vascular Platform to Define Hematopoietic Stem Cell Factors and Enhance Regenerative Hematopoiesis |
| GSE64510 | 3 | Pericytes | Young | 26019175 | https://www.ncbi.nlm.nih.gov/pubmed/26019175 | Genes Dev | 2015 | PDGFR? signaling drives adipose tissue fibrosis by targeting progenitor cell plasticity |
| GSE84492 | 3 | Whole Brain | Adult | NA | NA | NA | 2017 | RNA−seq analysis of Kdm5c deficient brains in mice |
| GSE85657 | 3 | bMECs | Adult | 28970240 | https://www.ncbi.nlm.nih.gov/pubmed/28970240 | J Exp Med | 2017 | Thrombospondin1 (TSP1) replacement prevents cerebral cavernous malformations |
| GSE90459 | 3 | Brain Endothelial | Adult | NA | NA | NA | 2017 | RNA−seq analysis of mouse brain endothelial cells and glioma endothelial cells |
| GSE147357 | 2 | Pericytes | Young | 34096078 | https://www.ncbi.nlm.nih.gov/pubmed/34096078 | Immunol Rev | 2021 | The fibroblast: An emerging key player in thymic T cell selection |
| GSE229292 | 2 | Kidney Endothelial | Adult | 37667913 | https://www.ncbi.nlm.nih.gov/pubmed/37667913 | J Cell Sci | 2023 | Shock drives a STAT3 and JunB−mediated coordinated transcriptional and DNA methylation response in the endothelium |
| GSE230022 | 2 | Lung Endothelial | Adult; Young | 38098255 | https://www.ncbi.nlm.nih.gov/pubmed/38098255 | Aging Cell | 2024 | Viral uptake and pathophysiology of the lung endothelial cells in age−associated severe SARS−CoV−2 infection models |
